# Arabidopsis NIM1-INTERACTING1 (NIMIN1) is a multi-domain protein controlling transition from systemic acquired resistance (SAR) to cell death

**DOI:** 10.1101/2024.07.22.604541

**Authors:** Mathias Saur, Kristin Steilner, Ashir Masroor, Philipp Hubel, Artur J.P. Pfitzner, Ursula M. Pfitzner

**Author notes:** Author for correspondence: Ursula Pfitzner, Email:. Department of Plant Pathology, University of Agriculture Faisalabad, Sub-Campus Burewala-Vehari, Pakistan (Ashir Masroor).

## Abstract

- NIM1-INTERACTING (NIMIN) proteins were identified by interaction with the systemic acquired resistance (SAR) key regulator NPR1. The Arabidopsis family comprises four related, yet distinct members. Ample evidence predicts that NIMIN1 is a multi-domain protein which comes into action after the onset of salicylic acid (SA) signaling prior to induction of the *PR-1* defense gene.
- Bioinformatics, protein-protein interaction assays and expression studies in tobacco were applied to explore functions of NIMIN1.
- The N-terminal sequence encompassing amino acids 1 to 15 conveyed accumulation of NIMIN1 protein *in planta*. This newly identified segment and the known conserved NIMIN1 domains, i.e., two distinct NPR1 binding sites and an EAR motif, are neatly separated from each other by disordered regions. Co-expression of *NIMIN1* and *NPR1* reinforced accumulation of NPR1 protein while ectopic overexpression of *NIMIN1* promoted emergence of cell death driven by the EAR motif and disturbed development of tobacco plants.
- We suggest that NIMIN1 acts as a dynamic signaling agent controlling transition of pathogen-infected leaves from survival to tissue collapse. Initially, NIMIN1 binding renders the NPR1 transcription complex sensitive to the SAR signal SA enabling *PR-1* transcription. At high levels of SA, NIMIN1 is outcompeted by SA from the NPR1 C-terminus, and accumulating NIMIN1 engages via its EAR motif with TOPLESS-RELATED3, thereby affecting global hormone signaling and inducing cell death in severely endangered tissue.

## Introduction

Plants have evolved different mechanisms to defend themselves against invading microorganisms. Pathogens adapted to their hosts are able to evade or counteract the first line of active defense by secretion of effector proteins which manipulate plant cellular processes and interfere with the immune response (Wang *et al*., 2022a). As an evolutionary response, plants have acquired resistance proteins that sense the presence of microbial effectors to induce effector-triggered immunity (ETI; Jones & Dangl, 2006). ETI is a strong defense response directed against all kinds of biotrophic pathogens and present in all higher plants. It is typically associated with localized cell death emerging in small zones around pathogen infection sites. The phenotype of local necrotic lesions is specified as hypersensitive response (HR; Balint-Kurti, 2019). HR restricts further spread of pathogens within the plant, and non-infected parts of the plant become protected against secondary attacks, a phenomenon known as systemic acquired resistance (SAR; Ross, 1961; Fu & Dong, 2013). SAR is not associated with cell death, but with cell survival (Zavaliev *et al*., 2020). Induction of both HR and SAR coincides with increases in the defense hormone salicylic acid (SA; Malamy *et al*., 1990; Métraux *et al*., 1990) which serves as a local signal for coordinate induction of the heterogeneous group of pathogenesis-related (PR) proteins (Ward *et al*., 1991; Vernooij *et al*., 1994). PR proteins display different antimicrobial activities and, all in all, are believed to mediate plant resistance (van Loon *et al*., 2006). While HR exhibits high induction of SA and PR proteins, SAR is characterized by more moderate hormone levels and defense gene activation (Malamy *et al*., 1990; Ward *et al*., 1991).

The SA signal is transduced through the SAR key regulatory protein NON-EXPRESSOR OF PR GENES1 (NPR1; aka NON-INDUCIBLE IMMUNITY1, NIM1) which is essential for SA-mediated expression of the SAR marker gene *PR-1* (Cao *et al*., 1994; Delaney *et al*., 1995; Cao *et al*., 1997; Ryals *et al*., 1997; Shah *et al*., 1997). In Arabidopsis, NPR1 is part of a small protein family. Four members, NPR1 to NPR4, share domain architecture including two highly conserved stretches in their C-termini (Maier *et al*., 2011). Recent evidence showed that NPR1 to NPR4 function as *bona fide* SA receptors (Canet *et al*., 2010a; Maier *et al*., 2011; Fu *et al*., 2012; Wu *et al*., 2012; Manohar *et al*., 2015; Castelló *et al*., 2018; Ding *et al*., 2018; Neeley *et al*., 2019; Wang *et al*., 2020; Konopka *et al*., 2022). All NPR1 to NPR4 perceive the SA molecule through the same basal mechanism engaging the arginine residue (R) in the conserved LENRV-like motif for binding the carboxyl of SA (Konopka *et al*., 2022). Nevertheless, NPR1 to NPR4 display unique biochemical properties to enable diverse signaling outputs (Castelló *et al*., 2018; Ding *et al*., 2018; Konopka *et al*., 2022). Defense gene activation via NPR1 is mediated by members of the TGA transcription factor family (Zhang *et al*., 2003; Powers *et al*., 2024). TGA factors bind, on the one side, to SA-responsive *cis*-regulatory elements present in *PR-1* promoters (Strompen *et al*., 1998; Lebel *et al*., 1998; Johnson *et al*., 2003) and, on the other side, to the NPR1 protein, itself (Zhang *et al*., 1999; Zhou *et al*., 2000; Kumar *et al*., 2022), thereby integrating the SA receptor and co-activator NPR1 in a transcription complex positioned on the *PR-1* gene. In addition to TGA factors, NPR1 binds NIM1-INTERACTING (NIMIN, N) proteins.

In Arabidopsis (At), there are four *NIMIN* genes, *N1*, *N1b*, *N2* and *N3*, encoding small structurally related, yet distinct, proteins that localize to the nucleus (Weigel *et al*., 2001). Tobacco (Nt) harbors multiple *NIMIN2*-type genes (Zwicker *et al*., 2007). Arabidopsis *NIMIN1* and *N2* and tobacco *N2*-type genes are themselves induced by SA thus corroborating their significance for SAR (Horvath *et al*., 1998; Weigel *et al*., 2001; Glocova *et al*., 2005; Zwicker *et al*., 2007; Fonseca *et al*., 2010; Hermann *et al*., 2013). Interaction of NIMIN proteins with AtNPR1 is independent from TGA factor binding (Weigel *et al*., 2001, 2005). While TGA factors bind to the central ankyrin repeat region of AtNPR1 (Zhang *et al*., 1999; Zhou *et al*., 2000; Kumar *et al*., 2022), NIMIN1 and N2 share a conserved domain by which they bind to a region in the C-terminus of NPR1 termed N1/N2 binding domain (N1/N2BD) (Weigel *et al*., 2001; Maier *et al*., 2011). Of note, in yeast, interaction of NIMIN1 and N2 proteins with NPR1 is compromised in presence of SA (Maier *et al*., 2011). From these findings and from genetic evidence (Canet *et al*., 2010a), it was deduced that the region originally defined as N1/N2BD may participate not only in NIMIN interaction, but also in perception of the SA signal. Indeed, NtNPR1, AtNPR3 and AtNPR4 were shown to directly sense SA via the essential arginine in their LENRV(-like) motifs and the N1/N2BD homologous regions (Neeley *et al*., 2019; Wang *et al*., 2020; Konopka *et al*., 2022). Hormone binding induces a conformational switch which, in turn, activates the NtNPR1 and AtNPR4 receptors. In contrast to NtNPR1 and AtNPR4, Arabidopsis NPR1 is, however, not highly susceptible to the SA molecule on its own (Fu *et al*., 2012; Konopka *et al*., 2022; Kumar *et al*., 2022), and it was suggested that NIMIN1/N2 interaction is vital to sensitize AtNPR1 to SA (Konopka *et al*., 2022). On the other hand, constitutive overexpression of *NIMIN1* in Arabidopsis and *Nicotiana benthamiana* led to strong suppression of SA and pathogen-induced *PR-1* gene expression (Weigel *et al*., 2005; Hermann *et al*., 2013). Together, the data imply that, in Arabidopsis, NIMIN1 and N2 control signal transduction through the SA sensitive C-terminal region of NPR1.

Here, we further characterize regulatory properties of NIMIN1. Altogether, NIMIN1 harbors four distinct domains that mediate differential interaction with NPR1, interaction with transcriptional co-repressors of the TOPLESS family and accumulation of NIMIN1 protein. The domains are clearly separated from each other by intermediate spacer regions of disorder thus allowing NIMIN1 to follow different paths in the course of the plant defense response.

## Materials and Methods

### Plant growth, virus infection and chemical treatment of plant tissue

Tobacco (*Nicotiana tabacum* cv. Samsun NN; *Nicotiana benthamiana*) plants were grown in commercial potting soil under natural light conditions in a greenhouse. Infection of leaves with *Tobacco mosaic virus* (TMV) was as described previously (Grüner & Pfitzner, 1994).

For chemical treatment, leaf disks were cut from young *N. tabacum* plants and from *N. benthamiana* plants 4 or 5 days after agroinfiltration and floated for 2 to 4 days on water or solutions of 1 mM SA or 0.3 mM benzothiadiazole (BTH).

### Yeast two-hybrid analyses

Yeast two-hybrid (Y2H) assays were conducted as reported earlier (Weigel *et al*., 2001; Maier *et al*., 2011; Neeley *et al*., 2019). Yeast cells were grown in absence or presence of 0.3 mM SA as indicated. *LacZ* reporter gene activities were tested in duplicate with at least three independent colonies. Experiments were repeated at least once with new yeast transformants. Representative results collected in one experimental set are shown. Results are depicted as mean enzyme activities in Miller units, plus and minus standard deviation (SD).

### Generation of transgenic plants

Agrobacterium-mediated transformation of tobacco with pBin19 and TurboID-based gene constructs was performed as described (Grüner & Pfitzner, 1994). Several independent primary transformants were selfed, and progeny plants of the T1, T2 and T3 generations were analyzed as indicated. Transgenic seedlings were selected on solid ½ MS medium with antibiotic. Proteins were extracted with GUS lysis buffer from two to four leaf disks for each plant and from pools of ten whole seedlings for each transgenic line.

### 3,3’-Diaminobenzidine (DAB) staining

DAB staining was according to Thordal-Christensen *et al*. (1997). Agroinfiltrated *N. benthamiana* leaves were cut from the stem and photographed. Then, leaves were infiltrated with 1 mg/ml DAB in 20 mM MES buffer (pH 6.5) in a needleless 60 ml syringe by applying gentle vacuum. After 4 hours of incubation, leaves were destained with a 4:1 solution of ethanol:acetone until chlorophyll was completely depleted. Leaves were photographed under white light.

### Bioinformatic analyses

Disorder profiles were generated with DISOPRED3 on the PSIPRED Workbench (bioinf.cs.ucl.ac.uk; Jones & Cozzetto, 2015).

## Results

### Domain structure of NIMIN1

Using the NPR1 bait, we isolated clones for *NIMIN1*, *N2* and *N3* in a single yeast two-hybrid screen from an Arabidopsis cDNA library. Sequence analysis revealed that the encoded proteins share several distinct amino acid motifs (Fig. S1; Weigel *et al*., 2001). All comprise prolonged stretches of acidic amino acids marked orange in Figure S1. Furthermore, NIMIN1 and N2 harbor canonical nuclear localization signals (marked yellow). The sequence motif marked blue occurring in both NIMIN1 and N2 is an NPR1 interaction motif (Weigel *et al*., 2001). Via this motif, NIMIN1 and N2 bind to the conserved N1/N2BD regions in the C-termini of NPR1 proteins from Arabidopsis and tobacco (Maier *et al*., 2011; Hermann *et al*., 2013; Neeley *et al*., 2019).

In addition to the blue-labeled NPR1-binding site, NIMIN1 harbors still another domain for interaction with NPR1, here denoted EDF motif (marked green in Fig. S1). This motif was first recognized in the rice NIMIN homolog NEGATIVE REGULATOR OF RESISTANCE (NRR; Chern *et al*., 2005, 2012). NRR binds to rice NH1 (NPR1 HOMOLOG 1) predominantly via a sequence related to the green-labeled NIMIN1 motif and to Arabidopsis NPR1 predominantly via a sequence similar to the blue-labeled motif in NIMIN1. The EDF motif is shared by NIMIN1 and N3 (Fig. S1). Unlike NIMIN1, binding of NIMIN3 to NPR1 is not affected by SA (Fig. S2; Maier *et al*., 2011). To assess the roles of the two NPR1 interaction motifs in NIMIN1, we tested interaction of mutants with Arabidopsis NPR1 in Y2H assays. All NIMIN mutants used in this work are depicted in Table S1. While interaction of NIMIN3 with NPR1 was abolished by a double mutation in the EDF motif, interaction of the corresponding NIMIN1 mutant was just reduced, and binding of the mutant protein to NPR1 was still fully responsive to SA (Fig. S2). On the other hand, a double mutation in the blue NPR1 interaction motif nullified binding of NIMIN1 to NPR1 (Fig. S2; Weigel *et al*., 2001) underscoring the notion that interaction with NPR1 through the blue-labeled motif is basic to SA-controlled NIMIN1 activity *in planta*.

A third domain which was originally identified in all NIMIN1 to N3 and termed LxL motif (Weigel *et al*., 2001) is an ethylene-responsive element binding factor–associated amphiphilic repression (EAR) motif of the type LxLxL (Ohta *et al*., 2001; Kagale & Rozwadowski, 2011). The EAR motif occurs at the C-terminal ends of NIMIN proteins (marked red in Fig. S1). The EAR motif constitutes a repression domain that recruits the co-repressor TOPLESS (TPL) and related proteins (Causier *et al*., 2012; Dinesh *et al*., 2016; Plant *et al*., 2021). We were not able to show Y2H interaction of full-length TPL with NIMIN1 or N3 which contain nearly identical EAR motifs (L_1_DL_2_NL_3_AL_4_ in N1; L_1_DL_2_NL_3_SL_4_ in N3). However, both bound to the N-terminal TPL(1-333) fragment encompassing the EAR-interacting TPL domain (TPD, aa 1-204; Fig. S3a; Ke *et al*., 2015; Martin-Arevalillo *et al*., 2017). NIMIN2, on the other hand, was not bound (data not shown). NIMIN3 exhibited an approximately equal interaction strength with TPL(1-333) and NPR1, while NIMIN1 binding to NPR1 was clearly preferred over binding to TPL(1-333) (Fig. S3b). Deletion of three of the four leucine residues in the EAR motif (L_2_ to L_4;_ NIMINΔEAR) and mutation of the two core leucine residues L_2_ and L_3_ to alanine abrogated interactions of NIMIN1 and N3 with TPL(1-333), but not with NPR1 (Fig. S3b). On the contrary, mutants of NIMIN1 in the two NPR1 interaction motifs (blue and green-labeled domains) were still able to bind to TPL(1-333), but not to NPR1 (data not shown). Although NIMIN proteins can interact simultaneously with NPR1 and TGA factors (Weigel *et al*., 2001; Hermann *et al*., 2013), we were not able to demonstrate ternary complex formation of NIMINs with NPR1 and TPL(1-333) (BD–NPR1/NIMIN+AD–TPL(1-333); data not shown).

### The 5’-end of the *NIMIN1* coding region harbors a structural motif to reinforce protein accumulation *in planta*

The conserved amino acid motifs originally identified by us through alignment of NIMIN1 to N3 fall together with deep valleys in the NIMIN1 disorder profile (Fig. 1a) supporting the view that NIMIN1 is a multi-domain protein, and that the identified sequence motifs are vital for NIMIN1 function. The structural motifs are neatly separated from each other by about 20-amino acid long highly disordered regions designated by clusters of polar amino acids, mostly glutamic acid. The disorder profile also revealed another potential domain at the immediate N-terminus of NIMIN1, likewise isolated from the blue NPR1-binding motif by a stretch of disorder (Fig. 1a). Sequences homologous to the NIMIN1 N-terminus are not present in N2 or N3 nor in the highly similar N1b protein (Fig. 1b; Weigel *et al*., 2001).

**Fig. 1.**
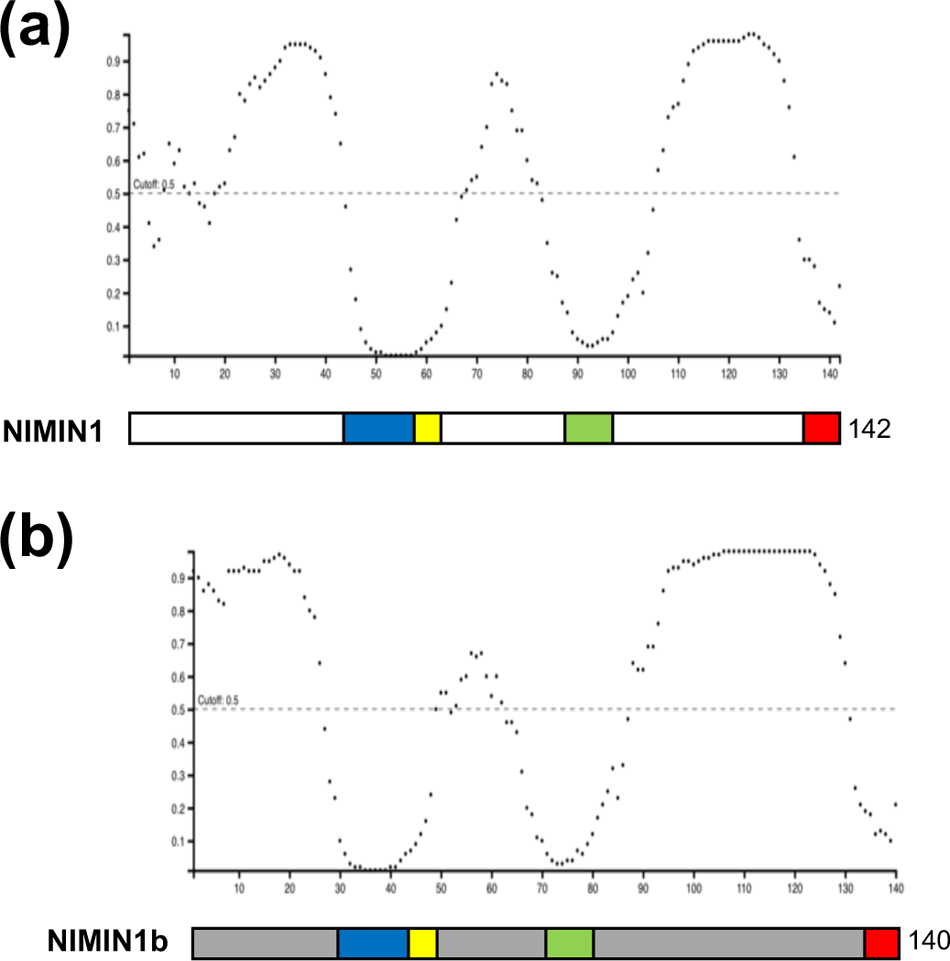
Domain structures and disorder profiles of Arabidopsis NIMIN1 and NIMIN1b. The disorder profiles were generated with DISOPRED3 on the PSIPRED Workbench (Jones & Cozzetto, 2015). The diagrams are drawn approximately to scale. Colored boxes denote conserved structural motifs. Nuclear localization signal (NLS, yellow); interaction motif mediating N1 binding to the SA sensitive C-terminus of NPR1 (blue); a second NPR1 interaction motif not linked to the SA response (EDF motif, green); EAR motif (red). (a) Domain structure and disorder profile of NIMIN1. (b) Domain structure and disorder profile of NIMIN1b.

To test a potential function of the newly identified segment, we fused the sequence encoding the 15 N-terminal amino acids of *NIMIN1* (*nTN1*) to the *N1b* cDNA. Figure S4 shows the domain structure and disorder profile of the encoded nTN1–N1b fusion protein. Initially, we tested interaction of the chimeric protein in Y2H. NIMIN1b, like N1, bound strongly to NPR1, and interaction was sensitive to SA (Fig. S5a). Yet, NIMIN1b did not bind TPL(1-333), although N1b and N1 harbor identical EAR motifs (Fig. S5b and c). The chimeric nTN1–N1b protein exhibited similar interaction with NPR1 as NIMIN1b (Fig. S5b). Quite surprisingly, however, nTN1–N1b, unlike NIMIN1b, was able to bind to TPL(1-333), and interaction depended on the EAR motif (Fig. S5b and d).

In a next step, we analyzed the nTN1–N1b chimeric protein *in planta*. To facilitate protein detection, the *nTN1–N1b* cDNA was fused to the sequence for VENUS, a variant of yellow fluorescent protein (YFP; Nagai *et al*., 2002). Similar constructs were generated for *N1* and *N1b* (Fig. 2a). All genes were expressed transiently under control of the *CaMV 35S* promoter in *N. benthamiana*, and accumulation of fusion proteins was analyzed in crude leaf extracts by immunodetection using either a polyclonal antiserum or a monoclonal antibody (mAb) against green fluorescent protein (GFP). For direct comparison of protein accumulation from different gene constructs, Agrobacterium strains were infiltrated in the left (L) and right (R) halves, respectively, of the same leaves. While N1b–VENUS fusion protein was at most faintly detectable in leaf extracts, nTN1–N1b–VENUS and N1–VENUS accumulated to similar levels (Fig. 2b). Most isolated protein appeared intact, but we also observed degradation products with the fusion proteins. On the other hand, N-terminal truncation of NIMIN1, yielding N1(16-142)–VENUS, prevented traceable accumulation of the mutant full-length protein (Fig. S6a).

**Fig. 2.**
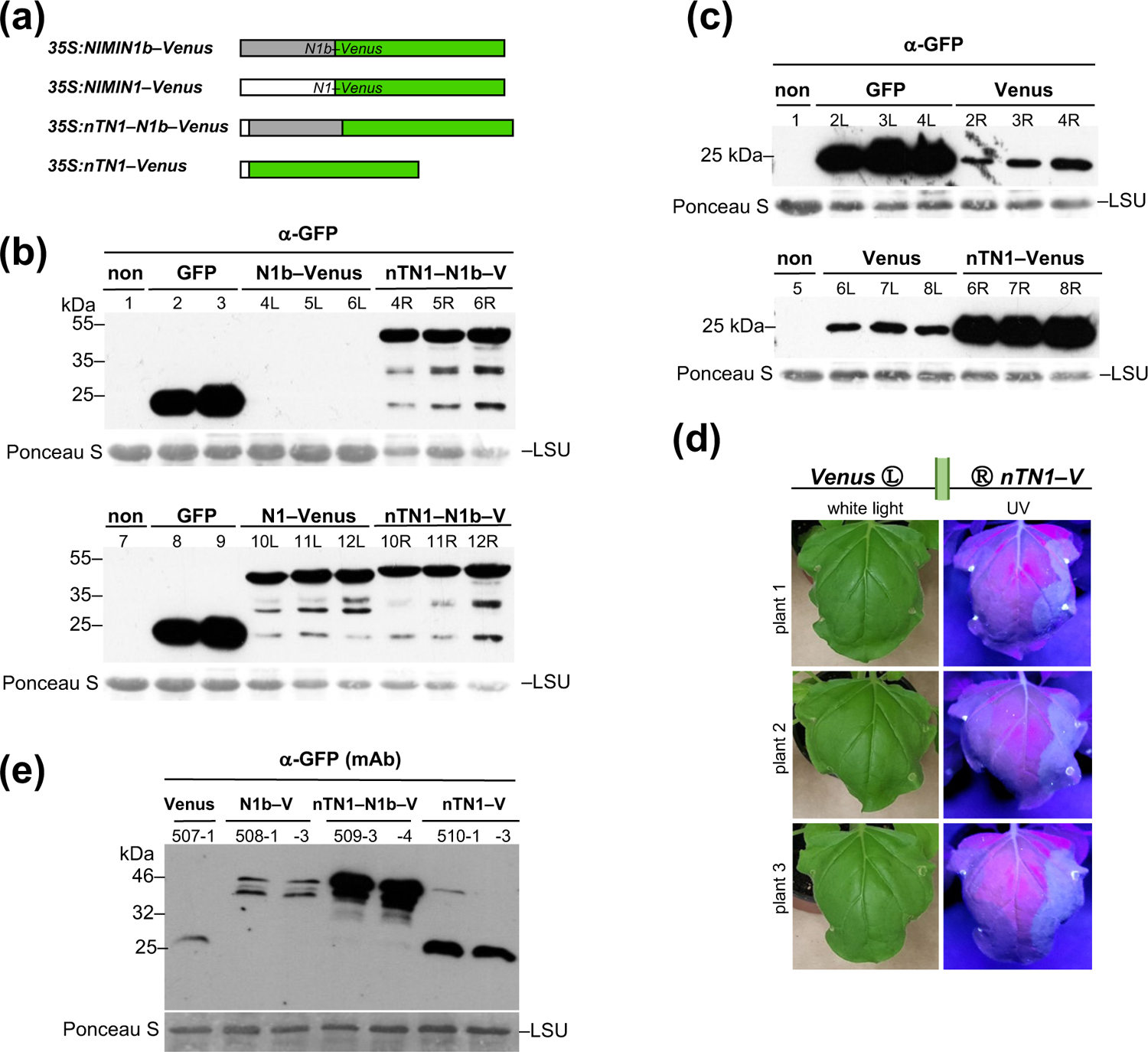
The N-terminal *NIMIN1* sequence *(nTN1)* reinforces protein accumulation *in planta*. (a) Schematic representation of *VENUS* fusion constructs. The diagram is drawn approximately to scale. (b) Accumulation of nTN1–N1b–VENUS in *Nicotiana benthamiana*. Accumulation of nTN1–N1b– VENUS is compared to accumulation of N1b–VENUS and N1–VENUS. For direct comparison of protein accumulation from different constructs, Agrobacterium strains were infiltrated in the left (L) and right (R) halves of the same leaves as indicated. Three plants were infiltrated in parallel. Immunodetection of VENUS fusion proteins in leaf extracts was with a polyclonal antiserum directed against GFP. Staining of the large subunit of RUBISCO (LSU) with Ponceau S demonstrates loading of the nitrocellulose filter. (c) Accumulation of nTN1–VENUS in *Nicotiana benthamiana* visualized by immunodetection. Accumulation of nTN1–VENUS is compared to accumulation of VENUS. (d) Accumulation of nTN1–VENUS in *Nicotiana benthamiana* visualized by fluorescence. Accumulation of nTN1–VENUS is compared to accumulation of VENUS. Photographs of leaves were taken under white light and under UV light as indicated. (e) Accumulation of nTN1–N1b–VENUS and nTN1– VENUS in transgenic tobacco seedlings. Accumulation of nTN1 fusion proteins is compared to accumulation of N1b–VENUS and VENUS. Results for independent transgenic lines are shown. Immunodetection was with a monoclonal antibody (mAb) directed against GFP.

To further characterize the effect of the *nTN1* sequence, we fused the sequence directly to the heterologous *VENUS* sequence, yielding *nTN1–VENUS* (Fig. 2a). VENUS appeared an appropriate heterologous target for this approach since the protein matures efficiently and exhibits reduced environmental sensitivity (Nagai *et al*., 2002). Fusion of the *nTN1* sequence to the *VENUS* 5’-end led to increased protein levels as disclosed by both immunodetection and enhanced fluorescence in *nTN1–VENUS*-overexpressing leaf tissue (Fig. 2c and d). We also shifted the position of the *nTN1* sequence yielding constructs *N1b– nTN1–VENUS* and *VENUS–nTN1*. However, at an internal or a 3’-end position, the *nTN1* sequence remained ineffective in the tested constructs (data not shown).

To reappraise results from transient overexpression studies, we integrated constructs stably into the tobacco genome using Agrobacterium-mediated transformation. Protein accumulation was analyzed in crude extracts from whole seedlings and from young leaf tissue in the T1 progeny of independent transgenic lines. Consistently, nTN1–N1b–VENUS fusion protein accumulated to significantly higher levels than N1b–VENUS, and nTN1–VENUS was more prominent than VENUS (Fig. 2e; Fig. S6b and c). Treatment of leaves with SA or the functional analog BTH (benzothiadiazole; Friedrich *et al*., 1996; Görlach *et al*., 1996) or induction of the HR by infection with TMV did not affect accumulation of nTN1-harboring fusion proteins (Fig. S6d and data not shown). Together, the data underpin the notion that the *NIMIN1* N-terminal coding sequence functions as an autonomous structural motif that fosters accumulation of fused proteins in plants.

### Co-expression of *NIMIN1* strengthens accumulation of NPR1 protein

In previous work, we established that overexpression of *NIMIN1* in transgenic Arabidopsis and transient overexpression in *N. benthamiana* suppressed SA-induced *PR-1* gene expression (Weigel *et al*., 2005; Hermann *et al*., 2013). Similar results were reported for the rice NIMIN2 homolog NRR (Chern *et al*., 2005, 2008). In our current work, we recognized yet two additional features of NIMIN1 that appear crucial to understand the functional significance of NIMIN1 liaisons in the course of the plant defense reaction.

First, we mixed Agrobacterium strains harboring *NPR1* or *NPR1–GFP* under control of the *CaMV 35S* promoter either with a strain harboring *35S:NIMIN1* or with control Agrobacteria without gene construct. The suspensions were infiltrated in the left (L) and right (R) halves of *N. benthamiana* leaves, respectively. Accumulation of NPR1 (fusion) protein was monitored in crude leaf extracts using a polyclonal antiserum directed against 6xHis– AtNPR1 (Stos-Zweifel *et al.,* 2018). With *NIMIN1* co-expression, we consistently observed increased levels of NPR1 and NPR1–GFP full-length proteins and their degradation products (Fig. 3a and b; Fig. S7a). In contrast, co-expression of *NIMIN2* or *N1b–VENUS* yielded lower NPR1 (fusion) protein levels (Fig. 3b and Fig. S7b; data not shown). To enable quantification of the positive effect of NIMIN1 on NPR1, we constructed *35S:NPR1–GUS* and *35S:NtNPR1–GUS* reporter genes (Fig. 3c; Fig. S7c to e). In yeast, NIMIN1 was able to bind to NPR1, NtNPR1 and NtNPR1–GUS, and, in all cases, interaction was sensitive to SA (Fig. S7f; Maier *et al*., 2011). Co-infiltration of Agrobacteria harboring *35S:NPR1–GUS* constructs with Agrobacteria carrying *35S:NIMIN1* in *N. benthamiana* leaf tissue yielded reporter activities increased by a factor of about 2 for AtNPR1–GUS and even by a factor of 2.9 for NtNPR1–GUS as compared to co-infiltrations with the wild-type Agrobacterium strain (Fig. 3d and e; Fig. S7g). This gain in GUS enzyme activity by *NIMIN1* co-expression is compatible with elevated levels of NPR1 and NPR1–GFP proteins recorded by immunodetection. Yet, (Nt)NPR1–GUS activities were not increased upon co-expression of *NIMIN2* or *NIMIN3* (Fig. 3d; Fig. S7g). The data suggest that presence of NIMIN1 moderately strengthens accumulation of NPR1 protein *in planta*.

**Fig. 3.**
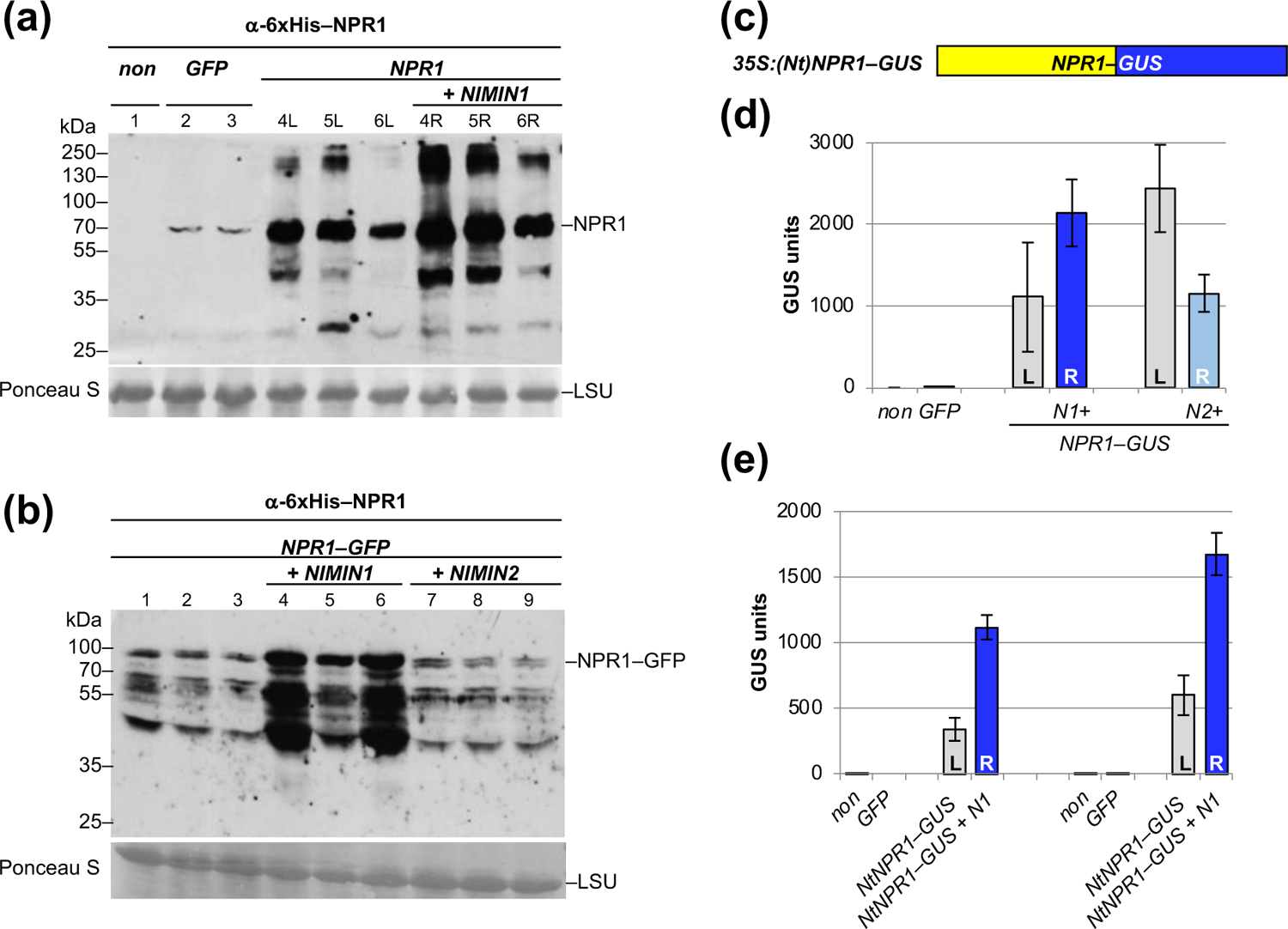
N*I*MIN1 co-expression strengthens accumulation of NPR1 *in planta*. (a) Accumulation of NPR1 with and without co-expression of *NIMIN1* in *N. benthamiana*. For direct comparison of protein accumulation, mixtures of Agrobacterium strains were infiltrated in the left (L) and right (R) halves of the same leaves as indicated. Three plants were infiltrated in parallel for each combination. Immunodetection in leaf extracts was with a polyclonal antiserum directed against 6xHis–AtNPR1. Staining of the large subunit of RUBISCO (LSU) with Ponceau S demonstrates loading of the nitrocellulose filter. (b) Accumulation of NPR1–GFP with and without co-expression of *NIMIN1* or *NIMIN2* in *N. benthamiana*. Three plants were infiltrated in parallel for each construct. (c) Schematic representation of *NPR1–GUS* reporter constructs. (d) Reporter activity of NPR1–GUS fusion protein with and without co-expression of *NIMIN1* or *NIMIN2* in *N. benthamiana*. For direct comparison of GUS enzyme activities, mixtures of Agrobacterium strains were infiltrated in the left (L) and right (R) halves of the same leaves as indicated. Three plants were infiltrated in parallel. (e) Reporter activity of NtNPR1–GUS fusion protein with and without co-expression of *NIMIN1* in *N. benthamiana*. Results from two independent experiments are shown.

### Overexpression of *NIMIN1* promotes emergence of cell death in tobacco

When we monitored *N. benthamiana* plants infiltrated with Agrobacteria harboring *35S:NIMIN1* on a regular basis over a period of two weeks, we observed yet another so far unrecognized quality of NIMIN1. At the earliest 5 dpi, confined tissue collapse became apparent. Then, small brown spots developed growing into larger patches which typically ended in death of the whole infiltrated leaf area (Fig. 4a). At the same time, leaf tissue accumulating GFP remained inconspicuous (Fig. 4a). Phenotypically, NIMIN1-related cell death was indistinguishable from cell death caused by overexpression of the gene for the pro-apoptotic Bcl-2-associated X protein (BAX; Oltvai *et al*., 1993; Fig. 4a) which we put under control of the *NIMIN1* promoter. The *NIMIN1* promoter is not active constitutively in bacteria and plants, but switched on in plant tissue after agroinfiltration (Glocova *et al*., 2005). Cell death manifestation by *NIMIN1* expression was not due to unrestricted proliferation of Agrobacteria or other microorganisms (Fig. S8) and coincided with accumulation of both phenolic compounds visualized by yellow tissue fluorescence and H_2_O_2_ visualized by 3,3’-diaminobenzidine (DAB) staining (Fig. 4b and c). As judged from side-by-side infiltrations in the right and left halves of the same leaves, an even stronger cell death-promoting effect was produced by overexpression of *NIMIN1–VENUS* in comparison to *NIMIN1*, both under control of the *CaMV 35S* promoter (Fig. 4d). NIMIN1–VENUS was able to bind to NPR1 and TPL(1-333) in Y2H assays, and interaction with NPR1 was sensitive to SA (Fig. S9). Likewise, overexpression of some tobacco *NIMIN2*-like genes, f.e., *NtN2c* and *NtN2d*, and their *VENUS* fusion genes elicited cell death in *N. benthamiana* (Fig. S10a to c, f and g).

**Fig. 4.**
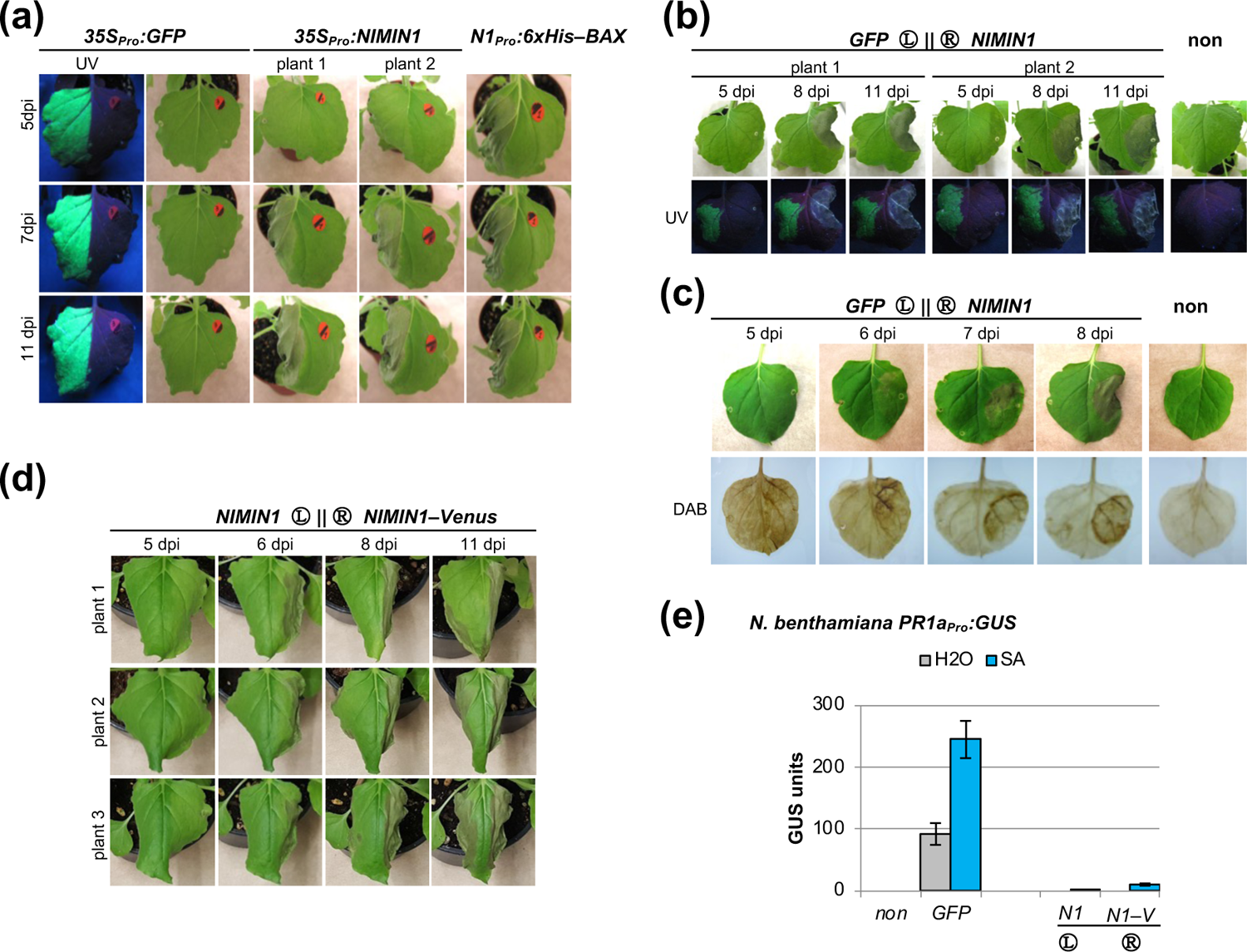
N*I*MIN1 overexpression promotes emergence of cell death *in planta*. (a) Phenotype of *Nicotiana benthamiana* leaves overexpressing *NIMIN1*. Agrobacteria were infiltrated in the left leaf halves. Phenotypes were recorded at different days post infiltration (dpi). Effects of *NIMIN1* overexpression are compared to overexpression of *GFP* and *BAX*. (b) Overexpression of *NIMIN1* produces tissue fluorescence. Agrobacteria harboring *35S:GFP* or *35S:NIMIN1* were infiltrated in the left (L) and right (R) halves of the same leaves as indicated. Photographs were taken under white light and under UV light. (c) Overexpression of *NIMIN1* coincides with accumulation of H_2_O_2_. Agrobacteria harboring *35S:GFP* or *35S:NIMIN1* were infiltrated in the left and right halves of leaves, respectively. Accumulation of H_2_O_2_ is visualized by staining with 3,3’-diaminobenzidine (DAB). (d) Phenotype of *Nicotiana benthamiana* leaves overexpressing *NIMIN1–VENUS*. Agrobacteria harboring *35S:NIMIN1* or *35S:NIMIN1–VENUS* were infiltrated in the left and right halves of leaves, respectively. (e) Salicylic acid-mediated *PR-1a* gene induction is suppressed in leaf tissue overexpressing *NIMIN1* or *NIMIN1– VENUS*. Agrobacteria were infiltrated in the left (L) and right (R) halves of the same leaves of *N. benthamiana* plants harboring a *-1533PR1a_Pro_:GUS* transgene as indicated. Three plants were infiltrated in parallel. GUS reporter activity was determined after floating of disks excised from infiltrated leaf areas on water or 1 mM SA. GUS activities are compared to enzyme activity in *GFP* overexpressing tissue.

However, overexpression of *NIMIN2* or *N1b* or their *VENUS* fusion genes did not produce macroscopic symptoms nor H_2_O_2_ (Fig. S11a, b and e; data not shown). This might be due to the fact that NIMIN2 and N1b fusion proteins did not accumulate strongly in leaf tissue (Fig. S11c; Fig. 2b). Indeed, overexpression of *nTN1–N1b–VENUS*, yielding elevated protein levels (Fig. 2b), promoted cell death similar to *NIMIN1–VENUS* (Fig. S11f).

Necrosis was also induced when we infiltrated *N. benthamiana* plants harboring a *PR1a_Pro_:GUS* transgene (Hermann *et al*., 2013) with *35S:NIMIN1* Agrobacteria. As reported previously, SA-mediated *PR1a_Pro_:GUS* induction was severely suppressed in *NIMIN1* overexpressing leaves, even when samples were taken at 4 or 5 dpi from tissue not yet visibly affected by cell death symptoms (Fig. 4e; Fig. S10d; Hermann *et al*., 2013). Likewise, in tissue determined to cell death by overexpression of *N1–VENUS*, *NtNIMIN2c*, *NtN2c–VENUS* or *Bax*, *PR1a* promoter activity was clearly impaired (Fig. 4e; Fig. S10d and e; data not shown). In contrast, overexpression of *NIMIN2* and *N2–VENUS* did not inhibit transcription from the *PR1a* promoter (Fig. S11d; Hermann *et al*., 2013).

To validate the cell death-promoting activity of NIMIN1, we generated transgenic tobacco plants with the *35S:NIMIN1–VENUS* construct. Initially, primary transformants carrying the transgene grew normally. However, at later stages, transformants tended to shed flowers and seed capsules resulting in an overall low seed production. The selfed progeny of *35S:NIMIN1–VENUS* transformants developed even more severe defects. Plants exhibited abnormal growth phenotypes with asymmetric statures, malformed and distorted leaves and furcated shoots (Fig. S12a to c). Furthermore, cell death symptoms developed spontaneously on leaves and shoots of plants, in particular at the shoot tip, confining flower setting and seed production (Fig. S12d and e). Altogether, plants were stunted compared to control transgenic plants raised under the same conditions (Fig. S12f). These phenotypes were correlated with accumulation of NIMIN1–VENUS protein in leaf tissue, but not with presence of PR-1 protein (Fig. S12g; data not shown).

### The EAR motif is instrumental in cell death induction by NIMIN proteins

To analyze whether the cell death-inducing activity could be assigned to one of the identified domains of NIMIN1, we tested diverse constructs with mutant genes fused to *VENUS* using the agroinfiltration assay. Initially, we used mutant genes with alterations in a single domain targeting just one activity of multi-functional NIMIN1 (single domain mutants). Results were, however, not always clear-cut and difficult to evaluate. Next, we tested mutant *NIMIN1* genes harboring amino acid exchanges in two or three domains (double and triple domain mutants; Table S1).

Overexpression of double domain mutant *N1m2–VENUS*, which abrogates binding to NPR1, still elicited cell death similar to *NIMIN1–VENUS* and clearly suppressed transcription from the *PR-1a* promoter (Fig. S13a to c). Thus, interaction with NPR1 does not seem essential for the cell death-inducing property of NIMINs. On the other hand, triple domain mutant *N1m3–VENUS*, devoid of NPR1 and TPL binding motifs (Fig. S13g and h), did not produce necrosis nor substantial *PR-1a* promoter suppression (Fig. S13d to f). Furthermore, cell death emergence was clearly compromised with EAR motif mutants *N1 L138/140A*, *N1 L138/140A–VENUS* and *NtN2cΔEAR–VENUS* (Fig. S13i and k; Fig. S14). Taken together, these data pointed to the EAR motif as a potential trigger for cell death mediated by NIMIN proteins.

To support this notion, we took a gain-of-function approach. We added the sequences for isolated intact NIMIN1 domains, namely the NLS, the main (blue) NPR1 interaction domain and the EAR motif, to the 3’-end of the strongly expressed *nTN1–VENUS* fusion gene (Fig. 5a). The nTN1–VENUS–NPR1BD protein did not accumulate to substantial amounts neither in yeast nor in plants (Fig. S15a and b). However, the EAR motif bearing fusion gene *nTN1–VENUS–N1EAR* was expressed steadily (Fig. 5b; Fig. S15a and c). The chimeric protein interacted with TPL(1-333) and with TPL(1-194) in yeast. Interaction of nTN1– VENUS–N1EAR with both TPL deletions was even stronger than interaction of NIMIN1 (Fig. 5c; Fig. S16). Furthermore, transient overexpression of *nTN1–VENUS–N1EAR* clearly produced cell death in *N. benthamiana* leaves similar to *NIMIN1–VENUS*, while overexpression of *nTN1–VENUS* or *nTN1–VENUS–NLS* did not elicit visible symptoms within a time lapse of 12 days (Fig. 5d; Fig. S17). Similarly, overexpression of a construct harboring the sequence for the EAR motif of NtN2d fused to *nTN1–VENUS*, *nTN1–VENUS– N2dEAR*, elicited leaf cell death (Fig. 5e and f). Interestingly, both tobacco cell death-inducing proteins, NtN2d and NtN2c, harbor EAR motifs of the DLNxxP-type which, in NtN2d, partly overlaps with an LxLxL motif (Fig. 5g).

**Fig. 5.**
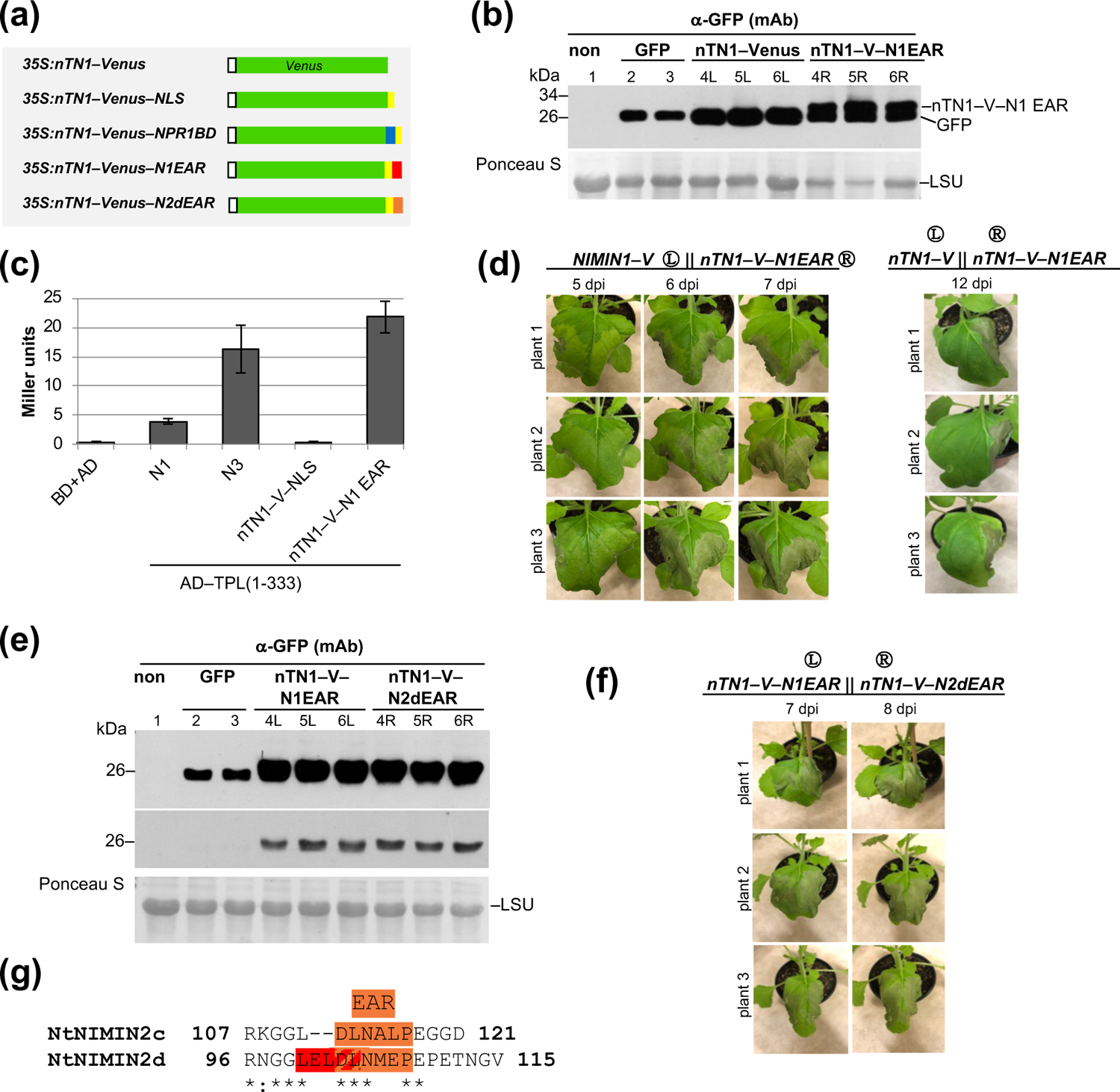
The EAR motif is instrumental in cell death induction via NIMIN proteins. (a) Schematic representation of *nTN1–VENUS* fusion constructs. Colored boxes denote conserved structural motifs of NIMIN1 and NtN2d. Nuclear localization signal (NLS, yellow); interaction motif mediating N1 binding to the SA sensitive C-terminus of NPR1 (blue); EAR motif (red, orange). (b) Accumulation of nTN1– VENUS–N1EAR fusion protein in *Nicotiana benthamiana*. Accumulation of nTN1–VENUS–N1EAR is compared to accumulation of nTN1–VENUS. Three plants were agroinfiltrated in parallel. Immunodetection of VENUS fusion proteins in leaf extracts was with a monoclonal antibody (mAb) directed against GFP. Staining of the large subunit of RUBISCO (LSU) with Ponceau S demonstrates loading of the nitrocellulose filter. (c) Y2H interaction of nTN1–VENUS–N1EAR with TPL(1-333). (d) Phenotype of *Nicotiana benthamiana* leaves overexpressing *nTN1–VENUS–N1EAR*. Agrobacteria were infiltrated in the right halves of leaves. Effects of *nTN1–VENUS–N1EAR* overexpression are compared to overexpression of *NIMIN1–VENUS* and overexpression of *nTN1–VENUS*. (e) Accumulation of nTN1–VENUS–N2dEAR fusion protein in *Nicotiana benthamiana*. Accumulation of nTN1–VENUS– N2dEAR is compared to accumulation of nTN1–VENUS–N1EAR. Three plants were agroinfiltrated in parallel. Two different exposures of the X-ray film are shown. (f) Phenotype of *Nicotiana benthamiana* leaves overexpressing *nTN1–VENUS–N2dEAR*. Agrobacteria were infiltrated in the right halves of leaves. Effects of *nTN1–VENUS–N2dEAR* overexpression are compared to overexpression of *nTN1– VENUS–N1EAR*. (g) Sequences of the EAR motifs in tobacco NIMIN2c and NIMIN2d.

### NIMIN1 interacts with TOPLESS-RELATED3 *in planta*

Current evidence clearly links NIMIN1 to SAR induced by SA and associated with cell survival in the non-infected leaves of plants exhibiting HR. Inconsistent with this notion, our results demonstrated that NIMIN1 promotes cell death in tobacco plants. Cell death emergence depended on the TPL-interacting EAR motif of NIMIN1 and not on interaction of NIMIN1 with NPR1. It thus seemed likely that NIMIN1 still contacts other proteins apart from NPR1 *in planta*. To identify additional interaction partners of NIMIN1, we used proximity labeling by TurboID (Branon *et al*., 2018; Mair *et al*., 2019).

The coding sequence for *NIMIN1* was cloned in a plant binary vector in frame with the sequences for Turbo and YFP (*T–N1–YFP*; Fig. 6a). YFP served as a reporter, while Turbo is a mutant version of *E. coli* BirA coding for a biotin ligase that adds biotin tags to endogenous proteins within a few nanometers of distance to the ligase. As a negative control, we used a vector harboring the Turbo sequence fused to sequences for YFP and the SV40 NLS (*T–YFP– NLS*). Of note, *NIMIN1* codes for an intrinsic NLS (Fig. 6a). *N. benthamiana* plants were infiltrated with Agrobacterium strains harboring these constructs, and crude leaf extracts were monitored for expression of fusion genes by immunodetection with the GFP monoclonal antibody. The T–YFP–NLS fusion protein accumulated to similar levels as NIMIN1– VENUS, while T–N1–YFP was even more prominent (Fig. 6b). Both fusion proteins located to nuclei of epidermal cells (Fig. S18; Mair *et al*., 2019). We also found that expression of *T– N1–YFP*, but not *T–YFP–NLS*, induced cell death in *N. benthamiana* leaves, albeit cell death emergence appeared less severe than with *NMIN1–VENUS* observed previously (Fig. 6c). Nevertheless, this observation indicated that NIMIN1 functions as expected in context of the biotin ligase fusion protein.

**Fig. 6.**
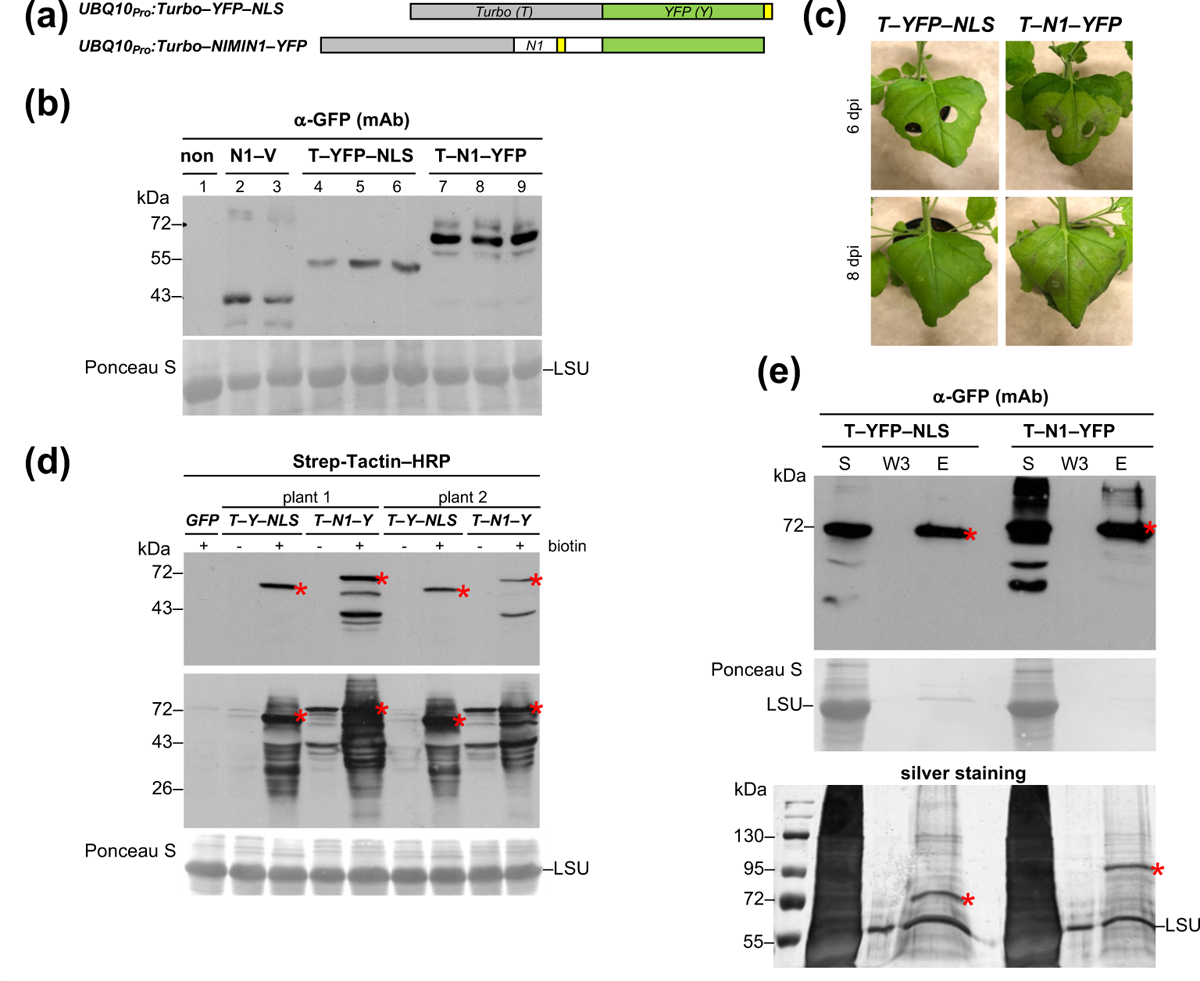
Proximity-dependent labeling of *Nicotiana benthamiana* proteins by NIMIN1–TurboID. (a) Schematic representation of TurboID fusion constructs. The diagram is drawn approximately to scale. Nuclear localization signals (NLS) are marked yellow. *Turbo–NIMIN1–YFP* harbors the NIMIN1 intrinsic NLS, while *Turbo–YFP–NLS* contains the SV40 NLS sequence added at the C-terminus of *YFP*. (b) Overexpression of TurboID fusion genes in *N. benthamiana*. Accumulation of Turbo–YFP– NLS (T–YFP–NLS) and Turbo–NIMIN1–YFP (T–N1–YFP) fusion proteins in *N. benthamiana* in comparison to accumulation of NIMIN1–Venus (N1–V) fusion protein. Three plants were infiltrated in parallel with Agrobacteria harboring the TurboID constructs. Immunodetection in leaf extracts was with a monoclonal antibody (mAb) directed against GFP. Staining of the large subunit of RUBISCO (LSU) with Ponceau S demonstrates loading of the nitrocellulose filter. (c) Phenotype of *Nicotiana benthamiana* leaves overexpressing *Turbo–NIMIN1–YFP*. Effects of *T–N1–YFP* overexpression are compared to overexpression of *T–YFP–NLS*. Results from two separate experiments are shown. (d) Detection of biotinylated proteins in *N. benthamiana* expressing TurboID constructs. *N. benthamiana* leaves were infiltrated in one half with Agrobacteria harboring *T–YFP–NLS* and in the other half with Agrobacteria harboring *T–N1–YFP*. Leaf disks were cut from infiltrated leaf areas and vacuum-infiltrated with water (-) or biotin (+) as indicated. Crude protein extracts were separated on SDS gels and biotinylated proteins were detected by Strep-Tactin–HRP blotting. Two different exposures of the X-ray film are shown. Red asterisks denote ligase self-biotinylation of the respective fusion protein. (e) TurboID pull-down. Biotinylated proteins were purified from crude extracts of *N. benthamiana* plants expressing TurboID constructs with Strep-Tactin-coated beads. Aliquots from the supernatant (S), the third wash fraction (W3) and the eluate (E) were separated on SDS gels and ligase fusion proteins were detected by anti-GFP(mAb) blotting and by silver staining of the gel. Red asterisks denote the positions in the gel of the respective fusion protein. In addition to the fusion proteins, the large subunit of RUBISCO (LSU) is detected in the Strep-Tactin pull-down by silver staining.

Next, we treated disks cut from agroinfiltrated *N. benthamiana* leaves with water or biotin and subjected the extracted proteins to detection with Strep-Tactin–HRP conjugate that binds to biotinylated proteins. In addition to the ligase fusion proteins, which self-biotinylate, we detected a range of bands chiefly with the biotin-treated samples (Fig. 6d). Pull-down with Strep-Tactin-coated beads yielded both ligase fusion proteins in an about even distribution in the respective eluates as visualized by anti-GFP blotting and by silver staining of SDS gels (Fig. 6e). Together, leaf extracts from *T–N1–YFP* overexpression appeared an excellent source for identification of *in planta* NIMIN1 interaction partners. Using an antiserum directed against AtNPR1, we detected the endogenous *N. benthamiana* protein in crude extracts. However, we were not able to demonstrate NPR1 in eluates from Strep-Tactin-coated beads (Fig. S19).

Harvested biotinylated proteins from overexpression of *T–YFP–NLS* and *T–N1–YFP* were identified by mass spectrometry (MS). The analysis was performed twice with crude extracts from the same agroinfiltrations (technical replicates) and additionally with a crude extract from a separate infiltration experiment (biological replicate). Altogether, about 200 proteins were identified for both baits at a peptide threshold of 95%, a protein threshold of 99% and a minimum number of four unique peptides (Keller *et al*., 2002; Nesvizhskii *et al*., 2003). As a matter of fact, most proteins were represented about evenly in the samples, f.e., ribulose-1,5-bisphosphate-carboxylase/oxygenase (RUBISCO) large subunit (LSU) and small subunit (SSU; Table S2). This is consistent with our observation that LSU was also identified on silver-stained SDS gels in the enriched biotinylated protein fractions from both *Turbo– YFP–NLS* and *Turbo–N1–YFP* overexpressing plants (Fig. 6e and S19). Evenly distributed proteins do not constitute specific binding partners of NIMIN1. In contrast, only a few proteins were identified without record in the T–YFP–NLS samples and a minimum of four unique peptides in the T–N1–YFP samples. These hits are listed in Table 1. First of all, the NIMIN1 bait protein was identified in all samples, except the negative controls, based on 11 to 13 unique peptides (Fig. S20). In addition, there was only a single hit which was common to all samples with the T–N1–YFP bait, i.e., TPL-RELATED3 (TPR3). The protein was represented in the different samples by 5 to 14 unique peptides (Fig. S21). Two peptides represented protein regions highly dissimilar among TPL family members, and identified peptides were derived from the N-terminal as well as the C-terminal end of TPR3 indicating that TPR3 is a likely NIMIN1 interactor *in planta*. Consistent with the failure of immunodetection, NPR family proteins were not revealed by MS analyses from agroinfiltrated *N. benthamiana* plants.

**Table 1.**
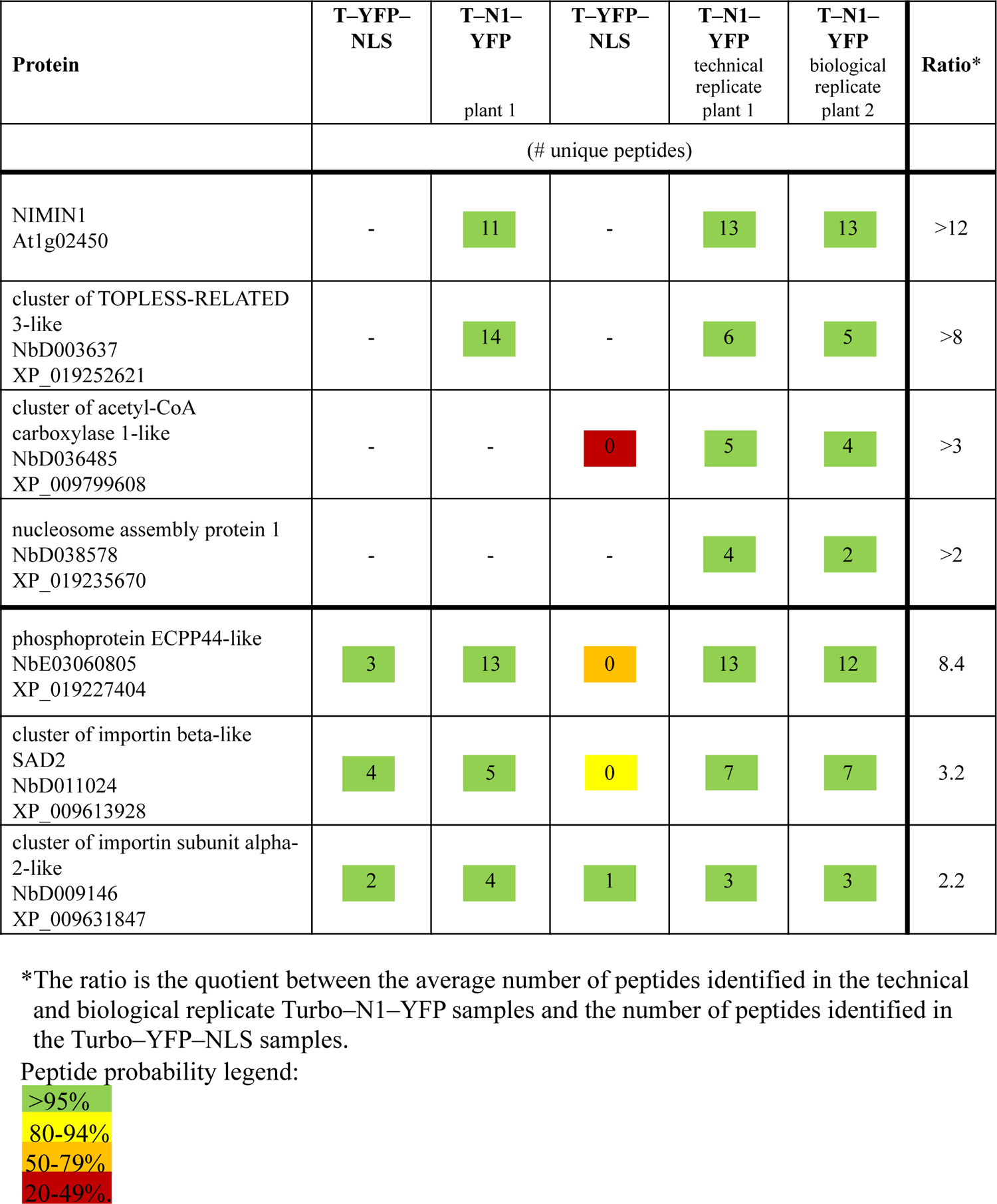
Proteins identifed by mass spectrometry with NIMIN1–TurboID in agroinfiltrated.

### *In planta* interaction of NIMIN1 with NPR1

*In planta* interaction of NIMIN1 with TPR3 as revealed by Agrobacterium-mediated transient overexpression of Turbo–NIMIN1–YFP was not surprising as TPR3 belongs to the TPL family of proteins which all harbor the EAR motif-interacting TPL domain (Ke *et al*., 2015; Martin-Arevalillo *et al*., 2017). Furthermore, NIMIN1 binding to TPL was already known in yeast (Arabidopsis Interactome Mapping Consortium *et al*., 2011; Fig. S3). Nevertheless, lack of identification of the NPR1 interaction partner in our analyses was somewhat disturbing. We therefore transferred the *Turbo–YFP–NLS* and *Turbo–NIMIN1–YFP* constructs in the tobacco genome to enable protein-protein interaction studies in non-challenged plants.

Expression of fusion proteins and biotin-labeling of proteins was checked in two T1 plants each from transgenic lines 167H-7 (*UBQ10_Pro_:T–YFP–NLS*) and 165H-5 (*UBQ10_Pro_:T–N1–YFP*). Unfortunately, plant 167H-7/1 did not accumulate the T–YFP–NLS protein (Fig. S22a). Also, biotinylation occurred not only in biotin-fed leaf disks, but also in controls floated on water, albeit less efficiently (Fig. S22b). This is likely due to endogenous biotin in the plants. We used the crude extract from plant 167H-7/2 as negative control and extracts from plants 165H-5/1 and 165H-5/2 for identification of NIMIN1 interaction partners in young biotin-treated *N. tabacum* leaf tissue (technical and biological repeat). Levels of Turbo fusion proteins and yield of biotinylated proteins in the Strep-Tactin pull-down were clearly lower with extracts from transgenic plants than with extracts from agroinfiltrated *N. benthamiana*. We were not able to allocate bands for the fusion proteins in the pull-down samples on silver-stained SDS gels. In the MS analyses, only 14 tobacco proteins were identified in total for the two baits when we applied the same stringency as used in the transient expression experiments (minimum of four unique peptides). Most proteins were about evenly distributed in the samples (Table S3). Only two proteins were not detected in the negative control and represented by at least four unique peptides in all NIMIN1 bait samples. These were NIMIN1 and importin subunit alpha-2-like (Table 2). Applying lower stringency in protein identification revealed a few more proteins including NPR family members NIM1-like1 and NPR1 (Table 2; Fig. S23) which both interact strongly with NIMIN1 in yeast (Maier *et al*., 2011). Most notably, however, TPR3 and other members of the TPL family were not identified in any extract from the transgenic plants. It is also worth mentioning that NIMIN1–TurboID MS analyses disclosed interaction of the NIMIN1 bait with various components of the nuclear trafficking machinery in both *N. benthamiana* and *N. tabacum* (Tables 1 and 2).

**Table 2.** Proteins identified by mass spectrometry with NIMIN1–TurboID in transgenic *Nicotiana tabacum* plants.

| Protein | T–YFP–<br>NLS<br>167H-7/2 | T–N1–<br>YFP<br>165H-5/1 | T–N1–<br>YFP<br>biological<br>replicate<br>165H-5/2 | T–N1–<br>YFP<br>technical<br>replicate<br>165H-5/2 | Ratio* |
| --- | --- | --- | --- | --- | --- |
|  | (# unique peptides) |  |  |  |  |
| NIMIN1<br>At1g02450 | - | 4 | 4 | 5 | >4.3 |
| cluster of importin subunit alpha-2-like<br>XP_009631847 | 0 | 4 | 6 | 4 | >4.7 |
| pyridoxal phosphate-dependent<br>transferase<br>XP_016473982 | - | 1 | 2 | 2 | >1.7 |
| NIM1-like1 (NPR3-like)<br>AY640382 | - | 2 | 0 | 1 | >1 |
| NPR1<br>KY402167 | - | - | 1 | 0 | >0.3 |
| exportin 1A-like<br>XP_016460061 | 1 | 3 | 2 | 1 | 2.0 |

\*The ratio is the quotient between the average number of peptides identified in the Turbo–N1–YFP samples and the number of peptides identified in the Turbo–YFP–NLS sample.
Peptide probability legend:

### Overexpression of the N-terminal TOPLESS fragment produces cell death in *Nicotiana benthamiana*

Given the fact that *NIMIN1* overexpression promotes cell death in plants and that the EAR motif is instrumental in this process, we would anticipate that the EAR motif interactor TPL may be involved in elicitation of cell death *in planta*.

To substantiate our idea, we overexpressed *TPL* and N-terminal *TPL(1-333)* under control of the *CaMV 35S* promoter next to *NIMIN1–VENUS* or *nTN1–VENUS–N1EAR* in *N. benthamiana*. While infiltration of Agrobacteria harboring the *TPL* gene did not produce visible symptoms on leaves, expression of *TPL(1-333*) elicited cell death. Cell death emergence was similar to cell death induction by expression of *NIMIN1–VENUS* and clearly stronger than cell death induction by expression of *nTN1–VENUS–N1EAR* (Fig. S24). The data indicate that signaling through the N-terminal third of TPL takes control over cell growth and death in agroinfiltrated *N. benthamiana* plants.

## Discussion

NIMIN1 to NIMIN3 were identified as members of a novel protein family in a single yeast two-hybrid screen by virtue of interaction with the positive SAR regulator NPR1. Although sharing distinct domains, there are nevertheless some striking differences among them. For example, both *NIMIN1* and *NIMIN2* are induced by SA. Yet, induction of *NIMIN1*, unlike *NIMIN2*, is dependent on NPR1, and *NIMIN1* is activated only transiently after *NIMIN2*.

Hence, *NIMIN1* classifies as an early SA-induced gene while *NIMIN2* and *PR-1* are immediate early and late SA-induced genes, respectively. Together, the data indicate that NIMIN1 and NIMIN2 may engage in different processes with NIMIN1 playing a role after NIMIN2 past the onset of SA signaling.

### NIMIN1 is a multi-domain protein with diverse capabilities

A major function of NIMIN1 is interaction with the SAR regulator NPR1. In our previous work we suggested that NIMIN2 and NIMIN1 control defense gene induction through NPR1 in Arabidopsis (Hermann *et al*., 2013; Konopka *et al*., 2022). Both NIMIN2 and NIMIN1 bind via a shared domain (blue domain in Fig. 1) near the SA perception site in the C-terminus of NPR1 thus rendering NPR1 receptive to the SA signal. In addition to this main interaction domain, NIMIN1 harbors still another potential NPR1 binding site (green domain in Fig. 1; EDF motif) which is not present in NIMIN2. In Y2H assays, this domain mediates at most weak interaction of NIMIN1 with NPR1. Its functional significance for NIMIN1 activity is currently unresolved. However, NIMIN3 comprises a similar sequence motif by which NIMIN3 binds to the N-terminal region of NPR1 (Weigel *et al*., 2001). Of note, interaction of NIMIN3 with NPR1, unlike NIMIN1 and NIMIN2 binding, is not sensitive to SA thus suggesting a function for NIMIN1 beyond its role in SA perception by NPR1.

NIMIN1 also harbors an EAR motif at its C-terminal end for interaction with TPL and a domain facilitating accumulation of NIMIN1 protein at its N-terminus. These four functional domains we identified in NIMIN1 are neatly separated from each other by short disordered regions indicating that the included structural motifs could function *in planta* without mutual interference in specific and diverse processes mediated by NIMIN1.

### NIMIN1 protein accumulates in threatened tissue

As judged from immunodetections, both NIMIN1 and NIMIN2 are unstable proteins that undergo degradation in plant tissue. However, due to a unique sequence at the N-terminus, NIMIN1 accumulates to higher levels than NIMIN2. This sequence stretch, termed *nTN1*, is also able to reinforce accumulation of NIMIN1b and of a heterologous protein, VENUS, when fused to their N-terminal ends. Enhanced accumulation of nTN1-harboring fusion proteins occurred upon transient overexpression and in transgenic plants and was not affected by biotic stress. The mechanism how the *nTN1* sequence potentiates protein accumulation is not clear. We used constructs with the constitutive *CaMV 35S* promoter fused directly to the translation start codon of the *NIMIN1* structural gene. Thus, protein accumulation was not under control of upstream open reading frames (uORFs) positioned 5’ to the *NIMIN1* translation start (Pajerowska-Mukhtar *et al*., 2012; Zhang *et al*., 2020) nor under control of mRNA motifs in the 5’ leader sequence which impact the translational efficiency of transcripts (Xu *et al*., 2017; Wang *et al*., 2022b). Our data would suggest that the *nTN1* sequence acts in a positive way, for example, as translational enhancer. Yet, the N-terminal NIMIN1 amino acids appear to exert some effect also on protein structure. On the one hand, the nTN1–N1b fusion protein, unlike NIMIN1b, was able to interact with TPL(1-333) in yeast, and, on the other hand, the nTN1–VENUS–NPR1BD fusion protein, unlike nTN1– VENUS and nTN1–VENUS–N1EAR, did not accumulate to substantial levels in yeast nor in plants. Taken together with the expression data (Hermann *et al*., 2013), our results qualify NIMIN1 as protein with elevated transient presence after the onset of SA signaling. As such, NIMIN1 binds to the C-terminal region of NPR1 and stabilizes NPR1 protein which undergoes continuous turnover by proteasomal degradation (Spoel *et al*., 2009; Skelly *et al*., 2019). Collectively, NIMIN1 boosts the plant’s response to the SAR signal SA in several ways, first by increasing the dose of NPR1 through NIMIN1-dependent stabilization, second by facilitating hormone sensing and activation of NPR1, and third by accumulation in threatened tissue.

### NIMIN1 accumulation promotes induction of cell death via EAR motif-mediated interaction with TPR3

Quite interestingly, SA is able to expel NIMIN1, but not NIMIN2, from ternary complexes with NPR1 and TGA2 or TGA6 transcription factors in yeast (Hermann *et al*., 2013) corroborating the notion that NIMIN1, apart from binding to the C-terminus of NPR1, may take still other functions in the further course of SAR when SA levels are high. Consistent with this idea, we revealed a novel activity of NIMIN1, i.e., promotion of cell death in tissue overexpressing *NIMIN1* or *N1* fusion genes. In agroinfiltrated *N. benthamiana*, we observed strong cell death while cell death symptoms in stably transformed *N. tabacum* plants were considerably milder. Notably, cell death symptoms were not elicited by *NIMIN2* overexpression. We would not assume that NIMIN1 protein on the whole induces necrosis. Visible cell death emerged at the earliest 5 days after agroinfiltration and coincided with substantial protein accumulation. Rather, we suppose that increased levels of NIMIN1 (fusion) protein disturb homeostasis and thus trigger a signaling cascade targeted at elicitation of cell death. Together, our results support the notion that the TPL-interacting LxLxL-type EAR motif of NIMIN1 plays an essential role in cell death emergence. We also detected cell death upon overexpression of tobacco *NIMIN2c* and *N2d* mediated by their DLNxxP-type EAR motifs. Consistently, using NIMIN1–TurboID proximity labeling, we identified TPR3, a member of the TPL family, as NIMIN1 interaction partner in agroinfiltrated *N. benthamiana* leaf tissue developing cell death. Other members of the tobacco TPL family were not uncovered. Our results are in accordance with a recent report identifying NIMIN1 and TPR3, among other proteins, in the NPR1 nuclear proxiome in SA-treated Arabidopsis (Powers *et al*., 2024). Of note, agroinfiltration is generally linked to elicitation of immune responses, in particular induction of PR-1 protein (Neeley *et al*., 2019), but not linked to emergence of immediate cell death. Hence, accumulation of NIMIN1 (fusion) protein in agroinfiltrated *N. benthamiana* leaves mimics a scenario of acute attack which is concomitant with NIMIN1-TPR3 interaction. In contrast, TPR3 was not revealed as NIMIN1 interaction partner in leaves of young non-challenged transgenic *N. tabacum* plants accumulating Turbo–NIMIN1–YFP. Rather, conditions of low SA levels disclosed proximity of NIMIN1 to NPR family members NIM1-like1 (NPR3-like) and NPR1, which were not identified in the transient assays.

### NIMIN1 shifts the plant response from defense gene activation associated with SAR to cell death

In fact, the TPL co-repressors have essential roles in plant growth and development and in response to extrinsic challenges (Causier *et al*., 2012; Plant *et al*., 2021). Initially, the *TPL* gene was identified from an Arabidopsis mutant with defects in apical embryonic development (Long *et al*., 2002). It was suggested that TPL is necessary to repress the expression of root-promoting genes in the top half of the embryo to allow proper differentiation of the shoot pole (Long *et al*., 2006). Apart from *TPL*, Arabidopsis and tobacco harbor four *TPL*-related genes, *TPR1* to *TPR4*. TPL family proteins interact with numerous transcription factors and adaptor proteins. They are central orchestrators of phytohormone-mediated signal transduction including auxin, gibberellin, brassinosteroid, abscisic acid, jasmonate (JA), SA and ethylene-induced gene expression (Arabidopsis Interactome Mapping Consortium *et al*., 2011; Causier *et al*., 2012; Plant *et al*., 2021). TPL/TPRs are recruited to target genes by EAR motif-bearing proteins with control of gene expression proceeding in an often very similar manner (Causier *et al*., 2012; Plant *et al*., 2021). For example, AUXIN/INDOLE-3-ACETIC ACID proteins (AUX/IAAs) function as transcriptional repressors that interact with promoter-bound AUXIN RESPONSE FACTORs activating transcription of auxin responsive genes. TPL and TPRs are bound by the EAR motifs of AUX/IAAs to act as co-repressors (Szemenyei *et al*., 2008). Increasing auxin levels cause proteasomal degradation of AUX/IAAs leading to release of TPL/TPRs and de-repression of downstream genes (Mockaitis & Estelle, 2008). Similarly, it was demonstrated that overexpression of *NIMIN1* in transgenic Arabidopsis prevents SA induction of *PR-1*, and it was suggested that NIMIN1 functions as a repressor of the NPR1-TGA transcription complex (Weigel *et al*., 2005), possibly via EAR-mediated recruitment of TPL. While this hypothesis seems plausible, there remain, however, some aspects to be kept in mind. First, NIMIN proteins cannot bind directly to promoter-bound TGA transcription factors, but rather exert their activity through interaction with NPR1. The NPR1-TGA transcription complex is inactive in non-challenged plants. NPR1 acts as a co-activator of *PR-1* transcription that must be activated by SA. NIMIN1 is a small SA-induced nuclear protein with increased transient presence that comes into action after NIMIN2. Driven by SA, NIMIN1 can leave the NPR1-TGA transcription complex, and, finally, accumulation of NIMIN1 in threatened tissue leads to strong cell death. Taken together, our data support a model where SA-induced NIMIN1 replaces NIMIN2 in the NPR1-TGA complex to repress NPR1 activity until a threshold level of SA has been reached. Further increases in SA concentration expel NIMIN1 from the NPR1-TGA transcription complex immediately resulting in massive defense gene induction. Accumulating released NIMIN1 can associate with members of the TPL family, in particular TPR3, to signal a severe problem which finally results in local induction of cell death to sacrifice the attacked tissue (Fig. 7). Indeed, we showed that overexpression of TPL(1-333) likewise elicited strong necrosis in *N. benthamiana*. Together, our findings support the idea that TPL/TPR3-mediated signaling is linked to promotion of cell death, and that SA-induced NIMIN1 functions as a crucial agent in this process.

**Fig. 7.**
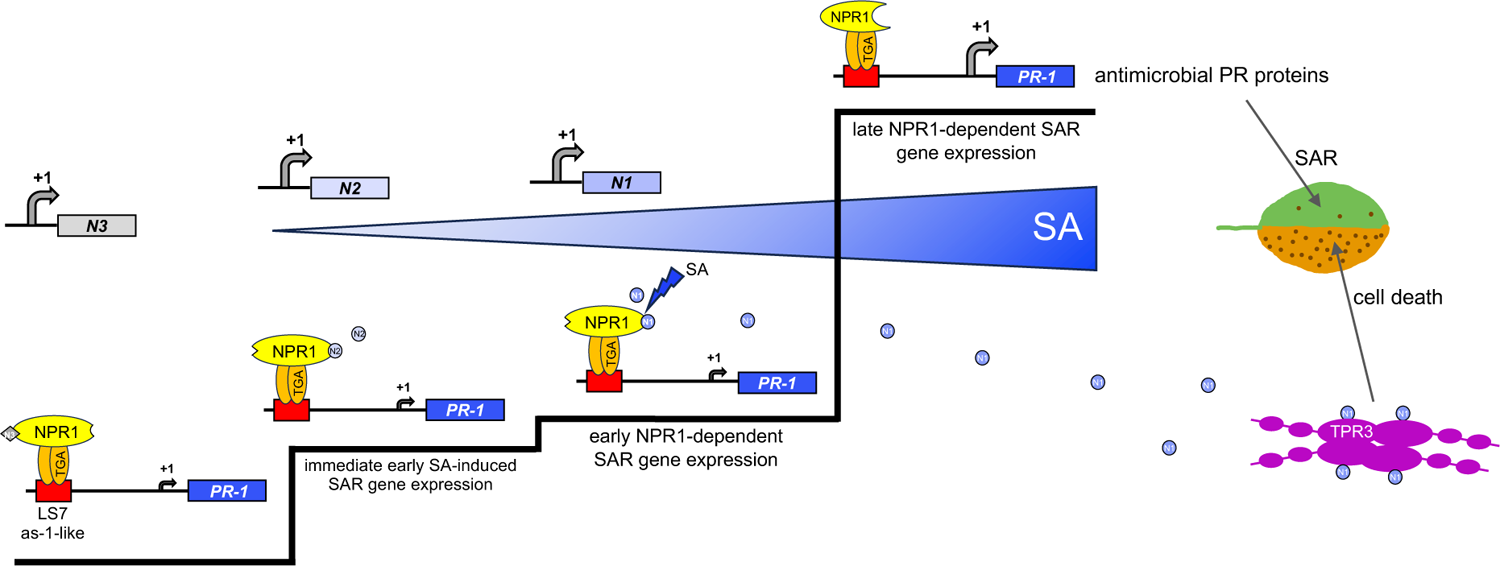
Working model for the function of NIMIN1. This is a revised version of the model presented by Hermann *et al*. (2013). With rising SA concentration, *NIMIN1* is expressed from the NPR1-TGA complex. Accumulating NIMIN1 protein binds to the NIMIN1/NIMIN2-binding domain (N1/N2BD) in the C-terminus of NPR1 to repress late SAR gene expression. At higher hormone concentration, NIMIN1 is outcompeted by SA bridging the LENRV and N1/N2BD regions to induce a conformational switch in the NPR1 C-terminus that induces transcription of late SAR genes, f.e., *PR-1* (Neeley *et al*., 2019). In slightly infected tissue, indicated by a low number of necrotic lesions, antimicrobial PR proteins combat the pathogen to secure survival. In heavily infected tissue, accumulating NIMIN1 is recruited via its EAR motif by TOPLESS-RELATED3 (TPR3) to initiate a cascade finally leading to cell death and tissue sacrifice. TPR3 acts as a tetramer (Ke *et al*., 2015; Dinesh *et al*., 2016; Martin-Arevalillo *et al*., 2017). For simplicity, dimerization of NPR1 (Kumar *et al.,* 2022) is not shown.

In this respect, the phenotypes of transgenic tobacco plants expressing *NIMIN1– VENUS* are of interest. Above all, plants of the T2 and T3 generations exhibited various malformations, spontaneous sporadic cell death symptoms, low seed production and stunted growth. These phenotypes were due to accumulation of NIMIN1–VENUS fusion protein, but not associated with presence of PR-1 proteins excluding that SA signaling was activated in *NIMIN1–VENUS* overexpressing plants. The plant phenotypes are reminiscent of Arabidopsis autoimmune mutants with constitutively elevated SA levels which typically display dwarfism and, in some cases, spontaneous lesion formation (Bowling *et al*., 1997; van Wersch *et al*., 2016). In tobacco, application of exogenous SA can reduce shoot growth and leaf epidermal cell size (Dat *et al*., 2000). Similarly, it was demonstrated that overexpression of *TPR1–Myc* causes dwarfism in transgenic Arabidopsis plants (Zhu *et al*., 2010; Niu *et al*., 2019).

In summary, it is well established that SA not only activates *PR* defense genes (Vlot *et al*., 2009) but impacts plant growth and development, in particular vegetative growth, flowering and seed production (Rivas-San Vicente & Plasencia, 2011; Li *et al*., 2022). SA-mediated growth regulation proceeds, at least in parts, through NPR1 (Vanacker *et al*., 2001; Canet *et al*., 2010a, 2010b) and hence, mediated by NIMIN1, may hit upon TPL/TPRs.

It is also noteworthy that the biotrophic fungus *Ustilago maydis* targets the TPL family of co-repressors as a central hub in infected plants (Darino *et al*., 2021; Bindics *et al*., 2022; Navarette *et al*., 2022). *U. maydis* releases several effectors into the plant cell nucleus that interact with TPL/TPRs thus manipulating auxin and JA/ethylene signaling to promote virulence of the fungus. Binding to TPL family proteins is mediated by EAR motifs of the types LxLxL or DLNxxP or by unknown domains of the fungal effectors, and interactions occur at different sites of TPL/TPRs. Thus, TPL/TPRs are approached, among others, not only by proteins from different phytohormone signaling pathways and by proteins from the SA, jasmonate (JA) and ethylene hormone defense signaling pathways, but also by various pathogen effector proteins. How interference of these manifold interactors is integrated by TPL/TPRs is currently unknown. Irrespective of the precise mechanism, our findings underscore the notion that TPL family proteins are key regulators of growth-defense antagonism in plants (Garner *et al*., 2021; Bindics *et al*., 2022; Navarette *et al*., 2022). Under growing threat by biotrophic pathogens, accumulating SA-induced NIMIN1 can modulate TPR3 signaling to initiate cell death processes at the expense of growth and development which, ultimately, may defeat the invader.

## Conclusion

We suggest that SA-induced NIMIN1 is a dynamic component of the plant pathogen defense system acting as an adaptor protein that goes into interaction with at least two partners, the central SAR regulator NPR1 and the phytohormone orchestrator TPR3. In infected plants, NIMIN1 is the moderator. At first, NIMIN1 competes with SA for binding to the NPR1 C-terminus thus controlling SA-driven activation of *PR-1* defense gene expression. At high SA concentrations, NIMIN1 can shift leaf tissue from cell survival associated with SAR to transition to cell death mediated by its interaction with TPR3. These dual roles of NIMIN1 are recorded in its unique domain structure. Figure 7 summarizes our working model which is a revised version of the model outlined by Hermann *et al*. (2013). Key features are that the physiological response of the plant to pathogen attack occurs coordinated after threshold levels of SA have been reached (Ward *et al*., 1991), and that the response can be adapted to the severity of the assault. Our model is in accordance with the phenomenon of priming (Hermann *et al*., 2013; Konopka *et al*., 2022). It also gives a tentative explanation for SA-mediated and NPR1-dependent cross-talk inhibition of JA defense gene induction (Caarls *et al*., 2015; Proietti *et al*., 2018) since SA-elicited initiation of cell death processes, which, in the end, are aimed at tissue sacrifice, is likely to hamper gene induction events. Finally, our model of dual-purpose NIMIN1 is supported by a recent report suggesting that NPR1 reprograms the transcriptome through multiple steps, instead of through parallel association with multiple transcription factors (Powers *et al*., 2024). Previous work (Arabidopsis Interactome Mapping Consortium *et al.,* 2011) and our work classify SA-induced and NPR1-dependent NIMIN1 as molecular messenger interconnecting the pathogen-induced defense response and the global hormone signaling network that controls plant growth and development thus enabling the plant to cope with growing pathogen pressure and secure reproduction.

## Acknowledgements

We are grateful to Marc Vidal (Harvard Medical School) and the Arabidopsis Biological Resource Center (Ohio State University) for providing pDEST-AD012F12 encoding AtTPL, to Dominique Bergmann (Stanford University) for the generous gift of TurboID vectors R4pGWB601_UBQ10p-Turbo-NES-YFP and R4pGWB601_UBQ10p-Turbo-YFP-NLS and to Andreas Schaller (University of Hohenheim) for providing pBI101.1. We also would like to thank Christine Arnold for generation of gene constructs; Ingrid Prießnitz-Hohos for plant transformation; Berit Würtz for MS analyses; students at the University of Hohenheim, in alphabetical order, Lena Fischer, Verena Häfner, Jaqueline Jung, Stefanie Schäuffele, Philipp Schreiber, Saskia Sommer, Markus Späth, Katharina Wagner and Nicole Wöhrle for producing affirmative evidence. Mass spectrometry-related research reported in this publication was supported by the Core Facility Hohenheim, Mass Spectrometry Unit (University of Hohenheim, Stuttgart, Germany). The Exploris 480 mass spectrometer was funded in part by the German Research Foundation (DFG-INST 36/171-1 FUGG). This work was supported by Land Baden-Württemberg and Universität Hohenheim.

## Author contributions

MS: gene constructs, Y2H, agroinfiltration, analysis of gene expression; KS: gene constructs, Y2H, agroinfiltration, analysis of gene expression, proximity labeling by TurboID; AM: agroinfiltration, DAB staining; PH: mass spectrometry; AJPP: subcellular localization, inestimable advice and support, fruitful discussions; UMP: gene constructs, agroinfiltration, analysis of gene expression, coordination and supervision of the work, writing of the article. All authors read and approved the manuscript.

## Supporting Information

Additional Supporting Information may be found online in the Supporting Information section at the end of the article.

**Fig. S1** Domain structure of Arabidopsis NIMIN1 as deduced from alignment of NIMIN1, NIMIN2 and NIMIN3.

**Fig. S2** Interaction of Arabidopsis NPR1 with NIMIN1 proteins harboring mutations in the two NPR1 binding motifs.

**Fig. S3** Interaction of NIMIN1 and NIMIN3 with Arabidopsis TOPLESS.

**Fig. S4** Domain structure and disorder profile of a chimeric NIMIN1b protein, nTN1– NIMIN1b, harboring the N-terminal region of NIMIN1.

**Fig. S5** Interaction of chimeric nTN1–NIMIN1b with Arabidopsis NPR1 and TPL(1-333).

**Fig. S6** The N-terminal *NIMIN1* sequence *(nTN1)* reinforces protein accumulation *in planta*.

**Fig. S7** *NIMIN1* co-expression strengthens accumulation of NPR1 *in planta*.

**Fig. S8** *NIMIN1* overexpression in *Nicotiana benthamiana* does not affect microbial growth.

**Fig. S9** Interaction of NIMIN1–VENUS with NPR1 and TOPLESS.

**Fig. S10** Overexpression of tobacco *NIMIN2c* and *NIMIN2d* promotes emergence of cell death and *PR-1a* gene suppression *in planta*.

**Fig. S11** Overexpression of *NIMIN2* or *NIMIN1b* does not promote emergence of cell death *in planta*.

**Fig. S12** Phenotypes of *NIMIN1–VENUS* overexpressing transgenic tobacco plants.

**Fig. S13** The EAR motif is instrumental in NIMIN1-mediated cell death induction.

**Fig. S14** The EAR motif is instrumental in NtNIMIN2c-mediated cell death induction.

**Fig. S15** Expression of *nTN1–VENUS–NPR1BD* and *nTN1–VENUS–N1EAR* fusion genes in yeast and in *Nicotiana benthamiana*.

**Fig. S16** Yeast two-hybrid interaction of nTN1–VENUS–N1EAR with TPL(1-194).

**Fig. S17** Overexpression of *nTN1–VENUS* or *nTN1–VENUS–NLS* does not induce cell death in *Nicotiana benthamiana*.

**Fig. S18** Subcellular localization of NIMIN1–TurboID fusion protein in *Nicotiana benthamiana* and *N. tabacum*.

**Fig. S19** Proximity-dependent labeling of *Nicotiana benthamiana* proteins by NIMIN1– TurboID does not reveal NPR1.

**Fig. S20** Peptides of the NIMIN1 bait identified by NIMIN1–TurboID mass spectrometry in *Nicotiana benthamiana*.

**Fig. S21** Peptides of TOPLESS-RELATED3 identified by NIMIN1–TurboID mass spectrometry in *Nicotiana benthamiana*.

**Fig. S22** Proximity-dependent labeling of *Nicotiana tabacum* proteins by NIMIN1-TurboID.

**Fig. S23** Peptides of NIM1-like1 (NPR3-like) and NPR1 identified by NIMIN1-TurboID mass spectrometry in transgenic *Nicotiana tabacum* plants.

**Fig. S24** The N-terminal TOPLESS fragment, TPL(1-333), induces cell death in *Nicotiana benthamiana*.

**Table S1** Domain structures of Arabidopsis NIMIN proteins and mutants used in this work.

**Table S2** Proteins identified by mass spectrometry with NIMIN1-TurboID and TurboID-NLS in agroinfiltrated *Nicotiana benthamiana* plants displaying an approximately even distribution in all samples.

**Table S3** Proteins identified by mass spectrometry with NIMIN1-TurboID and TurboID-NLS in transgenic *Nicotiana tabacum* plants displaying an approximately even distribution in all samples.

**Table S4** Primers used for gene construction.

**Methods S1** Detailed description of methods.

