## Supporting Information for "Arabidopsis NIM1-INTERACTING1 (NIMIN1) is a multi-domain protein controlling transition from systemic acquired resistance (SAR) to cell death"

ORCID ID Ursula M. Pfitzner: 0000-0002-7566-7614

† *Present address: IDT Biologika Dessau-Rosslau, Germany*

**Fig. S1** Domain structure of Arabidopsis NIMIN1 as deduced from alignment of NIMIN1, NIMIN2 and NIMIN3.

**Fig. S2** Interaction of Arabidopsis NPR1 with NIMIN1 proteins harboring mutations in the two NPR1 binding motifs.

**Fig. S3** Interaction of NIMIN1 and NIMIN3 with Arabidopsis TOPLESS.

**Fig. S4** Domain structure and disorder profile of a chimeric NIMIN1b protein, nTN1–NIMIN1b, harboring the N-terminal region of NIMIN1.

**Fig. S5** Interaction of chimeric nTN1–NIMIN1b with Arabidopsis NPR1 and TPL(1-333).

**Fig. S6** The N-terminal *NIMIN1* sequence *(nTN1)* reinforces protein accumulation *in planta*.

**Fig. S7** *NIMIN1* co-expression strengthens accumulation of NPR1 *in planta*.

**Fig. S8** *NIMIN1* overexpression in *Nicotiana benthamiana* does not affect microbial growth.

**Fig. S9** Interaction of NIMIN1–VENUS with NPR1 and TOPLESS.

**Fig. S10** Overexpression of tobacco *NIMIN2c* and *NIMIN2d* promotes emergence of cell death and *PR-1a* gene suppression *in planta*.

**Fig. S11** Overexpression of *NIMIN2* or *NIMIN1b* does not promote emergence of cell death *in planta*.

**Fig. S12** Phenotypes of *NIMIN1–VENUS* overexpressing transgenic tobacco plants.

**Fig. S13** The EAR motif is instrumental in NIMIN1-mediated cell death induction.

**Fig. S14** The EAR motif is instrumental in NtNIMIN2c-mediated cell death induction.

**Fig. S15** Expression of *nTN1–VENUS–NPR1BD* and *nTN1–VENUS–N1EAR* fusion genes in yeast and in *Nicotiana benthamiana*.

**Fig. S16** Yeast two-hybrid interaction of nTN1–VENUS–N1EAR with TPL(1-194).

**Fig. S17** Overexpression of *nTN1–VENUS* or *nTN1–VENUS–NLS* does not induce cell death in *Nicotiana benthamiana*.

**Fig. S18** Subcellular localization of NIMIN1–TurboID fusion protein in *Nicotiana benthamiana* and *N. tabacum*.

**Fig. S19** Proximity-dependent labeling of *Nicotiana benthamiana* proteins by NIMIN1–TurboID does not reveal NPR1.

**Fig. S20** Peptides of the NIMIN1 bait identified by NIMIN1–TurboID mass spectrometry in *Nicotiana benthamiana*.

**Fig. S21** Peptides of TOPLESS-RELATED3 identified by NIMIN1–TurboID mass spectrometry in *Nicotiana benthamiana*.

**Fig. S22** Proximity-dependent labeling of *Nicotiana tabacum* proteins by NIMIN1–TurboID.

**Fig. S23** Peptides of NIM1-like1 (NPR3-like) and NPR1 identified by NIMIN1–TurboID mass spectrometry in transgenic *Nicotiana tabacum* plants.

**Fig. S24** The N-terminal TOPLESS fragment, TPL(1-333), induces cell death in *Nicotiana benthamiana*.

**Table S1** Domain structures of Arabidopsis NIMIN proteins and mutants used in this work.

**Table S2** Proteins identified by mass spectrometry with NIMIN1–TurboID and TurboID–NLS in agroinfiltrated *Nicotiana benthamiana* plants displaying an approximately even distribution in all samples.

**Table S3** Proteins identified by mass spectrometry with NIMIN1–TurboID and TurboID–NLS in transgenic *Nicotiana tabacum* plants displaying an approximately even distribution in all samples.

**Table S4** Primers used for gene construction.

**Methods S1** Detailed description of methods.

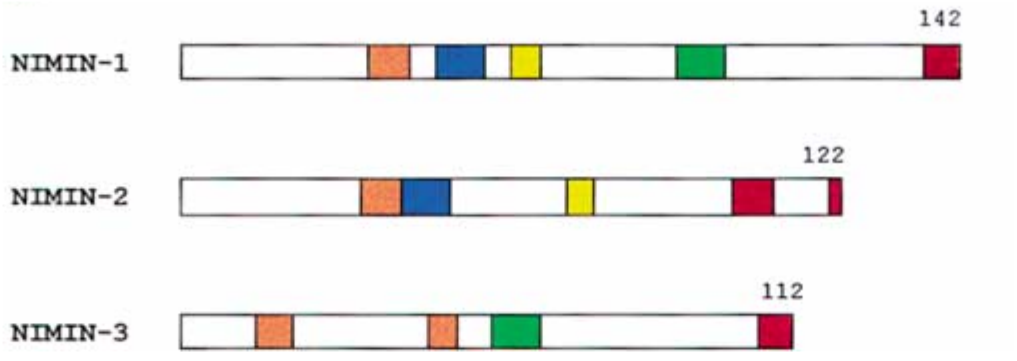


**Fig. S1** Domain structure of Arabidopsis NIMIN1 as deduced from alignment of NIMIN1, NIMIN2 and NIMIN3. The figure was adopted from Weigel *et al.* (2001). The diagram is drawn approximately to scale. Colored boxes denote conserved amino acid motifs. Nuclear localization signal (NLS, yellow); motif mediating NIMIN1 binding to the N1/N2BD in the SA sensitive C-terminus of NPR1 (blue); a second NPR1 interaction motif not linked to the SA response (EDF motif, green); EAR motif (red); clusters of acidic amino acids (orange).

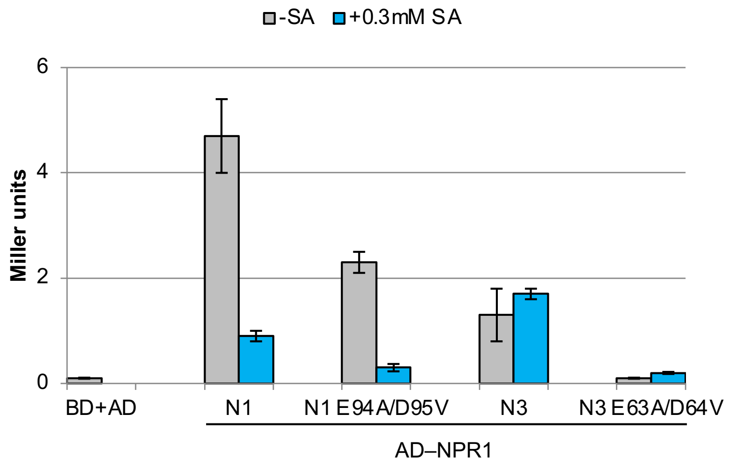


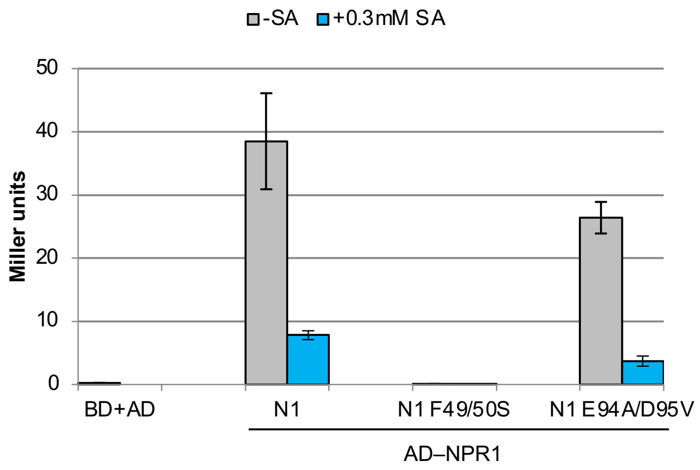


**Fig. S2** Interaction of Arabidopsis NPR1 with NIMIN1 proteins harboring mutations in the two NPR1 binding motifs. Interactions were determined with wild-type NIMIN1 and N3 and with mutants in the NPR1-binding motifs in absence and presence of salicylic acid (SA) in quantitative Y2H assays.

**(a)**

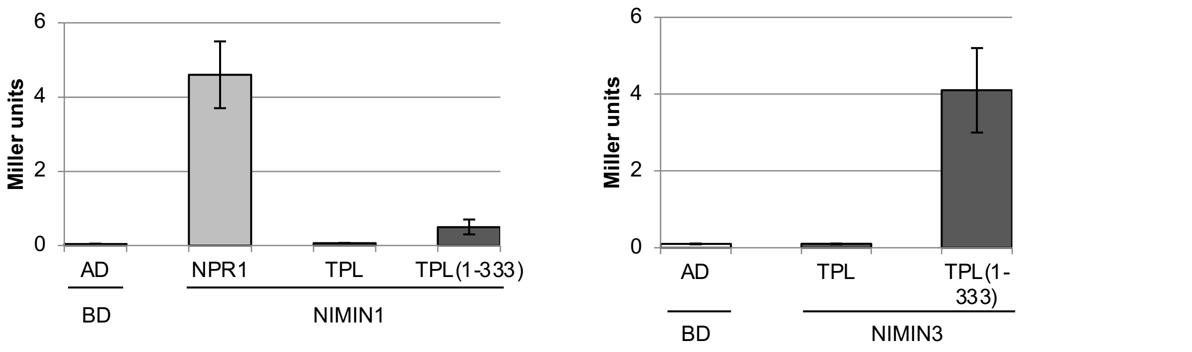


**(b)**

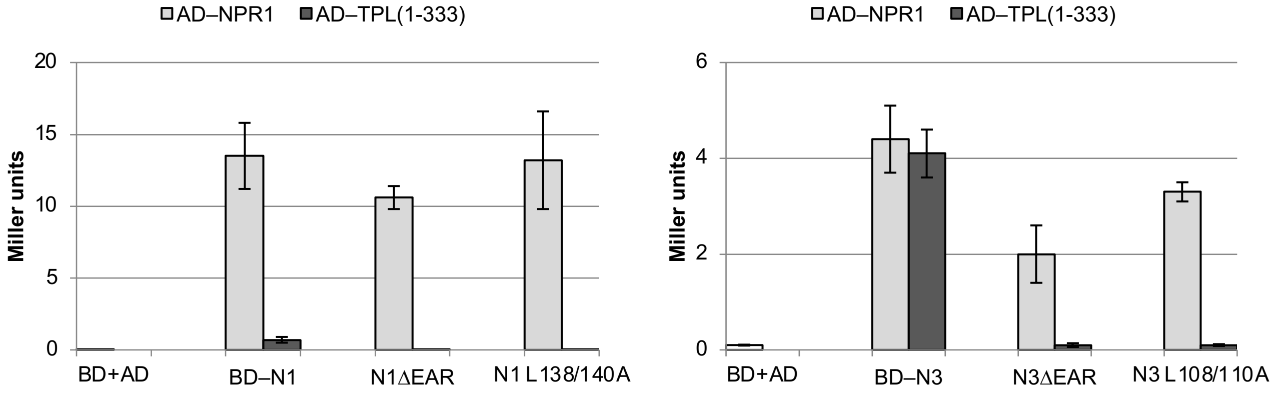


**Fig. S3** Interaction of NIMIN1 and NIMIN3 with Arabidopsis TOPLESS. Interactions were determined in quantitative Y2H assays. Interactions with AtNPR1 served as positive controls. (a) Interaction of NIMIN1 and N3 with full-length TPL and TPL(1-333). (b) Interaction of NIMIN1 and N3 mutants in the EAR motif with TPL(1-333) and NPR1.

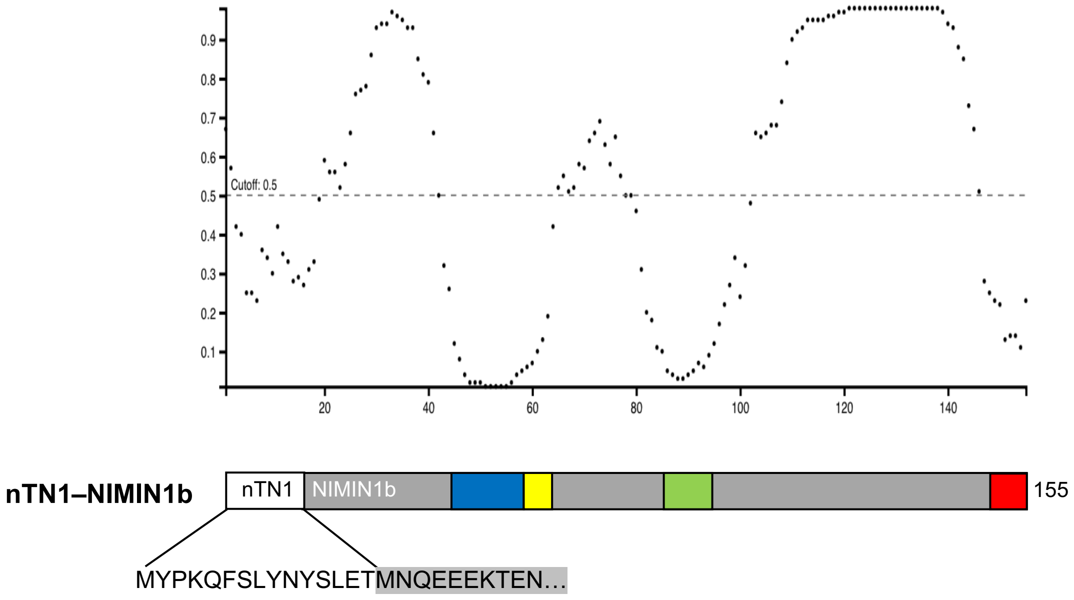


**Fig. S4** Domain structure and disorder profile of a chimeric NIMIN1b protein, nTN1–NIMIN1b, harboring the N-terminal region of NIMIN1. The diagram is drawn approximately to scale. Colored boxes denote conserved structural motifs as given in Figures 1 and S1. White, 15-amino acid long stretch from the N-terminus of NIMIN1 (nTN1).

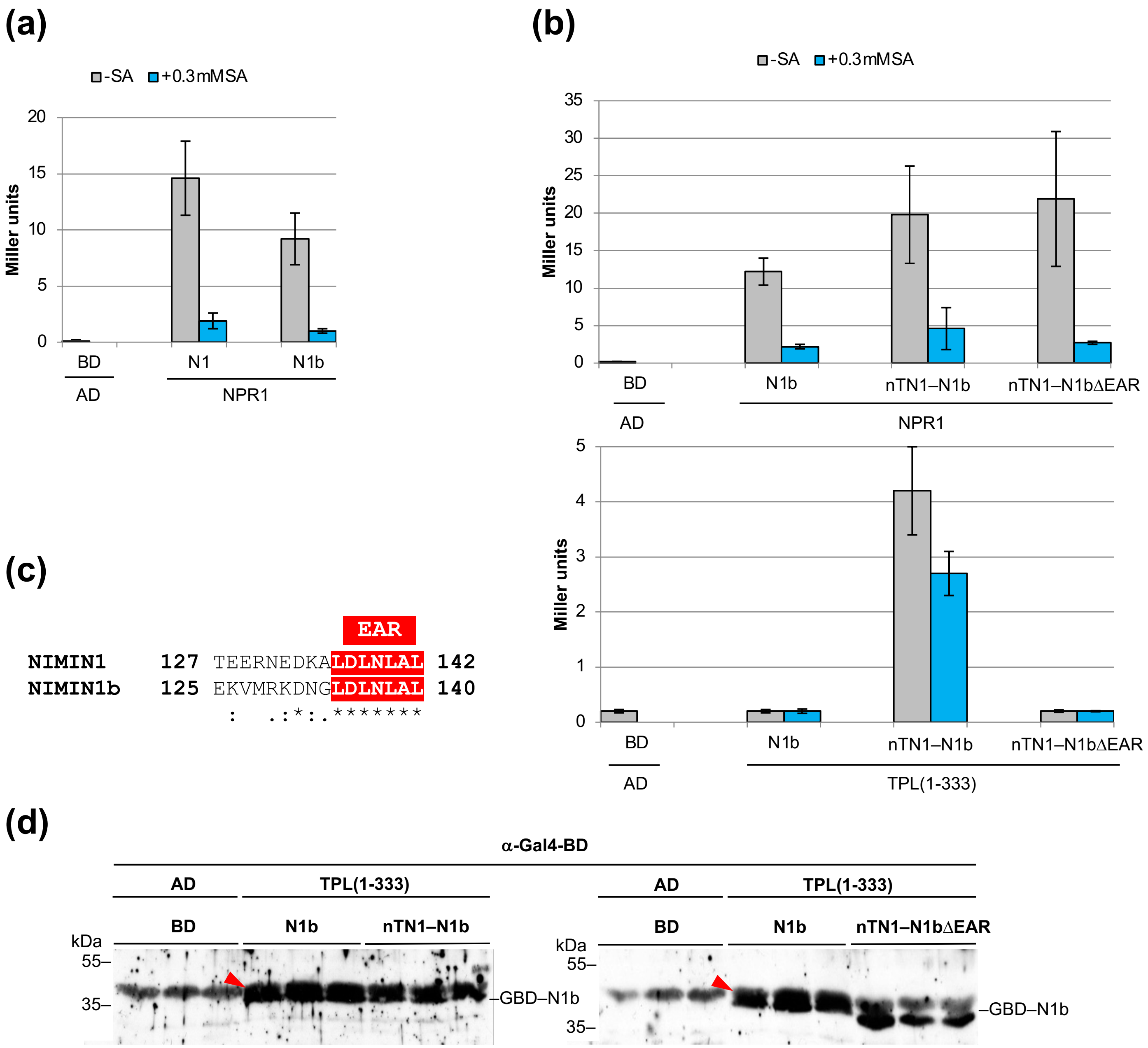


**Fig. S5** Interaction of chimeric nTN1–NIMIN1b with Arabidopsis NPR1 and TPL(1-333). Interactions were determined with wild-type proteins and mutants as indicated in absence and presence of salicylic acid (SA) in quantitative Y2H assays. (a) Interaction of NIMIN1b and NIMIN1 with NPR1. (b) Interaction of NIMIN1b, nTN1–NIMIN1b and a mutant in the EAR motif with NPR1 and TPL(1-333). (c) Alignment of the EAR motifs from NIMIN1 and NIMIN1b. (d) Accumulation of NIMIN1b, nTN1–NIMIN1b and nTN1–NIMIN1bΔEAR fusion proteins in yeast. Protein extracts from three independent colonies for each transformation were analyzed by immunodetection with an antiserum directed against the Gal4 DNA-binding domain (Gal4-BD, GBD). The position of GBD–NIMIN1b in the gel is indicated. The red arrowhead denotes an unspecific band detected in all lanes indicative for equal protein loading.

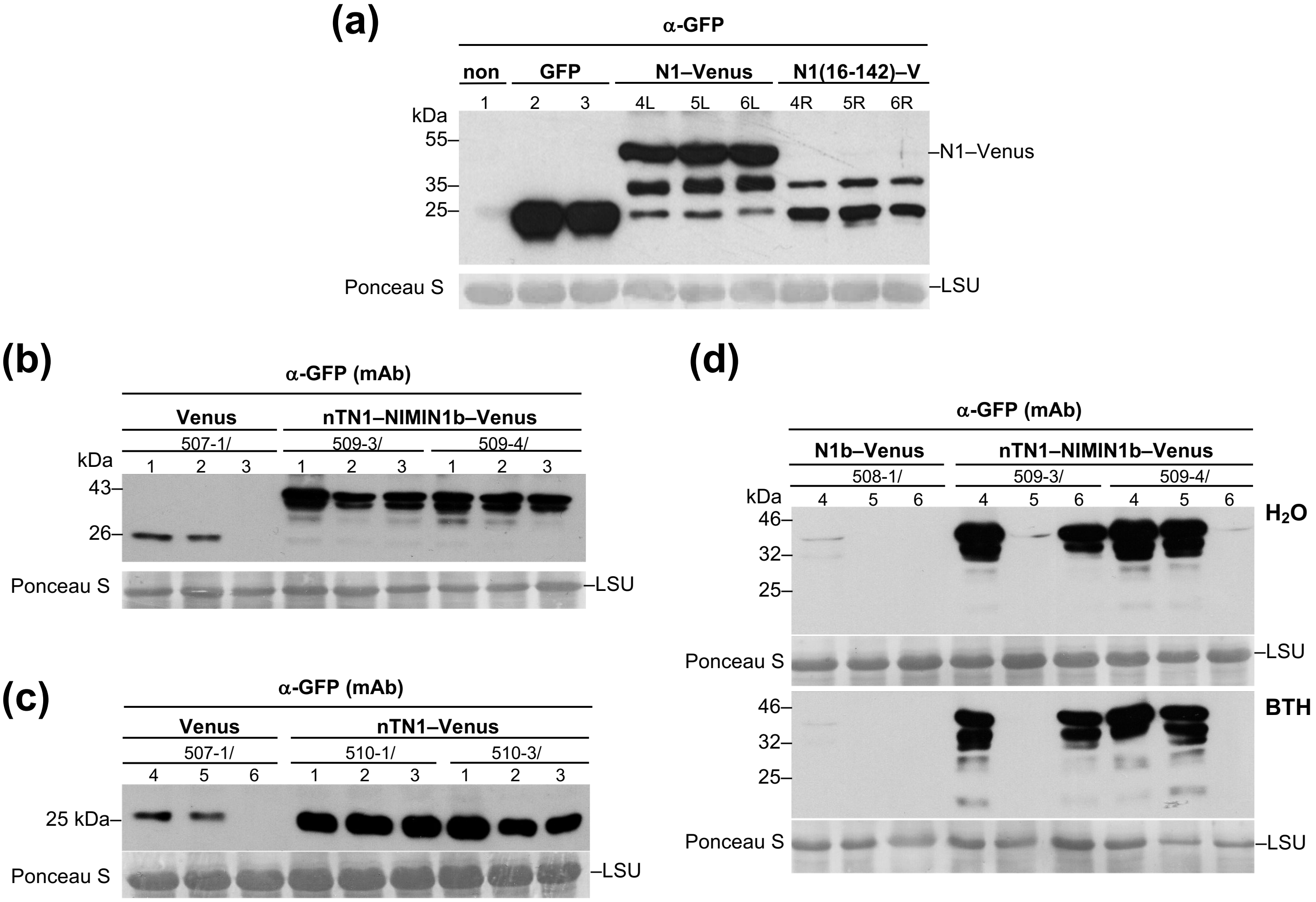


**Fig. S6** The N-terminal *NIMIN1* sequence *(nTN1)* reinforces protein accumulation *in planta*. Immunodetection of VENUS fusion proteins in leaf extracts was with a polyclonal antiserum (a) or with a monoclonal antibody (mAb) directed against GFP (b to d). Staining of the large subunit of RUBISCO (LSU) with Ponceau S demonstrates loading of the nitrocellulose filters. (a) Accumulation of NIMIN1(16-142)–VENUS in *N. benthamiana*. Accumulation of N1(16-142)–VENUS is compared to accumulation of NIMIN1–VENUS. For direct comparison of protein accumulation from the two gene constructs, Agrobacterium strains were infiltrated in the left (L) and right (R) halves of the same leaves as indicated. Three plants were infiltrated in parallel for each construct. (b) Accumulation of nTN1–NIMIN1b–VENUS in leaves of transgenic tobacco plants. Accumulation of nTN1–N1b–VENUS is compared to accumulation of VENUS. Three T1 plants were analyzed in parallel for each transgenic line. (c) Accumulation of nTN1–VENUS in leaves of transgenic tobacco plants. Accumulation of nTN1–VENUS is compared to accumulation of VENUS. (d) Accumulation of nTN1–NIMIN1b–VENUS in BTH-treated leaves of transgenic tobacco plants. Accumulation of nTN1–N1b–VENUS is compared to accumulation of N1b–VENUS in $H_2O$ and BTH-treated leaf tissue. Immunoblots were exposed to the same X-ray film.

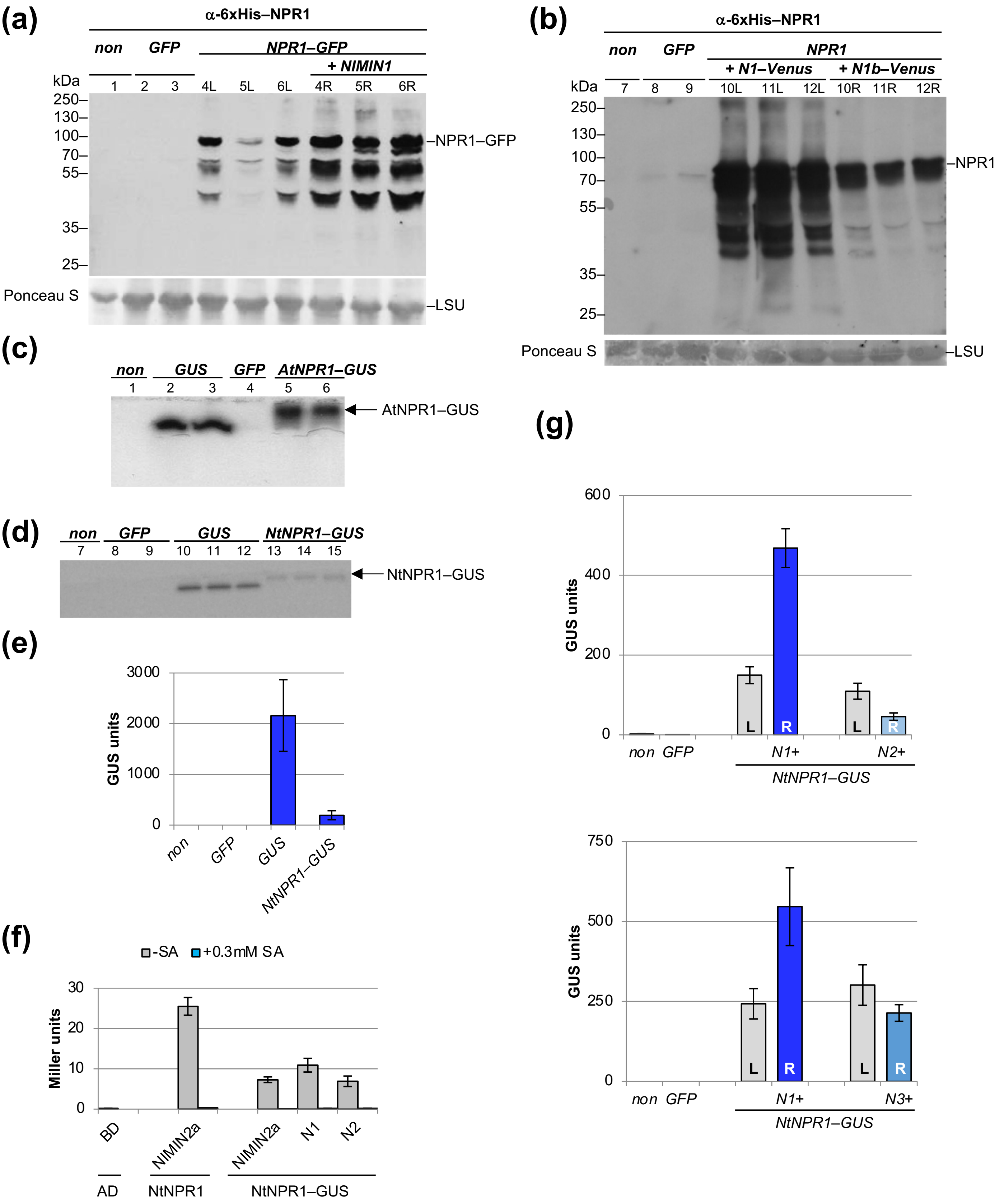
(a)
α-6xHis–NPR1
non
GFP
NPR1–GFP
+ NIMIN1
kDa
1 2 3 4L 5L 6L 4R 5R 6R
250–
130–
100–
70–
55–
35–
25–
–NPR1–GFP
Ponceau S
–LSU
(b)
α-6xHis–NPR1
non
GFP
NPR1
+ N1–Venus
+ N1b–Venus
kDa
7 8 9 10L 11L 12L 10R 11R 12R
250–
130–
100–
70–
55–
35–
25–
–NPR1
Ponceau S
–LSU
(c)
non GUS GFP AtNPR1–GUS
1 2 3 4 5 6
AtNPR1–GUS
(d)
non GFP GUS NtNPR1–GUS
7 8 9 10 11 12 13 14 15
NtNPR1–GUS
(e)
GUS units
3000
2000
1000
0
non
GFP
GUS
NtNPR1–GUS
(f)
-SA
+0.3mM SA
Miller units
30
20
10
0
BD
NIMIN2a
NIMIN2a
N1
N2
AD
NtNPR1
NtNPR1–GUS
(g)
GUS units
600
400
200
0
L R L R
non GFP N1+ N2+
NtNPR1–GUS
GUS units
750
500
250
0
L R L R
non GFP N1+ N3+
NtNPR1–GUS

**Fig. S7** *NIMIN1* co-expression strengthens accumulation of NPR1 *in planta*. (a) Accumulation of NPR1–GFP with and without co-expression of *NIMIN1* in *Nicotiana benthamiana*. For direct comparison of protein accumulation, mixtures of Agrobacterium strains were infiltrated in the left (L) and right (R) halves of the same leaves as indicated. Three plants were infiltrated in parallel. Immunodetection in leaf extracts was with a polyclonal antiserum directed against 6xHis–AtNPR1. Staining of the large subunit of RUBISCO (LSU) with Ponceau S demonstrates loading of the nitrocellulose filter. (b) Accumulation of NPR1 with co-expression of *NIMIN1–VENUS* or *NIMIN1b–VENUS* in *Nicotiana benthamiana*. (c) In-gel activity assay for NPR1–GUS. Crude leaf extracts from different plants infiltrated with Agrobacteria as indicated were separated in a non-denaturing gel and tested for GUS enzyme activity. (d) In-gel activity assay for NtNPR1–GUS. (e) Enzyme activity of NtNPR1–GUS fusion protein. Crude leaf extracts from three plants each infiltrated with Agrobacteria as indicated were tested for GUS activity. Mean activities plus and minus standard deviation are shown. (f) Interaction of NtNPR1–GUS with NIMIN proteins. Interactions were determined in absence and presence of salicylic acid (SA) in quantitative Y2H assays. Interaction of GBD–NtN2a with GAD–NtNPR1 served as positive control. (g) Reporter activity of NtNPR1–GUS fusion protein with and without co-expression of *NIMIN1, NIMIN2* or *NIMIN3* in *N. benthamiana*. For direct comparison of GUS enzyme activities, mixtures of Agrobacterium strains were infiltrated in the left (L) and right (R) halves of the same leaves as indicated. Three plants were infiltrated in parallel.

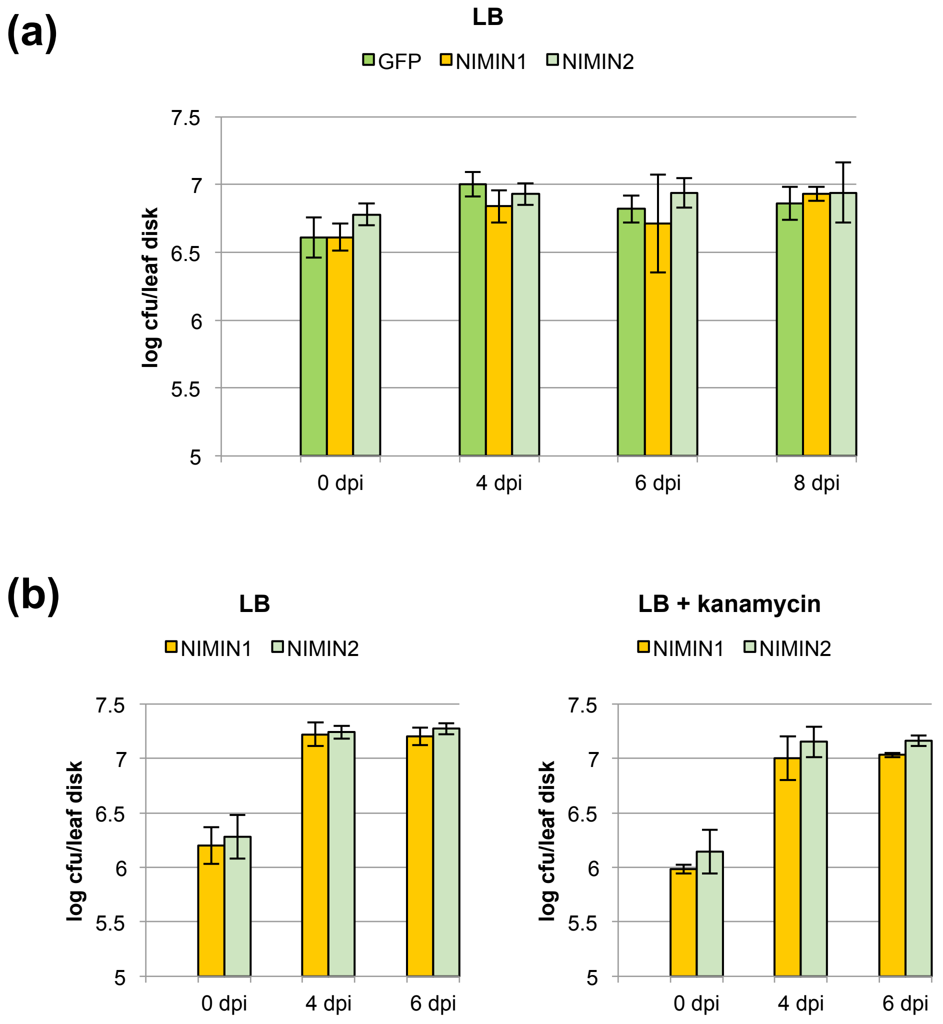


**Fig. S8** *NIMIN1* overexpression in *Nicotiana benthamiana* does not affect microbial growth. Leaf disks from three different plants and two different leaves for each plant were extracted at indicated times after agroinfiltration, and serial dilutions were plated on medium as indicated. Bacterial counts are given as log colony-forming units per leaf disk ± standard deviation (SD). (a) Microbial growth in plants infiltrated with Agrobacterium strains harboring *35S:GFP*, *35S:NIMIN1* or *35S:NIMIN2.* Dilutions of leaf extracts were plated on LB medium. (b) Microbial growth in plants infiltrated with Agrobacterium strains harboring *35S:NIMIN1* or *35S:NIMIN2.* Agrobacterium strains were infiltrated in the left and right halves of the same leaves, respectively. Dilutions of leaf extracts were plated on LB medium or on LB medium with kanamycin.

**(a)**

**(b)**

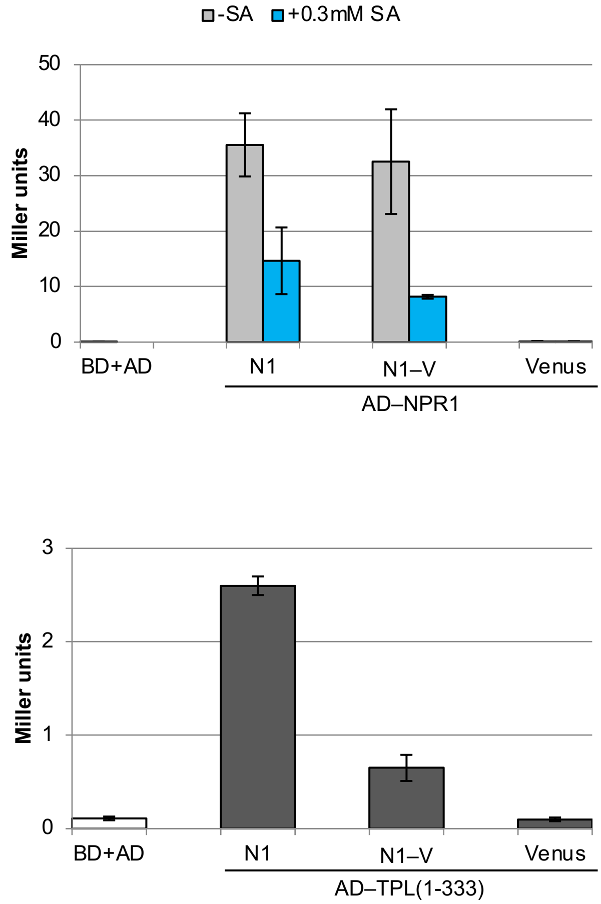


**Fig. S9** Interaction of NIMIN1–VENUS with NPR1 and TOPLESS. Interactions were determined in quantitative Y2H assays. Interactions with NIMIN1 served as positive controls and interactions with VENUS as negative controls. (a) Interaction of NIMIN1–VENUS with Arabidopsis NPR1 in absence and presence of salicylic acid. (b) Interaction of NIMIN1–VENUS with Arabidopsis TPL(1-333).

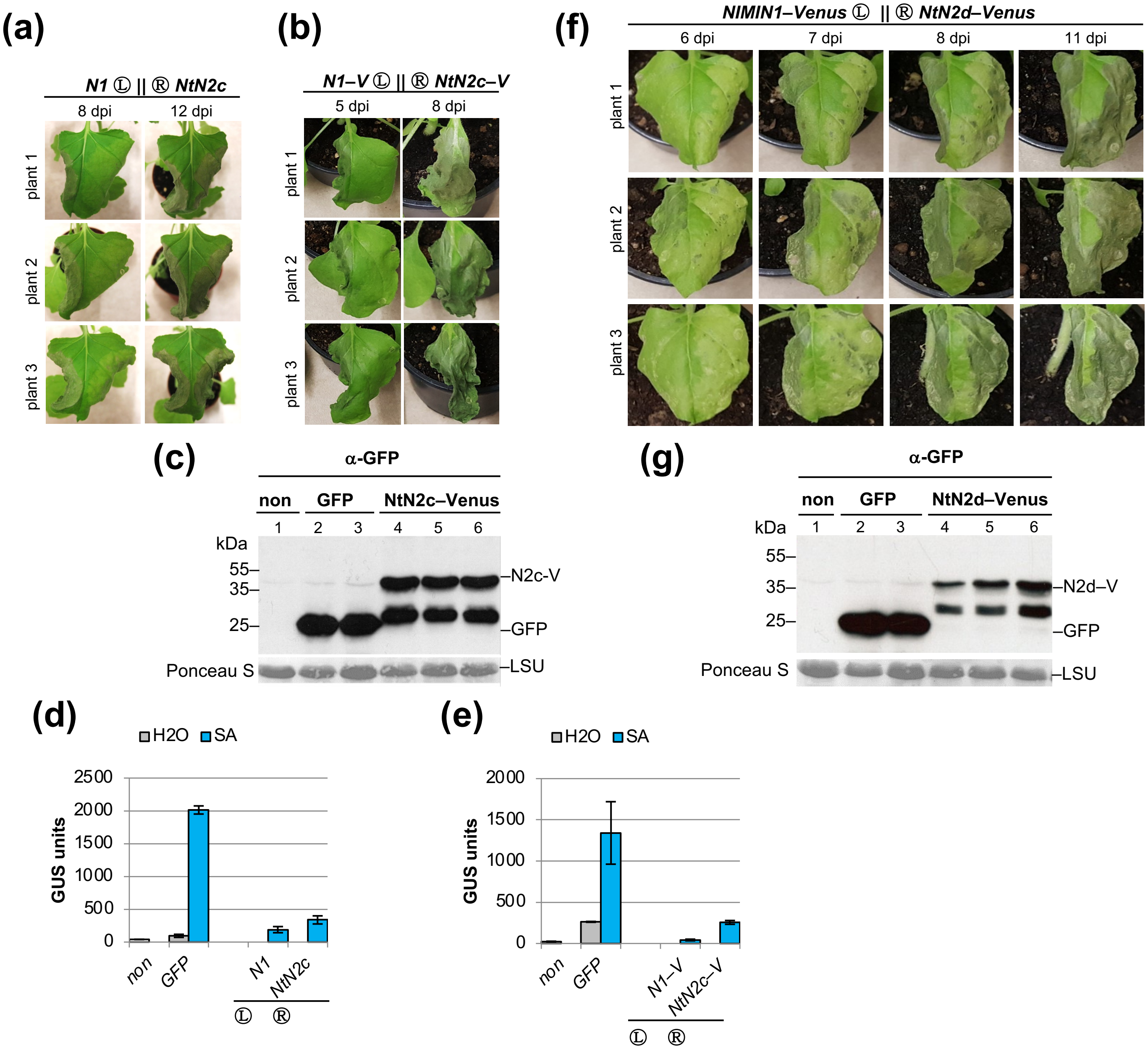


**Fig. S10** Overexpression of tobacco *NIMIN2c* and *NIMIN2d* promotes emergence of cell death and *PR-1a* gene suppression *in planta*. (a) Phenotype of *Nicotiana benthamiana* leaves overexpressing tobacco *NIMIN2c*. Agrobacteria were infiltrated in the right halves of leaves. Effects of Nt*NIMIN2c* overexpression are compared to overexpression of At*NIMIN1*. (b) Phenotype of *Nicotiana benthamiana* leaves overexpressing Nt*NIMIN2c–VENUS*. Agrobacteria were infiltrated in the right halves of leaves. Effects of Nt*NIMIN2c–VENUS* overexpression are compared to overexpression of At*NIMIN1–VENUS*. (c) Accumulation of NtNIMIN2c–VENUS in *Nicotiana benthamiana*. Three plants were agroinfiltrated in parallel. Immunodetection of VENUS fusion protein in leaf extracts was with a polyclonal antiserum directed against GFP. Staining of the large subunit of RUBISCO (LSU) with Ponceau S demonstrates loading of the nitrocellulose filter. (d) Salicylic acid-mediated *PR-1a* gene induction is suppressed in leaf tissue overexpressing Nt*NIMIN2c*. Agrobacteria were infiltrated in the right (R) leaf halves of *N. benthamiana* plants harboring a *-1533PR1a*$_{Pro}$*:GUS* transgene. Three plants were infiltrated in parallel. GUS reporter activity was determined after floating of disks excised from infiltrated leaf areas on water or 1 mM SA. GUS activity is compared to enzyme activities in *NIMIN1* and *GFP* overexpressing tissue. (e) Salicylic acid-mediated *PR-1a* gene induction is suppressed in leaf tissue overexpressing Nt*NIMIN2c–VENUS*. Agrobacteria were infiltrated in the right (R) leaf halves of *N. benthamiana* plants harboring a *-1533PR1a*$_{Pro}$*:GUS* transgene. GUS activity is compared to enzyme activities in *NIMIN1–VENUS* and *GFP* overexpressing tissue. (f) Phenotype of *Nicotiana benthamiana* leaves overexpressing Nt*NIMIN2d–VENUS*. Agrobacteria were infiltrated in the right halves of leaves. Effects of Nt*NIMIN2d–VENUS* overexpression are compared to overexpression of At*NIMIN1–VENUS*. (g) Accumulation of NtNIMIN2d–VENUS in *Nicotiana benthamiana*. Three plants were agroinfiltrated in parallel.

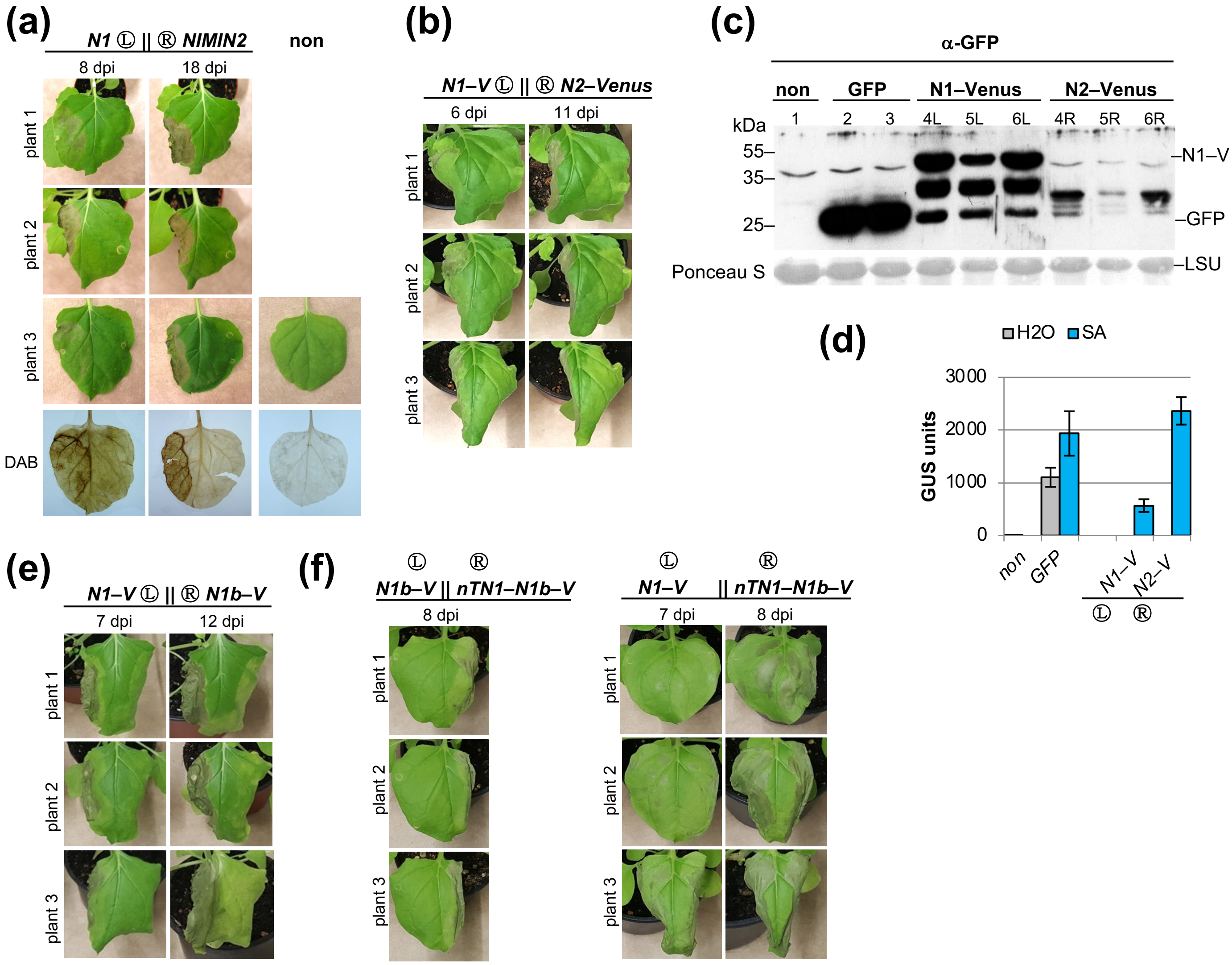


**Fig. S11** Overexpression of *NIMIN2* or *NIMIN1b* does not promote emergence of cell death *in planta*. (a) Phenotype of *Nicotiana benthamiana* leaves and production of $H_2O_2$ by overexpression of *NIMIN2*. Agrobacteria were infiltrated in the right leaf halves. Phenotypes and $H_2O_2$ accumulation visualized by DAB staining were recorded at different days post infiltration (dpi). Effects of *NIMIN2* overexpression are compared to overexpression of *NIMIN1*. (b) Phenotype of *Nicotiana benthamiana* leaves overexpressing *NIMIN2–VENUS*. Agrobacteria were infiltrated in the right leaf halves. Phenotypes were recorded at different days post infiltration (dpi). Effects of *N2–VENUS* overexpression are compared to overexpression of *N1–VENUS*. (c) Accumulation of NIMIN2–VENUS in *Nicotiana benthamiana*. Accumulation of N2–VENUS is compared to accumulation of N1–VENUS. For direct comparison of protein accumulation, Agrobacterium strains were infiltrated in the left (L) and right (R) halves of the same leaves as indicated. Three plants were infiltrated in parallel. Immunodetection of VENUS fusion protein in leaf extracts was with a polyclonal antiserum directed against GFP. Staining of the large subunit of RUBISCO (LSU) with Ponceau S demonstrates loading of the nitrocellulose filter. (d) Salicylic acid-mediated *PR-1a* gene induction is not suppressed in leaf tissue overexpressing *NIMIN2–VENUS*. Agrobacteria were infiltrated in the right (R) leaf halves of *N. benthamiana* plants harboring a *-1533PR1a*$_{Pro}$*:GUS* transgene. Three plants were infiltrated in parallel. GUS reporter activity was determined after floating of disks excised from infiltrated leaf areas on water or 1 mM SA. GUS activity is compared to enzyme activities in *NIMIN1–VENUS* and *GFP* overexpressing tissue. (e) Phenotype of *Nicotiana benthamiana* leaves overexpressing *NIMIN1b–VENUS*. Agrobacteria were infiltrated in the right (R) leaf halves. Phenotypes were recorded at different days post infiltration (dpi). Effects of *N1b–VENUS* overexpression are compared to overexpression of *N1–VENUS*. (f) Phenotype of *Nicotiana benthamiana* leaves overexpressing *nTN1–NIMIN1b–VENUS*. Agrobacteria were infiltrated in the right leaf halves. Phenotypes were recorded at different days post infiltration (dpi). Effects of *nTN1–N1b–VENUS* overexpression are compared to overexpression of *N1b–VENUS* and *N1–VENUS*.

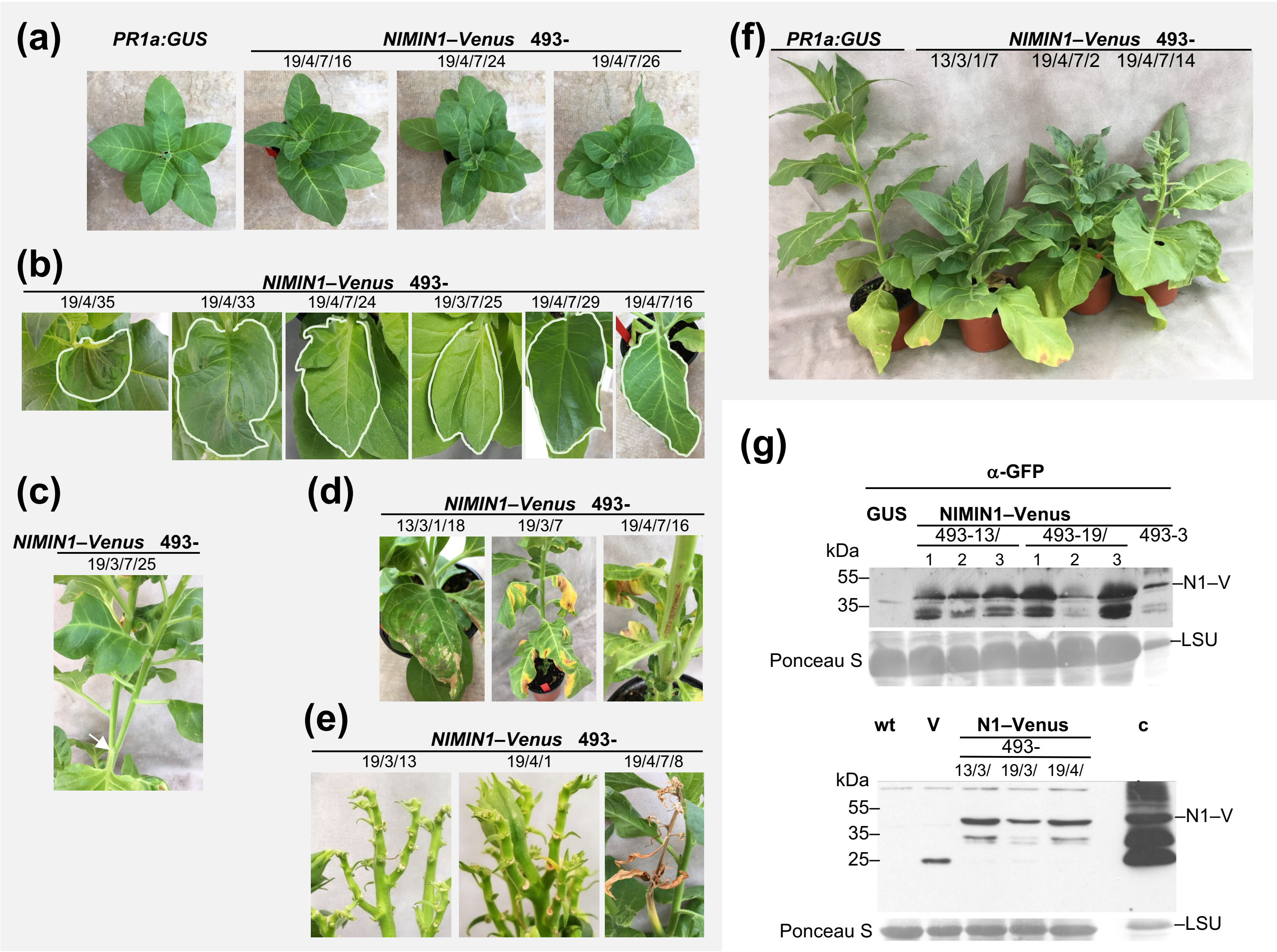


**Fig. S12** Phenotypes of *NIMIN1–VENUS* overexpressing transgenic tobacco plants. (a) Asymmetric statures of T3 plants of line 493-19. (b) Malformed leaves of T2 and T3 plants of line 493-19. The contours of leaves are highlighted. (c) Furcated shoot of a T3 plant of line 493-19. The arrow points to the furcation of the stem. (d) Cell death on leaves and shoots of T2 and T3 plants of lines 493-13 and 493-19. (e) Cell death on pedicels and shoot tips of T2 and T3 plants of line 493-19. (f) Stunted growth of T3 plants of lines 493-13 and 493-19. (g) Accumulation of NIMIN1–VENUS in transgenic tobacco plants. Upper panel: leaf extracts from a $PR1a_{Pro}$*:GUS* plant, from primary transformant 493-3 and from different T1 plants of lines 493-13 and 493-19. Lower panel: extracts of 10 seedlings each from wild-type (*N. tabacum* cv. Samsun NN), from a line expressing *35S:VENUS* and from T2 seedlings of lines 493-13 and 493-19. As a positive control, a leaf extract from *N. benthamiana* infiltrated with *35S:NIMIN1–VENUS* Agrobacteria was loaded. Immunodetection of NIMIN1–VENUS fusion protein was with a polyclonal antiserum directed against GFP. Staining of the large subunit of RUBISCO (LSU) with Ponceau S demonstrates loading of the nitrocellulose filter.

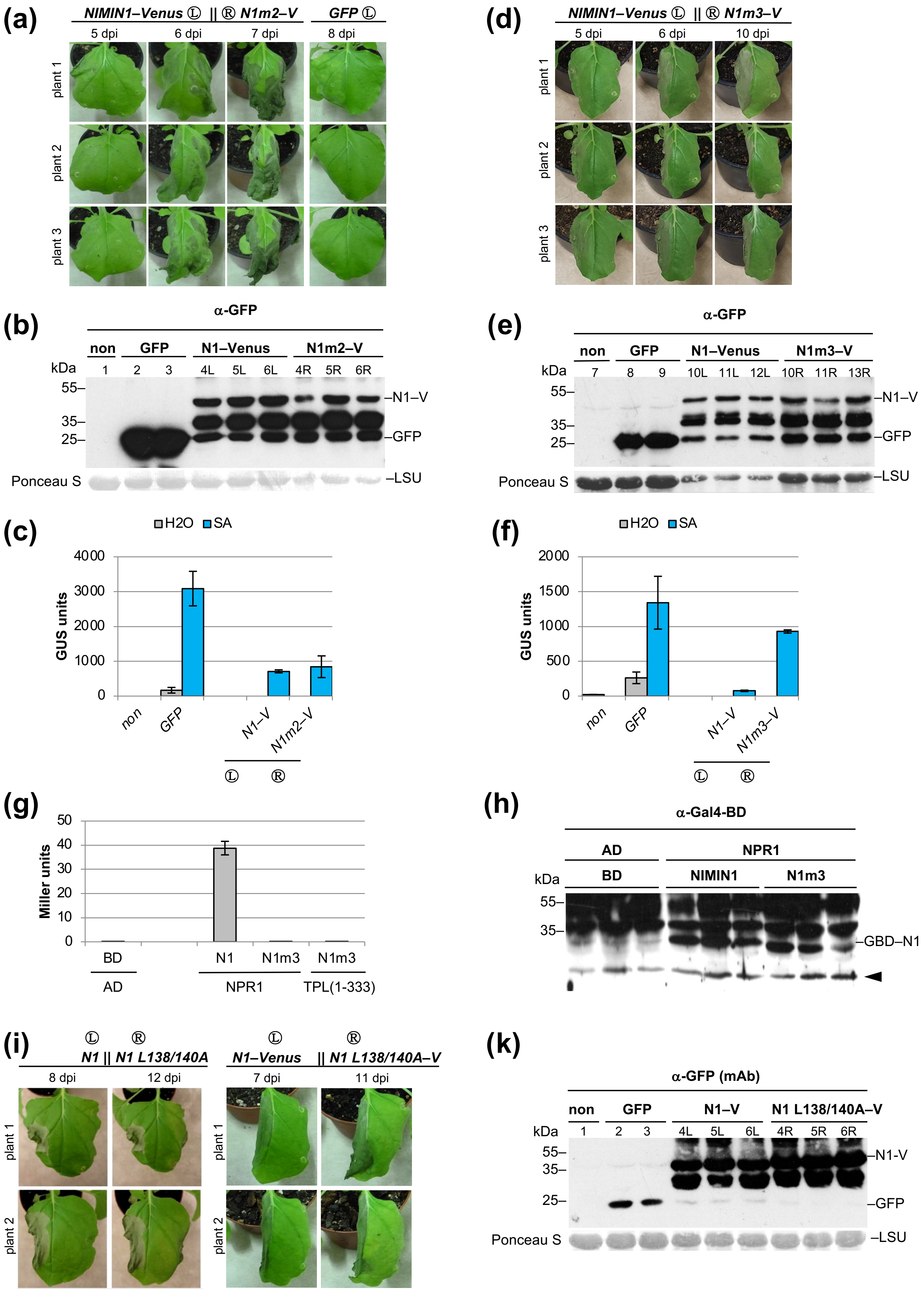
(a)
NIMIN1–Venus Ⓛ || Ⓡ N1m2–V
GFP Ⓛ
5 dpi
6 dpi
7 dpi
8 dpi
plant 1
plant 2
plant 3
(b)
α-GFP
non
GFP
N1–Venus
N1m2–V
kDa
1 2 3 4L 5L 6L 4R 5R 6R
55–
35–
25–
–N1–V
–GFP
Ponceau S
–LSU
(c)
H2O
SA
4000
3000
2000
1000
0
GUS units
non
GFP
N1–V
N1m2–V
Ⓛ Ⓡ
(d)
NIMIN1–Venus Ⓛ || Ⓡ N1m3–V
5 dpi
6 dpi
10 dpi
plant 1
plant 2
plant 3
(e)
α-GFP
non
GFP
N1–Venus
N1m3–V
kDa
7 8 9 10L 11L 12L 10R 11R 13R
55–
35–
25–
–N1–V
–GFP
Ponceau S
–LSU
(f)
H2O
SA
2000
1500
1000
500
0
GUS units
non
GFP
N1–V
N1m3–V
Ⓛ Ⓡ
(g)
50
40
30
20
10
0
Miller units
BD
N1
N1m3
N1m3
AD
NPR1
TPL(1-333)
(h)
α-Gal4-BD
AD
NPR1
BD
NIMIN1
N1m3
kDa
55–
35–
–GBD–N1
(i)
Ⓛ Ⓡ
N1 || N1 L138/140A
N1–Venus || N1 L138/140A–V
8 dpi
12 dpi
7 dpi
11 dpi
plant 1
plant 2
(k)
α-GFP (mAb)
non
GFP
N1–V
N1 L138/140A–V
kDa
1 2 3 4L 5L 6L 4R 5R 6R
55–
35–
25–
–N1-V
–GFP
Ponceau S
–LSU

**Fig. S13** The EAR motif is instrumental in NIMIN1-mediated cell death induction. (a,d) Phenotype of *Nicotiana benthamiana* leaves overexpressing *N1m2–VENUS* or *N1m3–VENUS*. Agrobacteria were infiltrated in the right halves of leaves. Effects of *N1m2–VENUS* (a) and *N1m3–VENUS* (d) overexpression are compared to overexpression of *NIMIN1–VENUS*. (b,e) Accumulation of N1m2–VENUS and N1m3–VENUS in *Nicotiana benthamiana*. Three plants were agroinfiltrated in parallel. Immunodetection of VENUS fusion proteins in leaf extracts was with a polyclonal antiserum directed against GFP. Staining of the large subunit of RUBISCO (LSU) with Ponceau S demonstrates loading of the nitrocellulose filter. (c,f) Salicylic acid-mediated *PR-1a* gene induction is suppressed in leaf tissue overexpressing *N1m2–VENUS* (c) and not suppressed in leaf tissue overexpressing *N1m3–VENUS* (f). Agrobacteria were infiltrated in the right (R) leaf halves of *N. benthamiana* plants harboring a $\textit{-1533PR1a}_{\textit{Pro}}$*:GUS* transgene. Three plants were infiltrated in parallel. GUS reporter activity was determined after floating of disks excised from infiltrated leaf areas on water or 1 mM SA. GUS activity is compared to enzyme activities in *NIMIN1–VENUS* and *GFP* overexpressing tissue. (g) Y2H interaction of NIMIN1m3 with NPR1 and TPL(1-333). (h) Accumulation of GBD–NIMIN1m3 in yeast. Protein extracts from three independent colonies for each transformation were analyzed by immunodetection with an antiserum directed against the Gal4 DNA-binding domain (Gal4-BD, GBD). The position of GBD–NIMIN1 in the gel is indicated. The arrowhead points to an unspecific band detected in all lanes indicating equal protein loading. (i) Phenotype of *Nicotiana benthamiana* leaves overexpressing *NIMIN1 L138/140A(–VENUS)*. Agrobacteria were infiltrated in the right halves of leaves. Effects are compared to overexpression of *NIMIN1* and *NIMIN1–VENUS*, respectively. (k) Accumulation of N1 L138/140A–VENUS in *Nicotiana benthamiana*. Three plants were agroinfiltrated in parallel. Immunodetection of VENUS fusion proteins in leaf extracts was with a monoclonal antibody directed against GFP. Staining of the large subunit of RUBISCO (LSU) with Ponceau S demonstrates loading of the nitrocellulose filter.

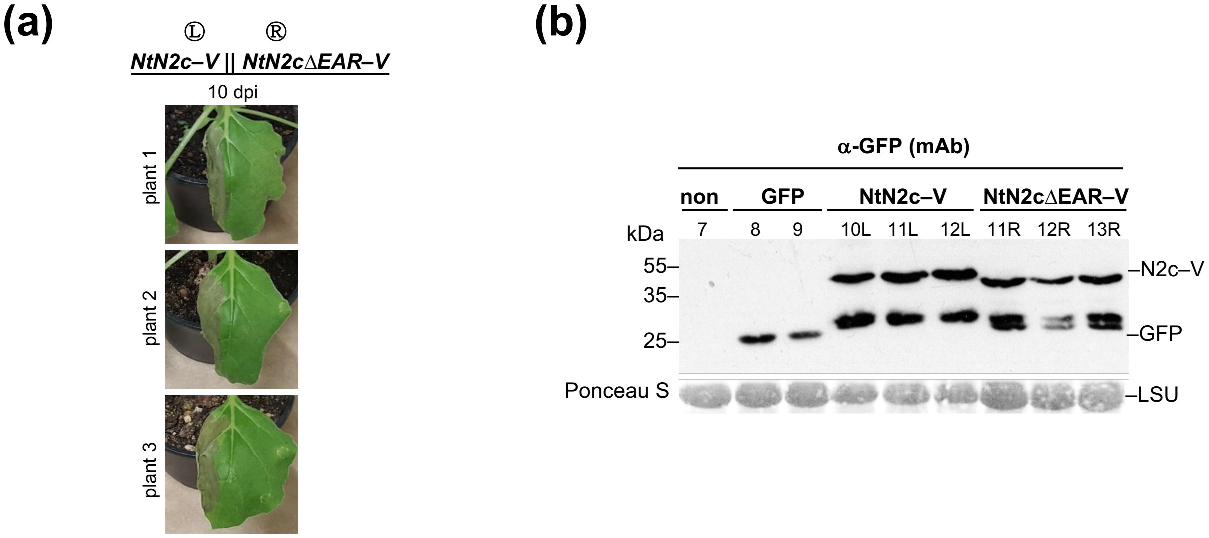


**Fig. S14** The EAR motif is instrumental in NtNIMIN2c-mediated cell death induction. (a) Phenotype of *Nicotiana benthamiana* leaves overexpressing *NtNIMIN2cΔEAR–VENUS*. Agrobacteria were infiltrated in the right halves of leaves. Effects are compared to overexpression of *NtNIMIN2c–VENUS*. (b) Accumulation of NtNIMIN2cΔEAR–VENUS in *Nicotiana benthamiana*. Three plants were agroinfiltrated in parallel. Immunodetection of VENUS fusion proteins in leaf extracts was with a monoclonal antibody directed against GFP. Staining of the large subunit of RUBISCO (LSU) with Ponceau S demonstrates loading of the nitrocellulose filter.

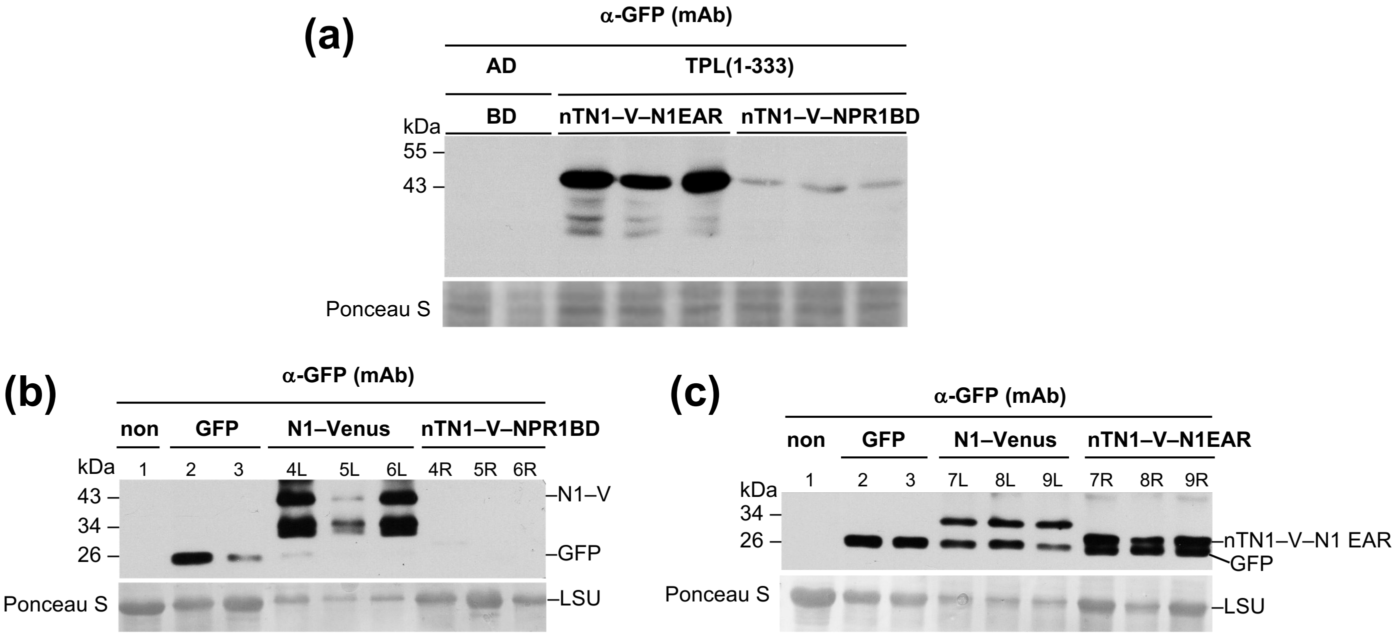


**Fig. S15** Expression of *nTN1–VENUS–NPR1BD* and *nTN1–VENUS–N1EAR* fusion genes in yeast and in *Nicotiana benthamiana*. Protein extracts were analyzed by immunodetection with a monoclonal antibody (mAb) directed against GFP. Staining of the nitrocellulose filters with Ponceau S demonstrates equal protein loading. (a) Expression in yeast. Proteins were extracted from three independent colonies for each transformation. (b) Expression of *nTN1–VENUS–NPR1BD* in *Nicotiana benthamiana*. Expression of *nTN1–VENUS–NPR1BD* is compared to expression of *N1–VENUS*. Three plants were agroinfiltrated in parallel. (c) Expression of *nTN1–VENUS–N1EAR* in *N. benthamiana*. Expression of *nTN1–VENUS–N1EAR* is compared to expression of *N1–VENUS*. Three plants were agroinfiltrated in parallel.

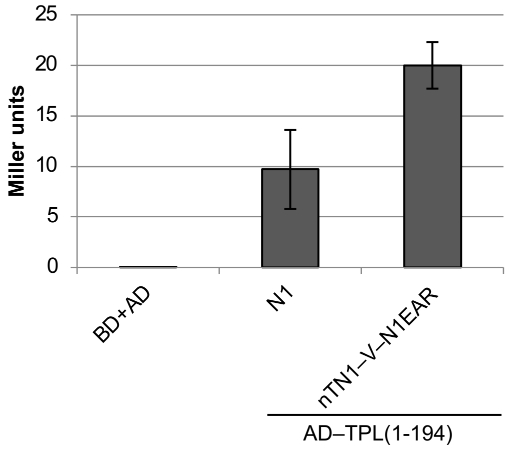


**Fig. S16** Yeast two-hybrid interaction of nTN1–VENUS–N1EAR with TPL(1-194). Interaction of nTN1–VENUS–N1EAR is compared to interaction of NIMIN1.

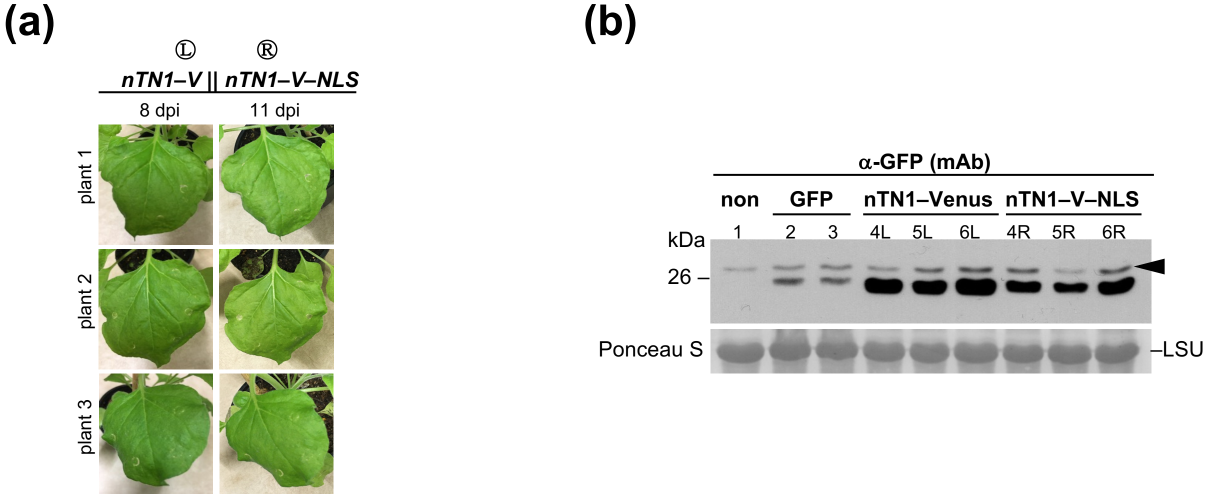


**Fig. S17** Overexpression of *nTN1–VENUS* or *nTN1–VENUS–NLS* does not induce cell death in *Nicotiana benthamiana*. (a) Phenotype of *Nicotiana benthamiana* leaves overexpressing *nTN1–VENUS* or *nTN1–VENUS–NLS*. Three plants were agroinfiltrated in parallel. Agrobacteria were infiltrated in the left and right halves of leaves, respectively. (b) Accumulation of nTN1–VENUS and nTN1–VENUS–NLS fusion proteins in *Nicotiana benthamiana*. Immunodetection of VENUS fusion proteins in leaf extracts was with a monoclonal antibody (mAb) directed against GFP. Staining of the large subunit of RUBISCO (LSU) with Ponceau S and detection of an unspecic band (indicated by the arrowhead) demonstrate loading of the nitrocellulose filter.

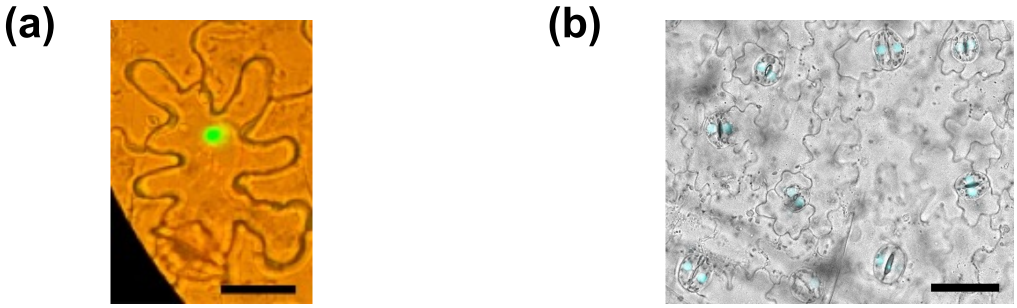


**Fig. S18** Subcellular localization of NIMIN1–TurboID fusion protein in *Nicotiana benthamiana* and *N. tabacum*. Overlay pictures representing typical results are shown**.** (a) Epifluorescence image of an epidermal cell of *N. benthamiana* leaves transformed with *UBQ10*$_{Pro}$*:Turbo–NIMIN1–YFP*. Scale bar, 50 µm. (b) Digital microscope image of the leaf epidermis of *N. tabacum* stably transformed with *Ubq10*$_{Pro}$*:Turbo–NIMIN1–YFP* (line 165H-5). YFP channel in light blue. Scale bar, 100 µm.

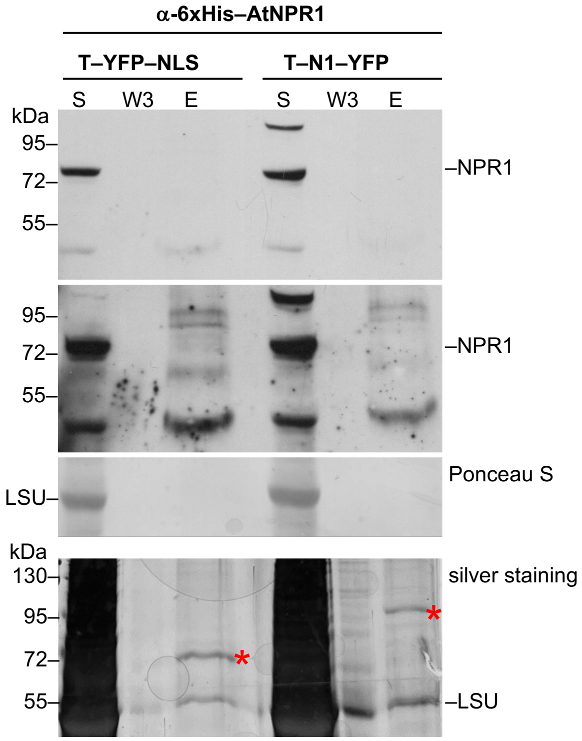


**Fig. S19** Proximity-dependent labeling of *Nicotiana benthamiana* proteins by NIMIN1–TurboID does not reveal NPR1. Biotinylated proteins were purified from crude extracts of agroinfiltrated *N. benthamiana* leaf tissue overexpressing TurboID constructs with Strep-Tactin-coated beads. Aliquots from the supernatant (S), the third wash fraction (W3) and the eluate (E) were separated on SDS gels. NPR1 was detected as a 72 kDa band in the crude extracts, but not in the Turbo–N1–YFP eluate, by anti-6xHis–AtNPR1 blotting. Two different exposures of the X-ray film are shown. Staining of the large subunit of RUBISCO (LSU) with Ponceau S demonstrates loading of the nitrocellulose filter. A parallel gel was stained with silver nitrate to illustrate presence of the fusion proteins (marked by red asterisks) in the eluates. The large subunit of RUBISCO (LSU) is also detected in the Strep-Tactin pull-downs.

```
MYPKQFSLYN YSLETMSKDE NVESKETIRV DKRVREDEEE EEEKKIDTFF KLIKHYQEAR  60
KRRREELAEN SGVVRRKSNG GERSGIVVPA FQPEDFSQCR TGLKPPLMFV SDHKEENTKV 120
EQEEDQTEER NEDKALDLNL AL 142
```

**Fig. S20** Peptides of the NIMIN1 bait identified by NIMIN1–TurboID mass spectrometry in *Nicotiana benthamiana*. Peptides identified in one MS run are highlighted in yellow and orange. The protein coverage is 65%.

```
NtTPL      DEVEKYLSGFTKVDDNRYSMKIFFEIRKQKYLEALDKRDRSKGVEILVKD  100
NtTPR-1    DETENYLSSFTGVSDNKFSTKMFFEIRKQKFLEALDRQDRKTALDILLKD   99
NtTPR-2    DEVERYLSGFTKVEDNRYSMKIFFEIRKQKYLEALDKNDRVKAVEILVKD  100
NtTPR-3    DEVEKYLSGFTKVDDNRYSMKIFFEIRKQKYLEALDGQDKAKAVEILVND  100
NtTPR-4    DEVEKYLSGFTKVDDNRYSMKIFFEIRKQKYLEALDRNDRPKAVDILIKD  100
           **.*.***.** *.**::* *:********:***** .*: ..::**::*

NtTPL      LKVFASFNEDLFKEITQLLTLENFRENEQLSKYGDTKSARAIMLVELKKL  150
NtTPR-1    LKVFAPSNEELYKEMAQLLTLDDFREYAPLAQYGDIISARKLMMRELKLI  149
NtTPR-2    LKVFASFNEDLFKEITQLLTLDNFRQNEQLSKYGDTKSARNIMLVELKKL  150
NtTPR-3    LKVFSTFNEDLYKEITQLLTLTNFRENEQLSKYGDTKTARSIMLIELKKL  150
NtTPR-4    LKVFSAFNEDLFKEITQLLTLDNFRDNEQLSKYGDTKSARSIMLLELKKL  150
           ****:.  **:*:**::***** :**:    *::***   :** :*: *** :

NtTPL      IEANPLFRDKLQFPNLKNSRLRTLINQSLNWQHQLCKNPRPNPDIKTLFV  200
NtTPR-1    IESNPQFKGRLRFPELNKSRLRRLINQSLNWQHMHCANPQLEPEIETLFT  199
NtTPR-2    IEANPLFRDKLTFPSFKASRLRTLINQSLNWQHQLCKNPRPNPDIKTLFT  200
NtTPR-3    IEANPLFREKLVFPTLRSSRLRTLINQSLNWQHQLCKNPRPNPDIKTLFT  200
NtTPR-4    IEANPLFRDKLNFPTLKNARLRTLINQSLNWQHQLCKSPKPNPDIKTLFV  200
           **:** *: :* ** :. :*** **********  * .*: :*:*:***.

NtTPL      PS-MDYPSGESDHVAKRTR-SLGISDEVN-LPVNVLPISFPGQGHNQALS  337
NtTPR-1    --------ATTSKAVQDSGNLYDVSSTRN-MNEKVLATGVTCQKQAVSSS  289
NtTPR-2    PG-MDYQMADSEHLMKRMR-A-GQSDEV----------SFSGSTHPPHMY  329
NtTPR-3    LGMLDYQNADHEQLMKRLR-PAPSVEEV----------TYP-TVRQQASW  332
NtTPR-4    PT-VDYQTADSEHMLKRSR-PFGVSDEVNNMPINILPGGYSGQSHAQSSY  343
                     .  .:  :          .               .    :

NtTPL      GGLPASPRIRFNKDGSLLAVSANENGIKILANSDGIRLVRTFENLAYDAS  686
NtTPR-1    GDLPASPYIRFNKKGTLLAVFADHNRIKILANDCGRFLLQTS----SDAS  635
NtTPR-2    GGLPASPRLRFNKEGSLLALTTSDNGIKVLANTDGQQMLRMLESRAFEGS  678
NtTPR-3    GGLPSLPRLRFNKEGNLLAVTTADNGIKILGNAAGMRSLRSVEASPFEAL  680
NtTPR-4    GGIPASPCLRFNKEGILLAVSTSDNGIKILANADGLRLLRSMENRQFDAS  692
           *.:*: * :****.* ***: : .* **:*.*  *   ::        :.

NtTPL      AAHPQDPNQFALGLSDGSVHVFEPLESEGKWGVPPPLENGSANGMPTAP- 1115
NtTPR-1    AAHPQNPNQFALGLTDGGVVIIEPPESEGRWLVHKGDDNNDHT------- 1050
NtTPR-2    AAHPSDSNQFALGMSDGTVHVIEPSDAEPKWGSSPPQDNGSMPSIPSSS- 1124
NtTPR-3    AAHPQEPSQFAVGLSDGTVKVIEPLESEGKWGLSPPVDNGILNGRTASSS 1120
NtTPR-4    AAHPSDPNQFALGLSDGAVIVLEPLESEGKWGTLPPAENGAGPS------ 1124
           ****.:..***:*::** * ::** ::* :*      :*.

NtTPL      SVGASGSEQAPR 1127
NtTPR-1    -----------K 1051
NtTPR-2    ALNCQPSETPSR 1136
NtTPR-3    TTSNHVADQVQR 1132
NtTPR-4    TSGAANSDQPQR 1136
                     :
```

**Fig. S21** Peptides of TOPLESS-RELATED3 identified by NIMIN1–TurboID mass spectrometry in *Nicotiana benthamiana*. Multiple sequence alignment of tobacco TOPLESS (TPL) and TOPLESS-RELATED (TPR) proteins. Dashes indicate gaps introduced to maximize alignments, and identical amino acids are marked by asterisks. The amino acid positions are given on the right. Only protein regions with identified peptides are displayed. Peptides identified in the same MS run as peptides of the NIMIN1 bait shown in Figure S20 are highlighted. Peptides marked in green are common to more than one member of the TPL family, while peptides marked in yellow and orange are unique to one family member. The protein coverage of TPR3 is 10%.

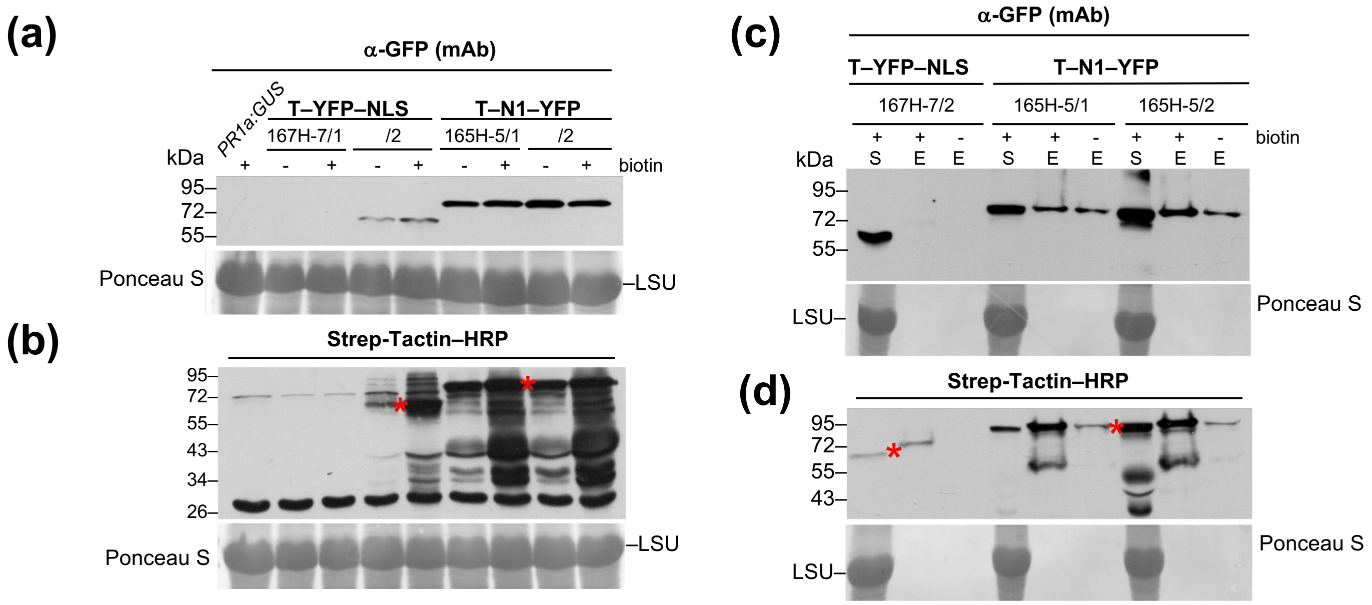


**Fig. S22** Proximity-dependent labeling of *Nicotiana tabacum* proteins by NIMIN1–TurboID. (a) Overexpression of TurboID fusion genes in *N. tabacum*. Accumulation of Turbo–YFP–NLS (T–YFP–NLS) and Turbo–NIMIN1–YFP (T–N1–YFP) fusion proteins in two T1 plants each from transgenic lines 167H-7 (*UBQ10$_{Pro}$:T–YFP–NLS*) and 165H-5 (*UBQ10$_{Pro}$:T–N1–YFP*). A transgenic plant harboring the *PR1a$_{Pro}$:GUS* reporter (Grüner & Pfitzner, 1994) served as negative control. Leaf disks were treated with water (-) or biotin (+) as indicated. Immunodetection in leaf extracts was with a monoclonal antibody (mAb) directed against GFP. Staining of the large subunit of RUBISCO (LSU) with Ponceau S demonstrates loading of the nitrocellulose filter. (b) Detection of biotinylated proteins in *N. tabacum* expressing TurboID constructs. Crude protein extracts were separated on SDS gels. Loading of the gel was in the same order as displayed in (a). Biotinylated proteins were detected by Strep-Tactin–HRP blotting. Red asterisks denote ligase self-biotinylation of the respective fusion protein. (c) TurboID pull-down. Biotinylated proteins were purified from crude extracts of *N. tabacum* plants expressing TurboID constructs with Strep-Tactin-coated beads. Aliquots from the supernatant (S) and the eluates from biotin-treated (E+) and water-treated leaf disks (E-) were separated on SDS gels, and ligase fusion proteins were detected by anti-GFP(mAb) blotting. (d) TurboID pull-down. Loading of the gel was in the same order as displayed in (c). Biotinylated proteins were detected by Strep-Tactin–HRP blotting. Red asterisks denote ligase self-biotinylation of the respective fusion protein.

```
NIM1-like1      GQEANKDRVCIDVLEREMRRNPMAGDALFSSPMLADDLHMKLHYLENRVA 437
NPR1            GKSASKDRLCIEILEQAERRDPLLGEASVSLAMAGDDLRMKLLYLENRVG 433
                *:.*.***:**::**:  **:*: *:* .* .* .***:*** ******.

NIM1-like1      FARLLFPLEARLAMQIANAETAAEVAVR-LASKSTSGNLREVDLNETPIK 486
NPR1            LAKLLFPMEAKVAMDIAQVDGTSEFPLASIGKKMANAQRTTVDLNEAPFK 483
                :*:****:**::**:**:.: ::*..:  :..* :..:   *****:*:*

NIM1-like1      QKERLLSRMQALSKTVELGKRYFPHCSQVLDKFME-DDLPDLIFLEMGTP 535
NPR1            IKEEHLNRLRALSRTVELGKRFFPRCSEVLNKIMDADDLSEIAYMGNDTA 533
                 **. *.*::***:*******:**:**:**:*:*: ***.:: ::  .*.

NIM1-like1      EEQKIKRKRFKELKDDVQRAFNKDKAELHCSRLSSSSCSSSFKDG---AS 582
NPR1            EERQLKKQRYMELQEILSKAFTEDKEEFDKTNHLSSSCSSTSKGVDKPNK 583
                **:::*::*: **:: :.:**.:** *:. :.  ******: *.     .

NIM1-like1      VKLRKL 588
NPR1            LPFRK- 588
                : :**
```

**Fig. S23** Peptides of NIM1-like1 (NPR3-like) and NPR1 identified by NIMIN1–TurboID mass spectrometry in transgenic *Nicotiana tabacum* plants. Sequence alignment of tobacco NIM1-like1 (NPR3-like) and NPR1. Dashes indicate gaps introduced to maximize alignments, and identical amino acids are marked by asterisks. The amino acid positions are given on the right. Only the C-terminal regions with identified peptides highlighted in yellow are displayed.

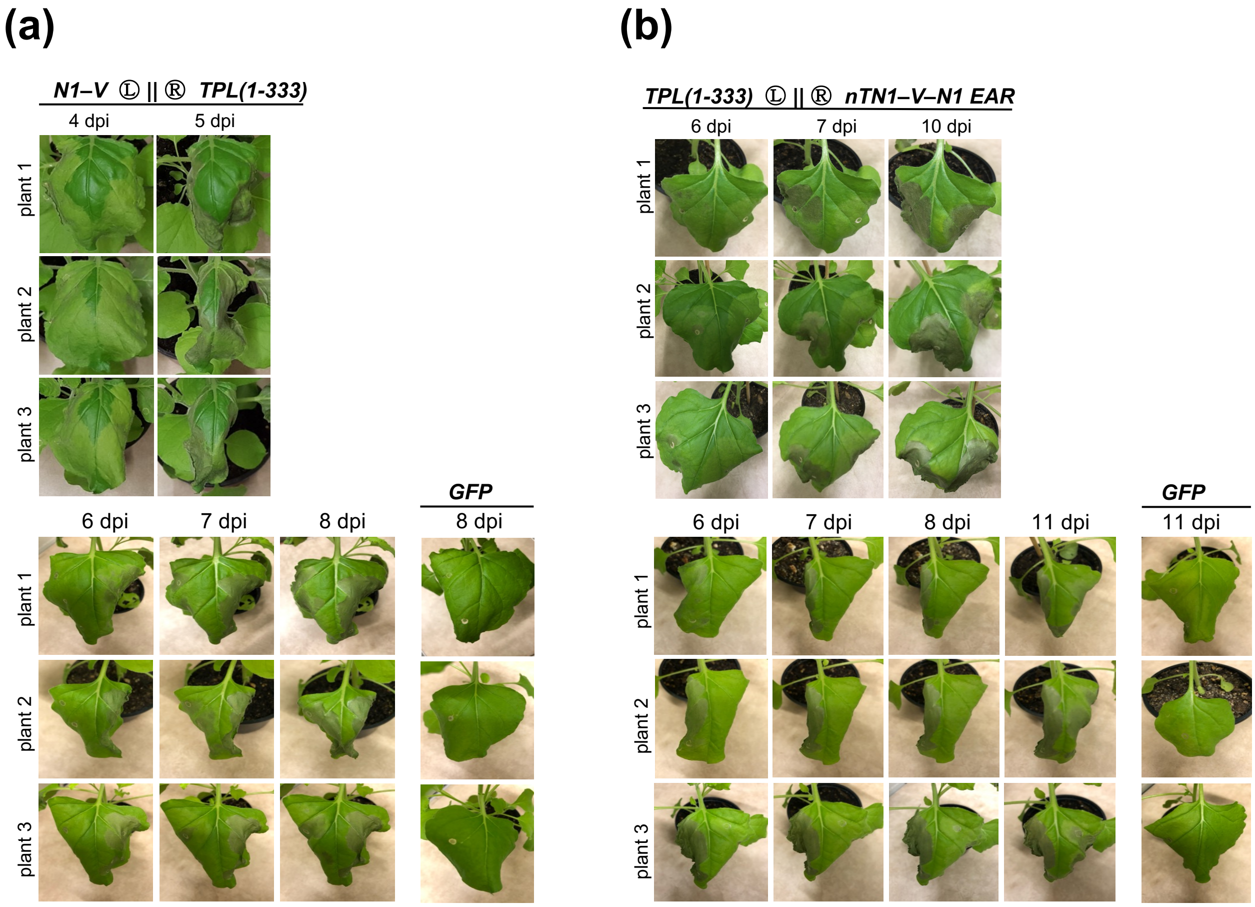


**Fig. S24** The N-terminal TOPLESS fragment, TPL(1-333), induces cell death in *Nicotiana benthamiana*. (a,b) Phenotype of *Nicotiana benthamiana* leaves overexpressing *TPL(1-333)*. Agrobacteria were infiltrated in the right (a) and left (b) halves of leaves as indicated. Effects of *TPL(1-333)* overexpression are compared to overexpression of *NIMIN1–VENUS* (a) and to overexpression of *nTN1–VENUS–N1EAR* (b).

**Table S1** Domain structures of Arabidopsis NIMIN proteins and mutants used in this work.

**NIMIN1**

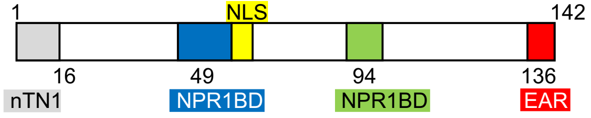


| Mutant | Kind of mutant | Domain(s) affected |
|---|---|---|
| N1ΔnT<br>N1(16-142) | single domain mutant | lacking N-terminal region |
| N1 F49/50S | single domain mutant | substitutions in blue NPR1BD |
| N1 E94A/D95V | single domain mutant | substitutions in green NPR1BD (EDF motif) |
| N1 L138/140A | single domain mutant | substitutions in EAR motif |
| N1ΔEAR<br>N1(1-137) | single domain mutant | lacking EAR motif |
| N1m2<br>N1 F49/50S E94A/D95V | double domain mutant | substitutions in both NPR1BDs |
| N1m3<br>N1 F49/50S E94A/D95V ΔEAR | triple domain mutant | substitutions in both NPR1BDs and lacking EAR motif |

**NIMIN1b**

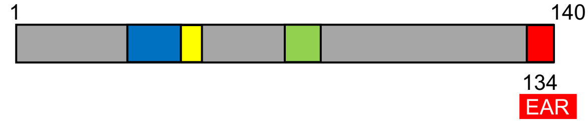


| Mutant | Kind of mutant | Domain affected |
|---|---|---|
| N1bΔEAR<br>N1b(1-135) | single domain mutant | lacking EAR motif |

**NIMIN3**

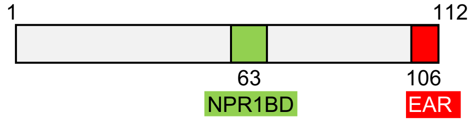


| Mutant | Kind of mutant | Domain(s) affected |
|---|---|---|
| N3 E63A/D64V | single domain mutant | substitutions in green NPR1BD (EDF motif) |
| N3 L108/110A | single domain mutant | substitutions in EAR motif |
| N3ΔEAR<br>N3(1-107) | single domain mutant | lacking EAR motif |

**Table S2** Proteins identified by mass spectrometry with NIMIN1–TurboID and TurboID–NLS in agroinfiltrated *Nicotiana benthamiana* plants displaying an approximately even distribution in all samples.

| Protein | T–YFP–NLS | T–N1–YFP plant 1 | T–YFP–NLS | T–N1–YFP technical replicate plant 1 | T–N1–YFP biological replicate plant 2 | Ratio* |
|---|---|---|---|---|---|---|
| | (# unique peptides) | | | | | |
| cluster of RUBISCO LSU NbE05064989 LOC109220728 | 23 | 24 | 50 | 41 | 38 | 0.94 |
| cluster of RUBISCO SSU S41 NbE03058344 XP_016461746 | 8 | 8 | 13 | 8 | 7 | 0.73 |
| cluster of hsp70-Hsp90 organizing protein 2-like NbD044252 XP_009595716 | 17 | 23 | 16 | 18 | 18 | 1.19 |
| cluster of GAPDH B NbD022568 XP_016441750 | 17 | 9 | 19 | 12 | 10 | 0.57 |
| cluster of HSP70 NbD020434 XP_016478491 | 23 | 12 | 14 | 8 | 10 | 0.54 |
| cluster of ATP synthase CF1 alpha NbD029578 NP_054481 | 10 | 9 | 12 | 7 | 7 | 0.70 |
| cluster of elongation factor 1-alpha-like NbD042613 XP_019260935 | 11 | 9 | 11 | 9 | 7 | 0.76 |
| cluster of elongation factor TuB NbD027633 XP_016502071 | 11 | 7 | 13 | 8 | 7 | 0.61 |
| cluster of PEP carboxylase-like NbE44072113 XP_016507808 | 11 | 9 | 10 | 6 | 8 | 0.73 |

*The ratio is the quotient between the average number of peptides identified in the technical and biological replicate Turbo–N1–YFP samples and the number of peptides identified in the Turbo–YFP–NLS samples.

**Table S3** Proteins identified by mass spectrometry with NIMIN1–TurboID and TurboID–NLS in transgenic *Nicotiana tabacum* plants displaying an approximately even distribution in all samples.

| **Protein** | **T–YFP–NLS** 167H-7/2 | **T–N1–YFP** 165H-5/1 | **T–N1–YFP** biological replicate 165H-5/2 | **T–N1–YFP** technical replicate 165H-5/2 | **Ratio*** |
|---|---|---|---|---|---|
| | (# unique peptides) | | | | |
| cluster of RUBISCO LSU Nitab4.5_0006084g0010 | 12 | 15 | 15 | 15 | 1.25 |
| RUBISCO SSU Nitab4.5_0000650g0110 | 2 | 4 | 3 | 2 | 1.50 |
| GAPDH type 1 Nitab4.5_0004979g0020 | 6 | 4 | 6 | 4 | 0.78 |
| cluster of HSP70 Nitab4.5_0004078g0050 | 4 | 5 | 5 | 7 | 1.42 |
| cluster of translation elongation factor EFtu/EF1A Nitab4.5_0000003g0860 | 2 | 5 | 4 | 5 | 2.33 |
| cluster of tetratricopeptide-like helical Nitab4.5_0003983g0040 | 38 | 39 | 41 | 32 | 0.98 |
| cluster of pre-ATP-grasp domain, carbamoyl-phosphate synthase Nitab4.5_0000469g0180 | 35 | 38 | 45 | 43 | 1.20 |
| cluster of biotin/lipoyl attachment, biotin-binding site Nitab4.5_0000361g0060 | 8 | 6 | 9 | 6 | 0.88 |
| cluster of ATPase, AAA-type Nitab4.5_0000722g0100 | 6 | 9 | 9 | 10 | 1.56 |

*The ratio is the quotient between the average number of peptides identified in the Turbo–N1–YFP samples and the number of peptides identified in the Turbo–YFP–NLS sample.

**Table S4** Primers used for gene construction.

| Construct | Primer name | Sequence 5´ to 3´ |
|---|---|---|
| ***NIMIN1*** | | |
| forward primer | N1fwd | CGGGATCCATATGTATCCTAAACAATTTAG |
| back primer | N1-4 | TTGGATCCCAATGCAAGATTAAGATC |
| *N1ΔnT* *N1(16-142)* | N1-10 | TTGGATCCATATGAGCAAGGATGAGAATGTGG |
| *N1 F49/50S* | N1-M1 | GGAAGCTTAGAGGACGTATCAATCTTC |
| *N1 E94A/D95V* | N1-6 | CACTGAGAGAAAACCGCCGGCTGAAACGC |
| | N1-5 | GCGTTTCAGCCGGCGGTTTTCTCTCAGTG |
| *N1 L138/140A* | N1-7 | TTGGATCCCAATGCAGCATTAGCATCTAAAGCCTTGTC |
| *N1ΔEAR* *N1(1-137)* | N1-13 | AAGGATCCTAAAGCCTTGTCTTCGTTTCGC |
| *N1NcoI* | 5N1NcoI | ACCATGGCAACTAGTATGTATCCTAAACAATTTAGTTTA |
| | 3N1NcoI | ACCATGGAACTAGTCAATGCAAGATTAAGATCTAA |
| ***NIMIN1b*** | | |
| forward primer | N1b-1 | GGGGATCCATATGAACCAAGAAGAAG |
| back primer | N1b-2 | CCGGATCCCAATGCAAGATTAAGATCTAAACC |
| back primer | N1b stop | CCGGATCCGAGCTCTTACAATGCAAGATTAAGATCTAAACC |
| *N1bΔEAR* *N1b(1-135)* | N1b-6 | CCGGATCCTAAACCATTATCTTTTCTCATCACC |
| *nTN1–N1b* | N1b-4 | GGGGATCCATATGTATCCTAAACAATTTAGTTTATACAATTATTCCCTAGAGACCATGAAC CAAGAAGAAGAAAAAACAGAG |
| *N1b–nTN1* | N1b-7 | CCGGATCCGGTCTCTAGGGAATAATTGTATAAACTAAATTGTTTAGGATACATCAATGCAA GATTAAGATCTAAACCATTATCTTTTCTC |
| **NIMIN2** | | |
| forward primer | N2fwd | GGGGATCCATATGAACAACTCTTTGAAG |
| back primer | N2-3 | GGGGATCCCAACGATAAACTAACGCTGTCTGG |
| ***NIMIN3*** | | |
| forward primer | N3fwd | GGGGATCCATATGGACAGAGACAGAAAGAG |
| back primer | N3-1 | TTGGATCCCAGAGAAAGATTCAAGTC |
| *N3 E63A/D64V* | N3-3 | CATGAAAATGAAAACTGCTGGCTGAAACG |
| | N3-2 | CGTTTCAGCCAGCAGTTTTCATTTTCATG |
| *N3 L108/110A* | N3-4 | TTGGATCCCAGAGAAGCATTCGCGTCTAAACAAACGTTAGTCTC |
| *N3ΔEAR* *N3(1-107)* | N3-5 | CCAGATCTGTCTAAACAAACGTTAGTCTCAGATCC |
| **Nt*NIMIN2c*** | | |
| forward primer | AD-10/2 | CCGGATCCATATGCTACTTACTATGGACG |
| back primer | AD-10/1 | AAGGATCCGTCTCCGCCTTCTGGTAAAGC |
| back primer | N2c-5 | AAGAGCTCTTAGTCTCCGCCTTCTGG |
| Nt*N2cΔEAR* Nt*N2c(1-111)* | N2c-9 | AAGGATCCCAAACCACCTTTTCGCACG |
| **Nt*NIMIN2d*** | | |
| forward primer | FS-1 | TTGGATCCATATGCCGCTAATGGAGGGTG |
| back primer | FS-2 | AAGGATCCAACGCCGTTAGTCTCTGG |
| ***AtNPR1 BamHI*** | | |
| forward primer | NPR1-8 | TTGGATCCATATGGACAAACGAGAACAAATTCC |
| back primer | NPR1-12B | TTGGATCCCCGACGACGATGAGAGAG |
| ***VENUS*** | | |
| forward primer | Venus-5 | TTGGATCCATGGTGAGCAAGGGCGAGGAGC |
| back primer | Venus-6 | TTGTCGACTTACTTGTACAGCTCG |
| *nTN1–VENUS* | Venus-7 | GGGGATCCATATGTATCCTAAACAATTTAGTTTATACAATTATTCCCTAGAGACATGGTGA GCAAGGGCGAGGAGC |
| back primer | Venus-4 | GGGAGCTCTTACTTGTACAGCTCG |
| *VENUS–nTN1* | Venus-8 | GGGAGCTCTTAGGTCTCTAGGGAATAATTGTATAAACTAAATTGTTTAGGATACATCTTGT ACAGCTCGTCCATGCCGAGAG |
| *nTN1–VENUS–NLS* | Venus-12 | AGAGCTCGTCGACCTATCGTCTACGTTTCCGCTTGTACAGCTCGTCCATGCCG |
| *nTN1–VENUS–NPR1BD* | Venus-16 | AGAGCTCGTCGACCTATGCTTCTTGATAGTGTTTGATAAGCTTAAAGAACGTATCAATCTT CTTCTCTCGTCTACGTTTCCGCTTGTACAGCTCGTCCATGCCG |
| *nTN1–VENUS–N1EAR* | Venus-9 | AGAGCTCGTCGACCTACAATGCAAGATTAAGATCTAAAGCCTTGTCTTCGTTTCGTCTACG TTTCCGCTTGTACAGCTCGTCCATGCCG |
| *nTN1–VENUS–N2dEAR* | Venus-15 | AGAGCTCGTCGACTTAAACGCCGTTAGTCTCTGGTTCTGGCTCCATGTTCAGGTCCAATTC CAAACCCCTCTTCCTCTTCTTCTTGTACAGCTCGTCCATGCCG |
| ***TOPLESS*** | | |
| forward primer | TPL-1 | AAGTCGACAGATCTATATGTCTTCTCTTAGTAGAGAGCTCG |
| back primer | TPL-2 | TTCTGCAGTCAAGATCTCTGAGGCTGATCAGATGCAGAGGC |
| *TPL(1-333)* | TPL-4 | TTCTGCAGCCATGGGCCTGCCCTGAAAATGAC |
| *TPL(1002-1131)* | TPL-3 | TTGTCGACAATGCATGAAACAGTGGGCTGTTCG |

**Methods S1** Detailed description of methods.

### DNA constructs for analysis in yeast

For protein-protein interaction assays in yeast, cDNA sequences were fused in-frame to the sequence for Gal4 DNA-binding domain (GBD) in pGBT9 and the sequence for Gal4 transcription activating domain (GAD) in pGAD424. Plasmids encoding AtNPR1, NtNPR1, NIMIN1, N2, N3 and NtN2a fusion proteins were described previously (Weigel *et al.*, 2001; Maier *et al.*, 2011). Generally, clones coding for full-length, truncated or chimeric NIMIN proteins were generated by PCR amplification, and clones encoding mutant NIMIN proteins (single domain mutants) were generated by overlap extension PCR (Ho *et al.*, 1989). The sequence for the double domain mutant N1 F49/50S E94A/D95V (N1m2) was assembled from the sequences of the respective single domain mutants by exploiting an internal *Hin*dIII restriction enzyme site. Primers used are listed in Table S4. All *NIMIN* sequences were cloned as *Bam*HI fragments in pGBT9. Mutant *NIMIN* genes are depicted in Table S1.

To generate the *NtNPR1–GUS* fusion gene, the sequence encoding GUS was excised from pBI101.1 (Jefferson *et al.*, 1987), inserted in *Bam*HI/*Pst*I cleaved pGAD424, and the *NtNPR1 Bam*HI fragment was ligated to this plasmid. The sequence for VENUS was amplified by PCR with primers Venus-5 and Venus-6, and the *NIMIN1* sequence was added to the 5′-end as *Bam*HI fragment. Constructs with single NIMIN domains added to the N-terminus or the C-terminus of VENUS were generated by PCR amplification with primers listed in Table S4. Fragments were inserted in pGBT9 digested with *Bam*HI and *Sal*I.

The sequence encoding TOPLESS(1-333) was amplified from pDEST-AD/TPL (Arabidopsis Biological Resource Center pDEST-AD012F12) and ligated as *Sal*I/*Pst*I fragment to pGAD424. From this, the plasmid encoding TPL(1-194) was generated by subcloning the 0.6 kb *Bam*HI/*Eco*RV fragment encompassing the TPL N-terminus in pGAD424. The TPL full-length aa sequence was assembled by adding PCR-amplified *TPL(1002-1131)* and an internal 2 kb *Nco*I/*Nsi*I fragment isolated from pDEST-AD/TPL to pUC19/TPL(1-333).

All clones generated by PCR amplification were verified by DNA sequence analysis.

### DNA constructs for analysis in plants

For expression studies in plants, constructs were inserted in derivatives of pBin19 (Bevan, 1984). Plasmids harboring *GFP, VENUS, AtNPR1, NtNPR1, N1, N2* and *N3* cDNAs under control of the *CaMV 35S* promoter and the plasmid harboring *N1$_{Pro}$:6xHis–Bax* were described previously (Haseloff *et al.*, 1997; Glocova *et al.*, 2005; Maier *et al.*, 2011; Hermann *et al.*, 2013; Stos-Zweifel *et al.*, 2018; Neeley *et al.*, 2019). Clones coding for NIMIN1b and NtN2c were generated by PCR amplification with forward and back primers (N1b stop and N2c-5, respectively) as given in Table S4, and PCR fragments were ligated to *Bam*HI/*Sac*I cleaved pBin19/35S:GUS (Jefferson *et al.*, 1987) from which the *GUS* reporter was removed. To express *NIMIN–VENUS* fusion genes, all *NIMIN* sequences were amplified by PCR using primers listed in Table S4, cloned in pGBT9 and added as *Bam*HI fragments to pBin19/35S:VENUS. *VENUS* fusions with sequences encoding isolated NIMIN domains were amplified by PCR with forward primer Venus-7 along with a specific back primer or Venus-4 (Table S4), and products were inserted as *Bam*HI/*Sac*I restriction fragments into pBin19/35S:GUS after excision of the *GUS* gene.

The *AtNPR1–GUS*, *NtNPR1–GUS* and *AtNPR1–GFP* fusion genes were generated by in-frame insertion of the respective *NPR1 Bam*HI fragment in pBin19/35S:GUS (pBI101.1) and pBin19/35S:mGFP4. The sequence encoding TPL(1-333) was excised from pGAD424 and ligated as *Bgl*II fragment to pBin19/35S:NPR1 from which *NPR1* was deleted by cleavage with *Bam*HI.

For proximity labeling by TurboID, *NIMIN1* was amplified with primers 5N1NcoI and 3N1NcoI and ligated to R4pGWB601_UBQ10p-Turbo-NES-YFP cut with *Nco*I. As a control, R4pGWB601_UBQ10p-Turbo-YFP-NLS was used directing the Turbo–YFP fusion protein to the nucleus (Mair *et al.*, 2019).

All clones generated by PCR amplification were verified by DNA sequence analysis.

### Transient gene expression

Transient gene expression by agroinfiltration was performed as described previously (Hermann *et al.*, 2013). Briefly, gene constructs in pBin19 and TurboID vectors were transferred to *Agrobacterium tumefaciens* strain LBA4404 by triparental mating. Bacterial strains were adjusted to an optical density (OD600) of 0.5 in 10 mM $MgCl_2$ supplemented with 150 µM acetosyringone. Each bacterial suspension was mixed with an equal volume (1+1) of a strain carrying the p19 suppressor from *Tomato bushy stunt virus*, and bacteria were infiltrated in leaves of three different *Nicotiana benthamiana* plants. To monitor effects of *NIMIN* overexpression on *PR-1* gene induction, reporter plants with an integrated *PR-1a$_{Pro}$:GUS* construct were agroinfiltrated (Hermann *et al.*, 2013). In co-infiltration experiments, the bacterial suspension was mixed with equal volumes of a strain expressing a second gene or with Agrobacteria without gene construct and two volumes of the p19 strain (1+1+2). To allow direct comparison of phenotypic effects and protein accumulation from two different gene constructs, Agrobacterium suspensions were infiltrated in the left (L) and right (R) halves of the same leaves, respectively. Plants infiltrated with a strain harboring *35S:mGFP4* served as positive controls for gene expression levels and immunodetection with GFP antibodies and as negative controls for phenotypic observations, DAB staining and GUS enzyme assays.

Plants were inspected daily for visible effects induced by gene expression, and leaves were photographed at indicated times. Accumulation of GFP protein and accumulation of phenolic compounds were monitored under UV light. To determine the subcellular localization of YFP fusion proteins, epidermal peels were viewed under epifluorescence and bright field conditions 4 to 7 days post-infiltration (dpi) as reported by Maier *et al.* (2011). Proteins were extracted with GUS lysis buffer from two leaf disks for each plant 4 to 6 dpi when strong GFP fluorescence was observed. In co-expression experiments, leaf disks were sampled from the left and the right halves of the same leaves.

To determine microbial growth in agroinfiltrated tissue, leaf disks were harvested from different leaves immediately after infiltration (0 dpi) and at indicated times and ground in 10 mM $MgCl_2$. Serial dilutions were plated on LB medium without or with antibiotic as indicated. Plates were incubated at 30°C. Bacterial counts are given as colony-forming units (cfu) per leaf disk. Results are depicted as mean values from three replicates, plus and minus standard deviation (SD).

**GUS enzyme activity**

Determination of GUS enzyme activity was performed as described (Weigel *et al.*, 2001; Glocova *et al.*, 2005). GUS activities were measured directly in crude extracts from leaf disks cut from agroinfiltrated tissue or from leaf disks floated on water or chemicals. GUS activities are depicted as mean activities, plus and minus SD. Representative results obtained in one experimental set are shown. GUS activities are given in units (1 unit = 1 nMol 4-MU formed per hour per mg protein).

To demonstrate expression of *NPR1–GUS* fusion genes, an in-gel GUS activity stain was performed. Crude protein extracts from *N. benthamiana* leaves infiltrated with

Agrobacteria harboring *35S:GUS* or *35S:(Nt)NPR1–GUS* constructs were incubated for 15 min at room temperature with SDS gel loading buffer, and samples were separated on a 7.5% SDS polyacrylamide gel. After electrophoresis, the gel was equilibrated for 2 h at 4°C in GUS lysis buffer and then incubated in 1 mM 4-methylumbelliferyl-β-D-glucuronide (MUG) in lysis buffer at 37°C. GUS enzyme activity was visualized as fluorescent band on a UV transilluminator.

**TurboID-based proximity labeling**

Disks were cut 4 or 5 dpi from leaf areas infiltrated with Agrobacteria or from leaves of young transgenic tobacco plants harboring TurboID gene constructs. Leaf disks were briefly vacuum-infiltrated with water or 50 µM biotin and then floated for 1 hour at room temperature. Proteins were extracted with GUS lysis buffer from at least 4 thoroughly dried disks per sample. For enrichment of biotinylated proteins, crude extracts were incubated under inversion with washed Strep-Tactin Superflow resin for 1 h at room temperature or overnight at 4°C. Strep-Tactin-loaded beads were washed three times with buffer (100 mM Tris-HCl, 150 mM NaCl, 1 mM EDTA, pH 8, 0.4% SDS) and eluted by boiling in SDS loading buffer. Fusion proteins, biotinylated proteins and endogenous proteins were detected by blotting and silver staining of SDS polyacrylamide gels.

**Staining and detection of proteins**

Proteins were extracted with GUS lysis buffer from leaf disks cut from agroinfiltrated tissue or from transgenic plants and from dried disks after floating on water or chemicals. Crude extracts were cleared twice by centrifugation, equal extract volumes were loaded on SDS polyacrylamide gels, and separated proteins were blotted to nitrocellulose filters. For immunodetection, membranes were incubated with polyclonal antisera raised against 6xHis–AtNPR1 (Stos-Zweifel *et al.,* 2018) and GFP (Santa Cruz Biotechnology sc-8334) or with a monoclonal antibody (mAb) directed against GFP (ThermoFisher Scientific G10362) as indicated. Immunodetection of GFP with the monoclonal antibody was considerably less sensitive than detection with the polyclonal antiserum. The same extracts used for immunodetection were also used for measuring GUS reporter enzyme activities. Biotinylated proteins were detected with a Strep-Tactin horse-radish peroxidase (HRP) conjugate.

Protein loading was checked by staining nitrocellulose filters with Ponceau S (0.1% in 5% acetic acid). For silver staining, gels were fixed for at least one hour in 50% ethanol/12%

acetic acid and rinsed three times in 50% ethanol. Then, gels were soaked exactly for 1 minute in 0.02% sodium thiosulfate pentahydrate and rinsed three times for 20 seconds each in water. Gels were transferred to silver nitrate solution (0.2% silver nitrate, 0.075% formaldehyde (37%)) for 20 minutes, rinsed three times for 20 seconds each in water and then submerged in developer (6% sodium carbonate, 0.5 ml formaldehyde (37%) and 20 ml 0.02% sodium thiosulfate pentahydrate per liter). The reaction was stopped with 50% methanol/12% acetic acid.

For detection of NIMIN fusion proteins in yeast, extracts were prepared from transformed cells as reported, and nitrocellulose filters were incubated with a polyclonal antiserum directed against the Gal4 DNA-binding domain (Weigel *et al.*, 2001). Unspecific bands reacting with the antiserum demonstrate gel loading.

**LC-MS/MS sample preparation**

For protein identification, Strep-Tactin-loaded beads were washed three times with SDS buffer (100 mM Tris-HCl, 150 mM NaCl, 1 mM EDTA, pH 8, 0.4% SDS) and six times with wash buffer without SDS. Bound proteins were reduced with DTT (dithiothreitol) for 30 min at room temperature at a final concentration of 10 mM. Subsequently, samples were alkylated with 5 mM CAA (chloroacetamide) for 30 min in the dark. Next, samples were diluted to a final concentration of 2 M urea with 50 mM Tris-HCl pH 8.5 and digested directly on the Strep-Tactin-loaded beads by 0.75 µg trypsin (Roche) overnight at 25°C. TFA (trifluoroacetic acid) was added to a final concentration of 0.5% to the samples, released peptides were concentrated and desalted on C18 Stage Tips as described by Rappsilber *et al.* (2007) and dried under vacuum. Dried samples were dissolved in 0.1% TFA.

**Nano-LC-MS/MS analysis**

Nano-LC-ESI-MS/MS experiments were performed on an Ultimate 3000 nano-RSLC (Thermo Fisher Scientific) coupled to an Exploris 480 mass spectrometer (Thermo Fisher Scientific) using a Nanospray-Flex ion source (Thermo Fisher Scientific). Peptides were first loaded on a trap column (5 mm x 30 µm, Thermo Fisher Scientific) and separated on a 25 cm x 75 µm nanoEase MZ HSS T3 reversed phase column (100Å pore size, 1.8 µm particle size, Waters, USA) operated at constant temperature of 35°C. Peptides were separated at a flow rate of 300 nL/min using a gradient with the following profile: 2% - 55% solvent B in 30 min, 55% - 95% solvent B in 10 min, isocratic 95% solvent B for 5 min and 95% - 2% solvent B in 10 min.

Solvents used were 0.1% FA (solvent A) and 0.1% FA in ACN/H2O (80/20, v/v, solvent B). The Orbitrap Exploris 480 and the Ultimate 3000 were operated under the control of XCalibur software (version 4.4.16.14) and Sii Xcalibur (version 1.6.0.6893; Thermo Fisher Scientific Inc., USA.). MS spectra (m/z = 200-2000) were detected in the Orbitrap at a resolution of 60000 (m/z = 200). Maximum injection time (MIT) for MS spectra was set to 50 ms, the automatic gain control (AGC) value was set to 3 x 10E6. Internal calibration of the Orbitrap analyzer was performed using lock-mass ions from ambient air as described in Olsen *et al.* (2005). The MS was operating in the data dependent mode selecting the top 30 highest abundant peptide precursor signals for fragmentation (HCD, normalized collision energy of 30). For MS/MS analysis, only charge states from 2-6 were considered, the monoisotopic precursor selection was set to peptides and the minimum intensity threshold was set to 1 x 10E5. MS/MS scans were performed in the Orbitrap with a resolution of 150000, isolation width was set to 1.6 Da. The AGC target and max injection time for MS/MS scans was set to 5 × 10E4 and 50 ms respectively. Dynamic exclusion was set to 60 s with a tolerance of 10 ppm.

**MS data analysis**

Mascot 2.6 (Matrix Science, UK) was used as search engine for protein identification. Spectra were searched against the custom specific protein databases of *N. benthamiana* and *N. tabacum* proteins downloaded as FASTA-formatted sequences (Source Data Tables 1 and S2; Source Data Tables 2 and S3). Search parameters specified enzyme none, allowing no missed cleavages, a 5 ppm mass tolerance for peptide precursors and 0.02 Da tolerance for fragment ions. Methionine oxidation was allowed as variable modification and carbamidomethylation of cysteine residues was set as fixed modification. The Mascot results were transferred to ScaffoldTMSoftware 4.10.0 (Proteome Software, USA). Minimum probability cut-offs for peptide and protein identification, as specified by the Peptide Prophet algorithm (Keller *et al.*, 2002), are described in the Results section.

**Accession numbers**

Sequence data from this article can be found in the EMBL/GenBank databases under the following accession numbers:

At1g64280 (*NPR1*), AF480488 (*NtNPR1*), NW_015834793 (*NtNIM1-like1, NPR3-like*), At1g02450 (*NIMIN1*), At4g01895 (*NIMIN1b*), At3g25882 (*NIMIN2*), At1g09415 (*NIMIN3*),

EF015598 (*NtNIMIN2c*), FS401103 (*NtNIMIN2d)*, At1g15750 (*TOPLESS*), LOC107803083 (*NtTOPLESS*), LOC107818525 (*NtTOPLESS-RELATED1, TPR1*), LOC107793024 (*NtTPR2*), LOC107813542 (*NtTPR3*), LOC107812567 (*NtTPR4*) and X06930 (*NtPR-1a*).
